# Polyamines buffer labile iron to suppress ferroptosis

**DOI:** 10.1101/2025.06.30.662349

**Authors:** Pushkal Sharma, Heather R. Keys, Ryan P. Mansell, Jillian Stark, Louisa Girard, Christalyn Ausler, Rachel Anderson, Sebastian Müller, Shinya Imada, Ivan S. Pires, Tenzin Kunchok, Millenia Waite, Bingbing Yuan, Amy Deik, Luke Ferro, Paula T. Hammond, Raphaël Rodriguez, Maria-Eirini Pandelia, Whitney S. Henry, Ankur Jain

**Author notes:** Correspondence (A.J.), (W.S.H.).

## Abstract

Polyamines are essential and evolutionarily conserved metabolites present at millimolar concentrations in mammalian cells. Cells tightly regulate polyamine homeostasis through complex feedback mechanisms, yet the precise role necessitating this regulation remains unclear. Here, we show that polyamines contribute to endogenous buffering of redox-active iron, providing a molecular link between polyamine metabolism and ferroptosis. Using a genome-wide CRISPR screen, we identified a synthetic lethal dependency between polyamine depletion and the key ferroptosis suppressor, GPX4. Mechanistically, we show that polyamine deficiency triggers a redistribution of cellular iron, increasing the labile iron pool and upregulating ferritin. To directly visualize this iron buffering in living cells, we developed a genetically encoded fluorescent reporter for redox-active iron. Live-cell analysis revealed a striking inverse correlation between intracellular polyamine levels and redox-active iron at single-cell resolution. These findings reposition polyamines as key regulators of iron homeostasis, with implications for ferroptosis-linked disease states and cellular redox balance.

## Introduction

Polyamines are essential, evolutionarily conserved metabolites found in nearly all living organisms^1^. In mammalian cells, the major polyamines are putrescine, spermidine, and spermine (Figure 1A). These metabolites are present at millimolar concentrations and exist predominantly in protonated forms, enabling extensive interactions with negatively charged biomolecules, particularly RNA^2^. Despite their abundance, the molecular functions of polyamines remain incompletely understood.

**Figure 1:**
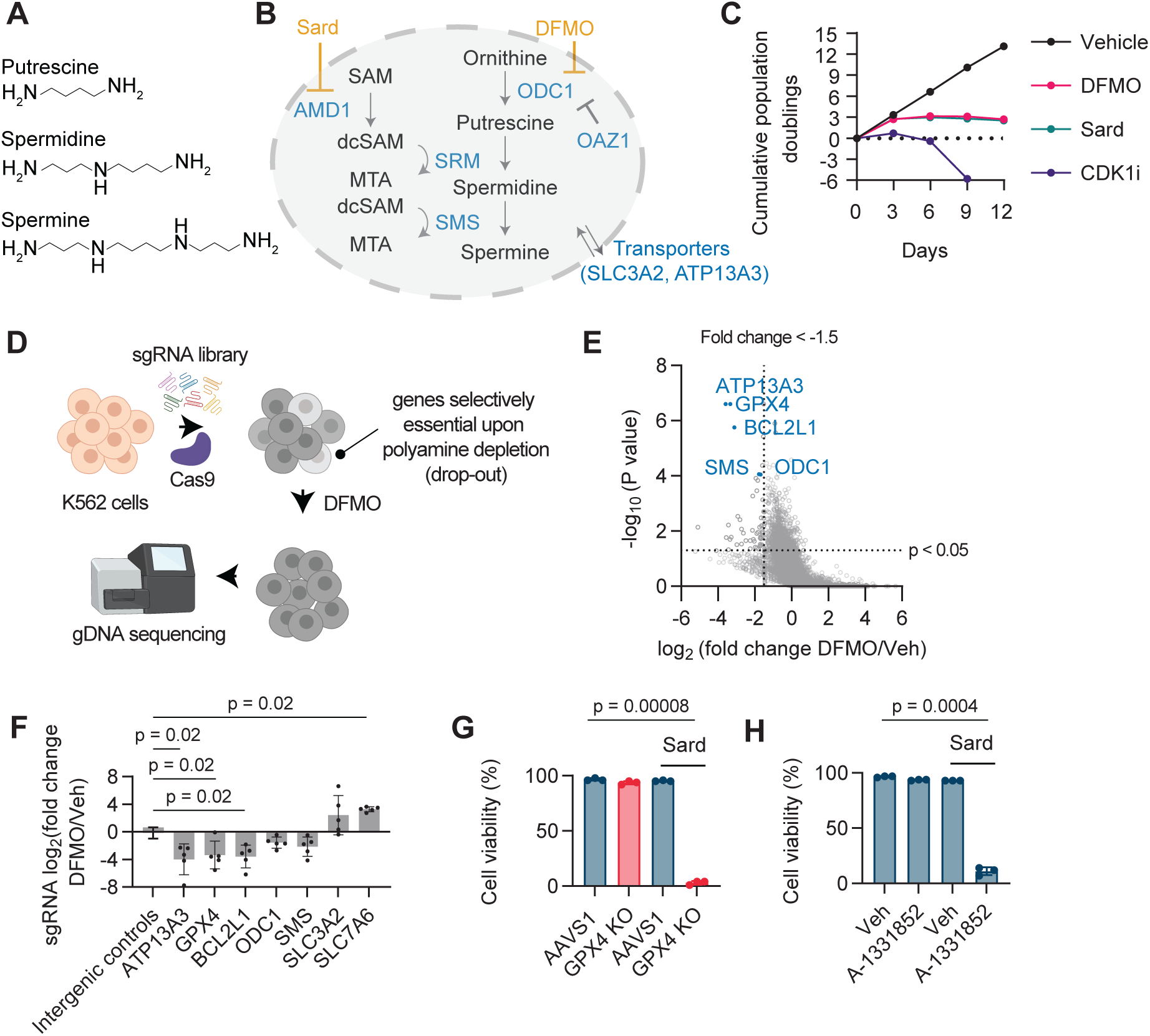
CRISPR screen identifies modulators of polyamine sensitivity. (A) Chemical structures of mammalian polyamines. (B) Schematic of the polyamine metabolic pathway. DFMO, difluoromethylornithine; Sard, sardomozide. (C) Cumulative population doublings of K562 cells treated with DFMO (0.5 mM), sardomozide (1 μM), or RO-3306 (5 μM, CDK1 inhibitor) over 12 days (passaged every 3 days; n = 1). (D) Schematic of the CRISPR knockout screen workflow. (E) Volcano plot of -log10(P-value) versus median log2(fold depletion) for genes in the CRISPR drop-out screen. Lines: p = 0.05, fold change = -1.5. Top hits are highlighted in blue. (F) Fold change of sgRNAs targeting indicated genes (mean ± SD, n = 5 sgRNAs/gene). (G) Viability of AAVS1 and GPX4 knockout (KO) K562 cells treated with sardomozide (5 μM, 96 h) (n = 3). GPX4 KO cells were propagated in liproxstatin-1 (1 μM), withdrawn 48 h before the assay. (H) Viability of K562 cells treated with A-1331852 (0.1 μM, 48 h) after sardomozide pre-treatment (5 μM, 72 h); DMSO as vehicle control (n = 3).

Cells tightly regulate polyamine homeostasis through an elaborate feedback network (Figure 1B). ODC1, the rate-limiting biosynthetic enzyme, has a short half-life and is uniquely regulated by polyamine-induced expression of antizyme (OAZ1) via programmed +1 ribosomal frameshifting^3,4^. Polyamine catabolism and transport are similarly under intricate feedback control, enabling cells to rapidly adjust intracellular concentrations in response to perturbations^5,6^. This raises a central question: why do cells maintain polyamines at millimolar concentrations and encode such extensive regulatory machinery? Their best-defined role is the hypusination of eIF5A, a spermidine-dependent modification which is essential for translation^7^. However, hypusination requires sub-micromolar substrate concentrations^8,9^ and does not explain the enormous metabolic investment in tightly regulating polyamine concentrations at millimolar levels. Polyamine dysregulation also affects diverse cellular processes including chromatin compaction^10^, cell differentiation^11^, tissue regeneration^12^, autophagy^13^, and immune function^14^. Polyamine levels decline with age, and dietary supplementation in model organisms improves memory^15^, cardiac function^16^, and lifespan^17^. Mutations in polyamine pathway genes cause disorders such as Parkinsonian syndromes^18^, Snyder-Robinson syndrome^19^ and Bachmann-Bupp syndrome^20^, and elevated levels of polyamines are observed across many cancers^21^. These pleiotropic effects suggest additional molecular functions that remain undefined.

To identify cellular dependencies that emerge during polyamine stress, we performed a genome-wide CRISPR-Cas9 screen^22,23^ in human cells under polyamine-depleted conditions. This synthetic lethal screen revealed a surprising link between polyamines and iron homeostasis: polyamine depletion rendered cells highly dependent on GPX4 (glutathione peroxidase 4), an antioxidant enzyme that protects membranes from lethal lipid peroxidation and ferroptotic cell death^24,25^. This dependency stems from an increase in labile iron pools upon polyamine depletion. Using Mössbauer spectroscopy and Fe^2+^-binding and competition assays, we show that polyamines can coordinate Fe^2+^. To visualize labile iron in living cells, we developed a genetically encoded fluorescent reporter. Single-cell analysis revealed that labile iron levels increase as polyamines decline. Together, these data support a model in which millimolar polyamines help restrain labile iron availability, linking polyamine metabolism to redox balance and ferroptosis sensitivity.

### CRISPR screen identifies genetic dependencies under polyamine depletion

The polyamine biosynthetic pathway involves two rate-limiting steps: the formation of putrescine by ODC1, and the generation of aminopropyl donors by AMD1, which are then incorporated into spermidine and spermine^1^. Polyamine depletion using the ODC1 inhibitor DFMO or the AMD1 inhibitor sardomozide induces cytostasis without causing substantial cell death (Figure 1C). To identify genetic dependencies under varying polyamine levels, we conducted a genome-wide CRISPR-Cas9 knockout screen in K562 cells (Figure 1D). Cells transduced with a CRISPR sgRNA library^26^ were cultured with or without DFMO for 14 days. Polyamine depletion was confirmed via enzymatic assay (Figure S1A), and sgRNA abundance was compared between treated and untreated conditions^27^ (Table S1).

Loss of genes related to polyamine biosynthesis, including ODC1 and SMS (spermine synthase), sensitized cells to DFMO treatment, validating our screening approach (Figure 1E-F). This analysis identified ATP13A3, BCL2L1, and GPX4 as the top synthetic lethal hits (Figure 1E-F). In contrast, knockout of the membrane proteins SLC3A2 and SLC7A6 enhanced survival under DFMO treatment (Figure 1F), likely reflecting their roles in polyamine export. SLC3A2 is an adapter protein that regulates several amino acid transporters and promotes polyamine excretion^28^. SLC7A6 forms a complex with SLC3A2^29^ and may similarly contribute to polyamine export. Although SLC3A2 also functions as the heavy-chain subunit of the cystine antiporter system Xc□, this is unlikely to explain the protective phenotype, since loss of system Xc□ activity would be expected to sensitize cells to ferroptosis^30^. Consistent with this, SLC7A11, the light-chain subunit of system Xc□, was not enriched (Table S1).

The synthetic lethality of ATP13A3, a known polyamine transporter^31^, likely reflects its role in the uptake of residual polyamines from the cell culture medium. Combined inhibition of polyamine synthesis (via DFMO) and transport (via ATP13A3 knockout) could fully deplete intracellular polyamines, inducing cell death. BCL2L1 and GPX4, two additional top hits, are key survival factors in distinct cell death pathways. BCL2L1 encodes an anti-apoptotic protein that binds the voltage-dependent anion channel (VDAC) and regulates mitochondrial membrane potential^32^. GPX4 is a lipid quality control enzyme that reduces phospholipid hydroperoxides to their corresponding alcohols, and is a central regulator of ferroptosis: an iron-dependent cell death pathway characterized by lipid peroxidation^24,25^.

We validated these findings using sardomozide, which targets AMD1 and blocks spermidine and spermine synthesis (Figure 1B). This mechanism is distinct from that of DFMO, which targets ODC1 and inhibits putrescine production. Sardomozide induced substantial cytotoxicity in GPX4 knockout cells but not in control cells targeted at the AAVS1 safe-harbor locus (Figure 1G). Because GPX4 loss alone is sufficient to trigger ferroptosis^24^, GPX4 knockout K562 clones were generated and propagated in the presence of the ferroptosis inhibitor liproxstatin-1, which was withdrawn 48 h before each assay (Methods). Under these conditions, GPX4 knockout cells proliferated more slowly than AAVS1-targeting controls but remained viable for the short-term assays (Fig. S1B-C). Pharmacological inhibition of BCL2L1 with A-1331852^33^, likewise sensitized cells to polyamine depletion, without reducing viability on its own (Figure 1H), confirming synthetic lethality. Given the central role of GPX4 in preventing ferroptosis, we focused on how polyamine depletion primes cells for this iron-dependent cell death.

### Polyamine depletion sensitizes cells to ferroptosis

Long-term maintenance of GPX4 knockout clones under ferroptosis suppression could select for compensatory GPX4-independent defenses^34^. We therefore tested whether polyamine depletion also sensitizes otherwise unadapted cells to acute GPX4 inhibition. Co-treatment with AMD1 inhibitor sardomozide and GPX4 inhibitors (ML162 or RSL3) led to substantial cell death whereas individual treatments did not (Figure 2A). Importantly, spermidine supplementation largely rescued cell death induced by combined sardomozide and ML162 treatment (Figure 2B), validating that the synthetic lethality results from on-target inhibition of polyamine biosynthesis. Because sardomozide is a polyamine analog (Figure S1D) and competes with spermidine for uptake (Figure S1E, Methods), exogenous spermidine may not fully restore intracellular polyamine levels, potentially explaining the incomplete rescue. Interestingly, high concentrations of polyamines in the medium re-sensitized cells to ferroptosis (Figure 2B), consistent with previous reports^35,36^ (see below).

**Figure 2:**
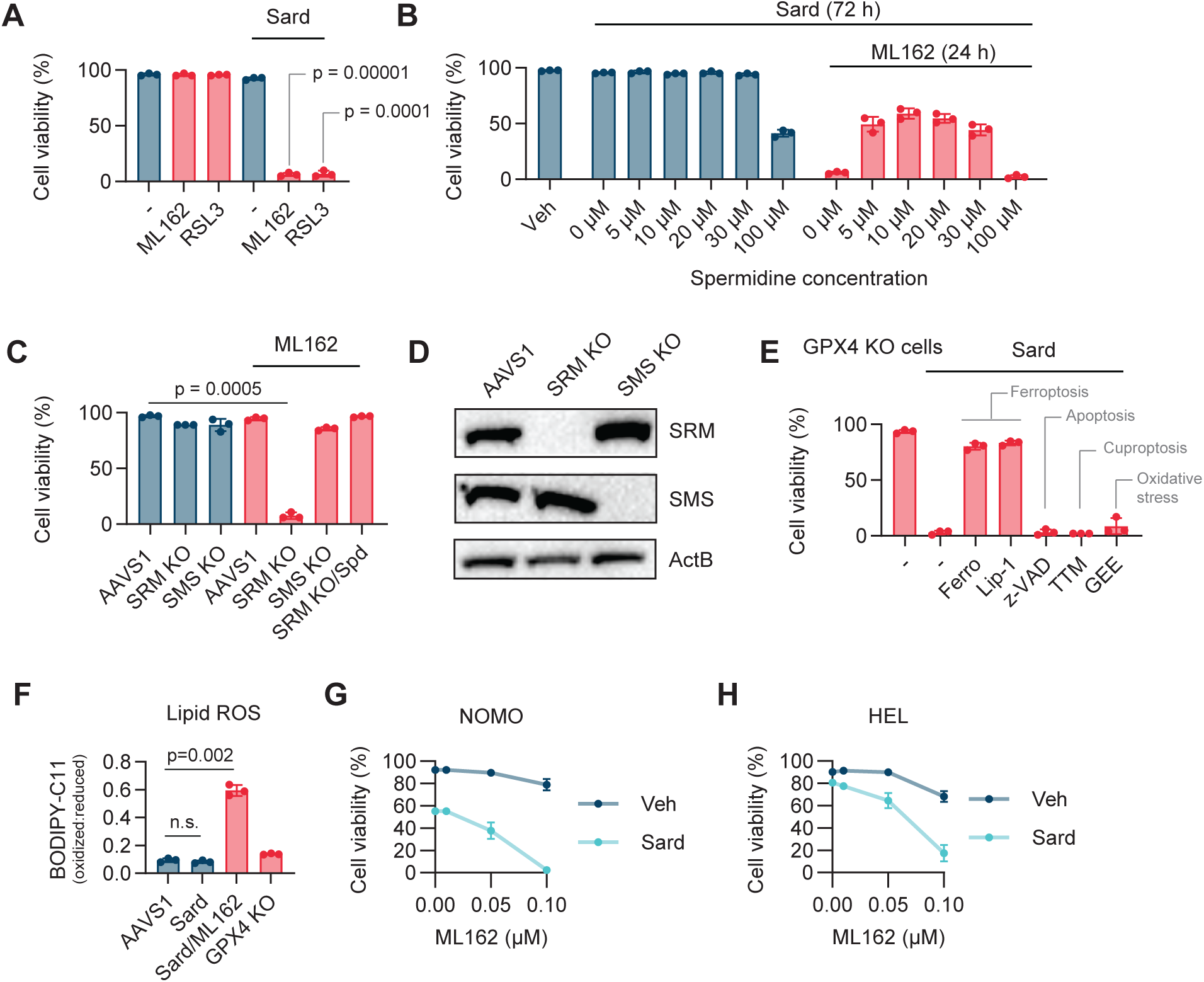
Polyamine depletion promotes ferroptosis. (A) Viability of K562 cells treated with ML162 (1 μM, 24 h) or RSL3 (2 μM, 24 h) after sardomozide pre-treatment (5 μM, 96 h) (n = 3). (B) Viability of K562 cells treated with ML162 (1 μM, 24 h) after sardomozide (5 μM, 72 h) plus the indicated spermidine concentrations (72 h) (n = 3). (C) Viability of AAVS1, SRM, and SMS KO K562 cells treated with ML162 (1 μM, 24 h); Spermidine (10 μM) was withdrawn from SRM KO cells 6 days before treatment (n = 3). (D) Immunoblot confirming SRM and SMS knockout (representative, n = 2 independent experiments). (E) Viability of GPX4 KO K562 cells treated with sardomozide (5 μM, 96 h) plus ferrostatin-1 (2.5 μM), liproxstatin-1 (1 μM), z-VAD-FMK (15 μM), TTM (tetrathiomolybdate, 10 μM), or GEE (glutathione ethyl ester, 100 μM) for 48 h (n = 3). (F) Lipid peroxidation measured as the C11-BODIPY oxidized:reduced fluorescence ratio in K562 cells after sardomozide (5 μM, 96 h) ± ML162 (1 μM, 2.5 h); mean fluorescence intensity ± SD (n = 3). (G-H) Viability of NOMO (G) and HEL (H) cells treated with ML162 (24 h) after sardomozide pre-treatment (5 μM, 72 h) (n = 3).

To dissect the contribution of individual polyamines to ferroptosis sensitivity, we used genetic perturbations. GPX4 inhibition induced substantial cell death in spermidine synthase (SRM) knock-out cells (Figure 2C) but not in control AAVS1-targeted cells. SRM knockout depletes both spermidine and spermine, resulting in the accumulation of the diamine putrescine. This vulnerability was rescued by spermidine supplementation, confirming the on-target activity of Cas9 (Figure 2C). In contrast, SMS knockout cells, which lack spermine but accumulate spermidine due to blocked conversion, were resistant to GPX4 inhibition (Figure 2C-D). These results suggest that spermidine is sufficient to protect against ferroptosis, while putrescine alone cannot maintain viability under GPX4 inhibition. While spermine depletion alone does not sensitize cells to ferroptosis, its role under physiological conditions cannot be excluded.

We tested whether ferroptosis inhibitors could rescue polyamine depletion-induced death under GPX4 inhibition. Treatment with known ferroptosis inhibitors, including lipophilic antioxidants^37^ ferrostatin-1 and liproxstatin-1 (also an iron chelator^38^), fully rescued GPX4 knockout cells from polyamine deprivation-induced death (Figure 2E). In contrast, inhibitors targeting other cell death pathways – such as apoptosis (z-VAD-FMK^39^), cuproptosis (tetrathiomolybdate^40^), and general oxidative stress (cell-permeable glutathione) – were ineffective (Figure 2E). Peroxidized lipid levels, assessed using the BODIPY-C11 probe^41^, were comparable across untreated controls, sardomozide-treated, and GPX4 knockout cells (Figure 2F). However, brief GPX4 inhibition (∼3 h, ML162) in polyamine-deficient cells markedly increased lipid peroxidation, consistent with ferroptotic lipid damage (Figure 2F). This synthetic lethal relationship was observed across multiple cell types including NOMO (human monocytic leukemia), HEL (human erythroleukemia), and MEL (mouse erythroleukemia) cells (Figure 2G-H, Figure S1F). Collectively, these results establish that polyamine depletion, specifically the loss of spermidine and spermine, creates a metabolic state selectively vulnerable to ferroptosis.

### Effects of polyamines are independent of major ferroptosis regulators

While our data demonstrate that polyamine depletion sensitizes cells to ferroptosis, recent studies report that excess intracellular polyamines can also promote ferroptosis. Excess spermidine and spermine are catabolized by the enzymes spermine oxidase (SMOX) and polyamine oxidase (PAOX), which convert them into lower-order polyamines such as putrescine while generating hydrogen peroxide and reactive aldehydes as byproducts^36^. These reactive intermediates can propagate lipid peroxidation, thereby sensitizing cells to ferroptotic cell death^35,42^. We therefore examined whether polyamine levels exhibit a biphasic relationship with ferroptosis sensitivity. Consistent with previous reports, high-dose spermidine supplementation (100 µM) increased susceptibility to GPX4 inhibition (Figure 3A). In contrast, lower concentrations used in our rescue experiments (10 µM), which are closer to the physiological extracellular levels^43^, were protective and restored viability in sardomozide-treated cells challenged with ML162 (Figure 2B). These findings indicate that both polyamine excess and deficiency can sensitize cells to ferroptosis, but likely through distinct biochemical routes, emphasizing that tight regulation of polyamine homeostasis is critical for redox balance.

**Figure 3:**
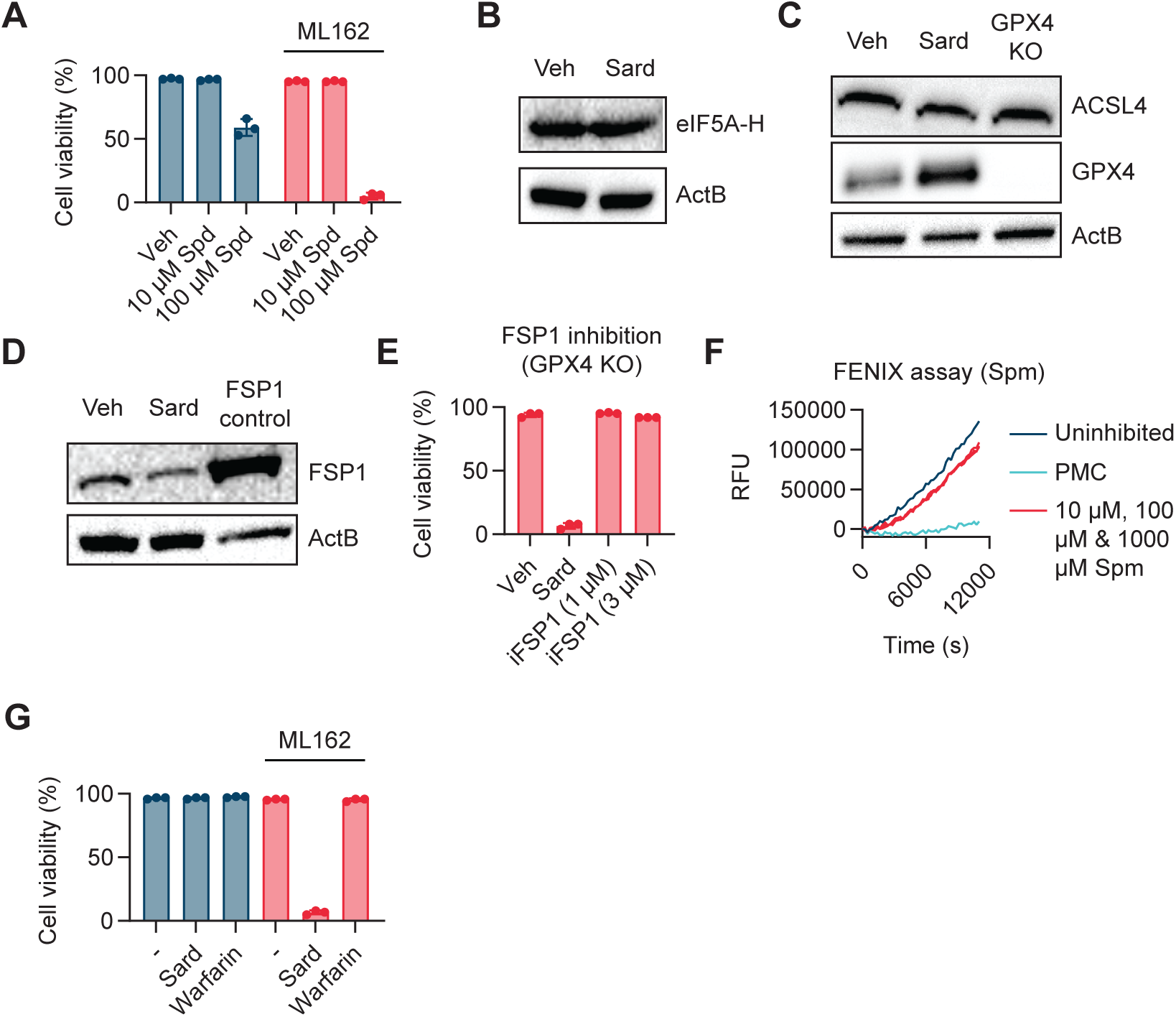
Polyamines act independently of canonical ferroptosis regulators. (A) Viability of K562 cells treated with ML162 (2 μM, 24 h) after spermidine pre-treatment (72 h) (n = 3). (B-D) Immunoblot of hypusinated eIF5A (B), ACSL4 and GPX4 (C), and FSP1 (D) in K562 cells treated with sardomozide (5 μM; 72 h in B, 96 h in C–D); OVCAR8, positive control. (E) Viability of GPX4 KO K562 cells treated with iFSP1 or sardomozide (5 μM, 96 h) (n = 3). (F) FENIX co-autoxidation traces with the indicated spermine concentrations or positive-control PMC (2,2,5,7,8-pentamethyl-6-chromanol; see STAR Methods). (G) Viability of K562 cells treated with ML162 (2 μM, 24 h) after warfarin (50 μM, 3 days) or sardomozide (5 μM, 72 h) pre-treatment (n = 3).

We next asked whether polyamine depletion sensitizes cells to ferroptosis through disruption of polyamine-dependent processes. Spermidine is required for hypusination of eIF5A, which regulates translation elongation, termination, and start codon selection^44,45^. If reduced hypusination was responsible for ferroptosis sensitivity, then polyamine depletion would be expected to decrease hypusinated eIF5A levels. However, acute sardomozide treatment under our assay conditions maintained normal hypusinated eIF5A levels (Figure 3B), likely because the existing hypusinated pool is stable and spermidine was incompletely depleted. Notably, direct hypusination blockade with GC7 sensitized cells to ML162-induced ferroptosis, an effect rescued by liproxstatin-1 (Figure S1G). Thus, while chronic hypusination loss can promote ferroptosis, it does not explain the acute effects of polyamine depletion in our system. Polyamine depletion has also been reported to induce cell-cycle arrest^46^. Because G1/S arrest can sensitize cells to ferroptosis in some contexts via downregulation of phospholipid remodeling enzymes^47^, we tested whether cell-cycle arrest alone could account for the ferroptosis phenotype. Pharmacologic G1 arrest with palbociclib effectively blocked cell-cycle progression (Figure S1H) but did not further reduce the viability of GPX4 KO cells (Figure S1I), indicating that cell-cycle arrest alone does not explain the increased ferroptosis sensitivity.

We then asked whether polyamine depletion sensitizes cells to ferroptosis by altering the expression or activity of known ferroptosis regulators. Besides GPX4, two key regulators of the ferroptosis pathway are: ACSL4 (acyl-CoA synthetase long-chain family member 4) which enriches membranes with polyunsaturated fatty acids (PUFAs) that serve as peroxidation substrates^34^, and FSP1 (ferroptosis suppressor protein 1) which suppresses ferroptosis by regenerating ubiquinol, a lipid-soluble antioxidant that scavenges lipid peroxyl radical^48,49^. ACSL4 and FSP1 levels were largely unchanged by polyamine depletion (Figure 3C-D). Interestingly, GPX4 expression increased (Figure 3C, bottom), likely reflecting a compensatory response to heightened ferroptosis vulnerability. To test whether polyamines might nevertheless modulate FSP1 activity independently of its abundance, we treated GPX4 knockout cells with the FSP1 inhibitor iFSP1. This had no effect on cell viability (Figure 3E), indicating that impaired FSP1 function does not explain the ferroptosis sensitivity induced by polyamine depletion.

Polyamines have also been proposed to function as antioxidants^50^, raising the possibility that their depletion sensitizes cells to ferroptosis by removing a radical-scavenging species. To test this possibility, we used the fluorescence-enabled inhibited autoxidation (FENIX) assay^51^, which monitors lipid peroxidation in native liposomes by tracking the oxidation of STY-BODIPY, a lipophilic fluorophore that is quenched upon reaction with lipid peroxyl radicals. Radical-trapping antioxidants (RTAs) compete with STY-BODIPY, thereby preserving fluorescence. Under our experimental conditions, polyamines did not exhibit any measurable lipid radical trapping activity (Figure 3F, Figure S1J). In contrast, PMC (2,2,5,7,8-pentamethyl-6-chromanol), a truncated vitamin E analog, robustly preserved fluorescence as expected (Figure 3F). These data indicate that direct radical scavenging by polyamines does not account for the ferroptosis sensitivity observed upon their depletion.

We then asked whether polyamine depletion compromises endogenous RTA pools. Untargeted metabolomics revealed NADPH, cysteine, and glutathione were largely unchanged upon polyamine depletion (Figure S1K), arguing against depletion of these canonical antioxidant pools. Because hydrophobic RTAs such as vitamin K and cholesterol-derived antioxidants are not reliably captured by standard metabolomics, we examined these pathways at the transcriptional level. We observed no major changes in the biosynthetic or salvage pathways of vitamin K, 7-dehydrocholesterol, coenzyme Q10, or tocopherol (Figure S1L-O). Consistent with this, inhibition of VKORC1-dependent vitamin K activation with warfarin^52^ did not increase sensitivity to GPX4 inhibition (Figure 3G), indicating that this arm of vitamin K recycling is unlikely to account for the observed ferroptosis sensitization. Because vitamin K can also be reduced to its radical-trapping form by FSP1 independently of this cycle^53^, this result does not by itself exclude a contribution from vitamin K-dependent RTAs; however, FSP1 inhibition did not reduce the viability of GPX4 KO cells (Figure 3E). Together, these results indicate that ferroptosis sensitization upon polyamine depletion is unlikely to be primarily mediated by compromised RTA pathways. Metabolomic profiling also revealed activation of the pentose phosphate pathway, reflected by increased levels of glucose-6-phosphate, 6-phosphogluconate, and downstream pentose intermediates, without expansion of the NADPH pool (Figure S2A). This pattern, where NADPH consumption outpaces production, is consistent with elevated redox demand^54^, suggesting that polyamine depletion increases oxidative burden through a mechanism other than RTA depletion.

Because ferroptosis sensitivity can also depend on oxidizable lipid substrates, we examined whether polyamine depletion alters the abundance of PUFA-containing phospholipids. Lipidomic profiling of intact lipid species, normalized to total annotated lipid signal, revealed only modest changes in lipid subtype abundance (Figures S2B-D, Table S2) and no upregulation of PUFA-containing phosphatidylcholine (PC) and phosphatidylethanolamine (PE) species after sardomozide treatment (Figures S2E-H, Table S2, see Methods). As this profiling measures intact lipid abundance rather than oxidized lipids, it reflects substrate availability rather than lipid peroxidation itself; polyamine depletion therefore does not substantially expand the pool of PUFA-containing PC and PE substrates. We also profiled transcriptional responses by RNA-seq, which revealed reduced expression of several ferroptosis-defense genes (Figure S2I). Although such changes could in principle modulate ferroptosis sensitivity, they did not produce a measurable antioxidant deficit: GPX4 protein increased rather than decreased (Figure 3C; bottom), and cysteine, glutathione, and NADPH pools were largely preserved (Figure S1K).

Collectively, these experiments indicate that the ferroptosis-sensitizing effects of polyamine depletion are unlikely to be explained by altered ACSL4 or FSP1 abundance, impaired hypusination of eIF5A, cell cycle arrest, compromised RTA pathways, or global expansion of PUFA-containing phospholipid substrates. Polyamine depletion is associated with elevated redox demand, yet polyamines themselves lack measurable lipid radical-scavenging activity under our assay conditions. While we cannot fully exclude contributions from these pathways, our findings strongly suggest that polyamine deficiency enhances ferroptosis susceptibility through a mechanism distinct from these established routes.

### Polyamine deficiency increases redox-active iron in cells

Given that ferroptosis is driven by iron-catalyzed lipid peroxidation, we examined cellular iron homeostasis. Polyamine depletion using sardomozide resulted in a dramatic increase in levels of ferritin (FTH1), the principal iron storage protein^55,56^ (Figure 4A). This induction mirrored the response to exogenous iron supplementation with ferric ammonium citrate (FAC) or ferrous ammonium sulfate (FAS) (Figure S3A). Similar ferritin upregulation was observed upon SRM knockout and was reversed by exogenous spermidine supplementation (Figure 4B), supporting that the ferritin response results from polyamine loss. Likewise, quantitative proteomics showed a modest but coherent induction of numerous proteins involved in maintaining iron homeostasis (e.g., FTL, CISD1, BOLA2, Figure S3B-C), consistent with increased iron stress in polyamine depletion condition.

**Figure 4:**
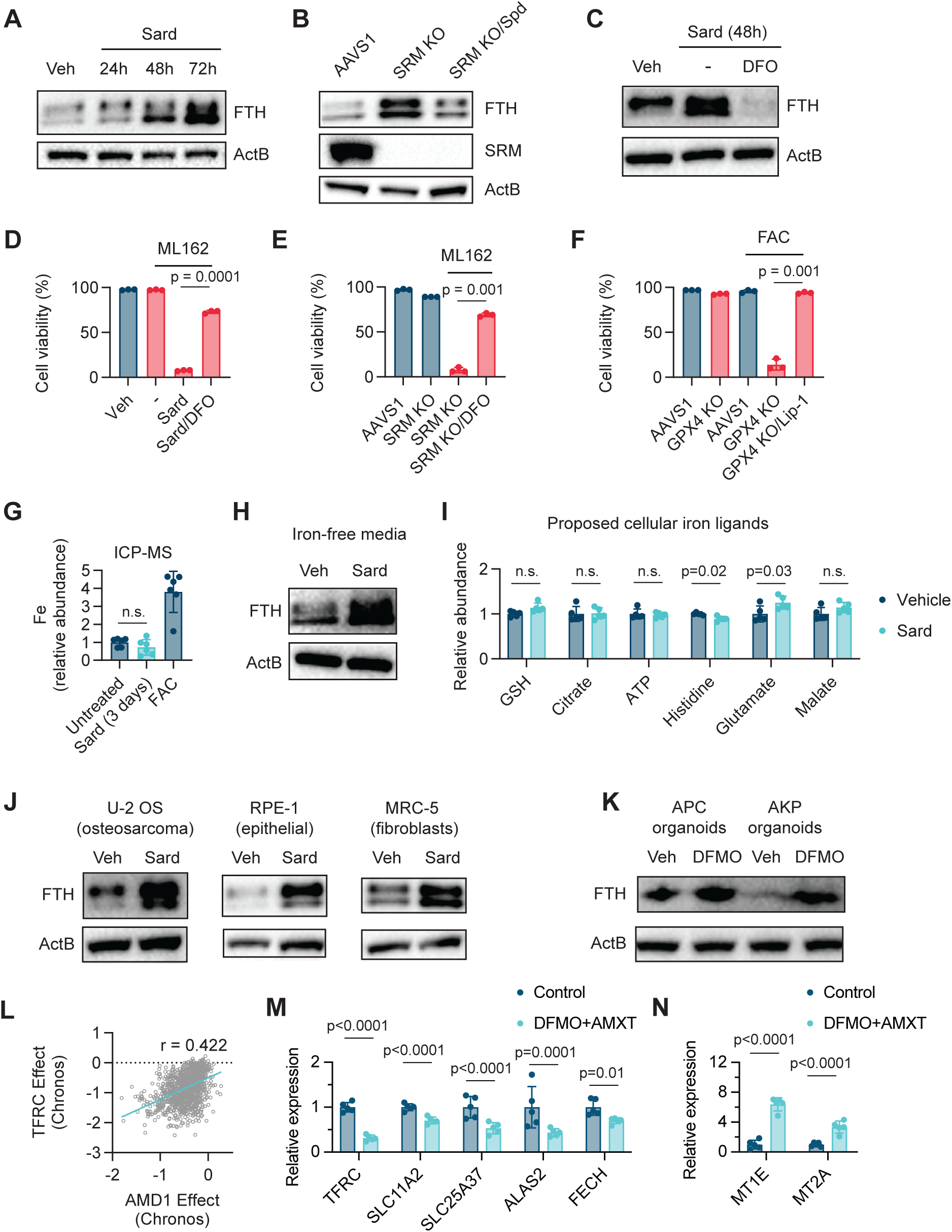
Polyamine deficiency increases redox-active iron. (A) Immunoblot of FTH in K562 cells treated with sardomozide (5 μM). (B) Immunoblot of FTH and SRM in SRM KO K562 cells; spermidine (10 μM) was withdrawn 6 days before lysis. (C) Immunoblot of FTH in K562 cells treated with sardomozide (5 μM, 48 h) with or without DFO (50 μM, 48 h). (D) Viability of K562 cells treated with ML162 (2 μM, 24 h) with or without DFO (100 μM, 24 h) after sardomozide pre-treatment (5 μM, 72 h) (n = 3). (E) Viability of AAVS1 and SRM KO K562 cells treated with ML162 (2 μM, 24 h); SRM KO cells supplemented with spermidine (10 μM), withdrawn 8 days before treatment; DFO (50 μM) added 24 h before ML162 (n = 3). (F) Viability of AAVS1 and GPX4 KO K562 cells treated with FAC (10 mM, 96 h) with liproxstatin-1 rescue (2 μM, 96 h) (n = 3). (G) ICP-MS quantification of cellular iron in K562 cells treated with sardomozide (5 μM, 72 h); FAC (2 mg/mL, 12 h) as positive control (n = 6). (H) Immunoblot of FTH in K562 cells cultured in iron-free IMDM with sardomozide (5 μM, 48 h); cells washed twice with HBSS before treatment. (I) Relative abundance of indicated metabolites in K562 cells treated with sardomozide (5 μM, 72 h) (n = 5). (J-K) Immunoblot of FTH in U-2OS, RPE-1, and MRC-5 cells treated with sardomozide (5 μM, 72 h) (J), or in APC and AKP intestinal organoids treated with DFMO (1 mM, 48 h) (K); DMSO vehicle control. (L) DepMap gene-effect correlation between TFRC and AMD1. (M-N) Relative mRNA abundance of indicated genes in DFMO/AMXT-1501-treated versus control K562 tumors (n = 5); edgeR FDR-adjusted P values.

Ferritin sequesters iron in a mineralized, non-reactive form, mitigating the risk of excessive redox-active iron accumulation^57^. We hypothesized that ferritin upregulation in polyamine-deficient cells represents an adaptive response to increased redox-active iron. Supporting this hypothesis, treatment with deferoxamine (DFO), an iron chelator, blunted the increase in ferritin levels from polyamine depletion (Figure 4C). We next tested whether elevated redox-active iron is functionally responsible for GPX4 dependence. DFO treatment substantially restored viability in polyamine-deficient cells (either treated with sardomozide or lacking SRM) challenged with ML162 (Figure 4D-E), supporting increased labile iron as a key driver of ferroptosis susceptibility. DFO rescued viability without major changes to the gene-expression program of polyamine-depleted cells (Figure S3D-E); the few differences were the expected iron-regulated genes such as TFRC, confirming on-target chelation (Figure S3D). The lipidomic profile of sardomozide plus DFO-treated cells was likewise similar to those with sardomozide treatment alone (Figure S3F-G, S4A-E, Table S2). Conversely, increasing the redox-active iron pool using FAC supplementation induced cell death specifically in GPX4 knockout cells, and this effect was fully rescued by the ferroptosis inhibitor liproxstatin-1 (Figure 4F). FAC had no impact on the viability of cells with functional GPX4 (Figure 4F), indicating that excess iron is not inherently toxic but becomes lethal when lipid peroxide repair is compromised. Collectively, these findings implicate increased redox-active iron accumulation as a critical mediator of ferroptosis under polyamine-deficient conditions.

We then investigated the source of this increased intracellular redox-active iron. Total intracellular iron, measured by inductively coupled plasma mass spectrometry (ICP-MS), showed negligible change upon polyamine depletion (Figure 4G). Thus, increased redox-active iron does not result from increased uptake or total intracellular iron but rather reflects a shift in the redox state or subcellular distribution of existing iron pools. Supporting this conclusion, ferritin levels were elevated even when polyamine-deficient cells were cultured in iron-free medium following extensive washing to remove extracellular iron (Figure 4H). Notably, this disruption was not explained by broad depletion of major abundant iron ligands^58^, including glutathione, ATP, and citrate, although histidine was modestly reduced and glutamate modestly increased (Figure 4I).

Finally, we asked whether the disruption of iron homeostasis induced by polyamine depletion is conserved across cell types. Sardomozide increased ferritin levels in U-2OS, RPE-1, and MRC-5 cells (Figure 4J). Extending these observations to a tissue-derived epithelial model, DFMO treatment also elevated ferritin levels in mouse colon cancer organoids with *Apc* mutations (APC) alone or in combination with *Kras*^G12D^, and *Trp53* mutations (AKP, a genotype observed in 40–45% colorectal cancer patients^59,60^) (Figure 4K). Consistent with a broader linkage between polyamine metabolism and iron programs, Cancer Dependency Map^61^ showed a striking correlation between cellular dependencies on AMD1, a rate-limiting enzyme in polyamine biosynthesis, and TFRC (transferrin receptor), the major mediator of cellular iron import (Pearson’s r□=□0.422, n = 1149 cell lines) (Figure 4L), with substantially stronger correlations in several lineages (e.g., prostate, r=0.906, n=10) (Figure S5A-E), indicating that cells with high iron demand (reflected by TFRC dependence) are especially reliant on polyamine synthesis.

To determine whether polyamine depletion similarly alters iron homeostasis in vivo, mice bearing subcutaneous K562 tumors expressing a genetically encoded polyamine reporter^62^ were treated with the polyamine biosynthesis inhibitor DFMO and the polyamine uptake inhibitor AMXT-1501^63^ (see Methods). Combined DFMO/AMXT-1501 treatment reduced reporter signal, confirming decreased polyamine availability in vivo (Figure S5F). RNA-seq of treated tumors revealed coordinated remodeling of iron and metal homeostasis pathways (Figure 4M-N, S5G), including reduced expression of iron uptake and trafficking genes (TFRC, SLC11A2/DMT1, SLC25A37/mitoferrin), induction of metallothioneins (MT1E, MT2A, and related family members), and suppression of heme biosynthetic genes (ALAS2, FECH). TFRC and IRE-containing SLC11A2/DMT1 transcripts are stabilized by IRP binding under iron-limited conditions so their reduction is consistent with IRP release and increased intracellular iron availability. Together, these changes are consistent with increased intracellular redox-active iron following pharmacologic polyamine depletion in vivo.

### Genetically encoded iron sensor reveals polyamine-iron buffering in living cells

Our biochemical and functional data support a model where polyamine depletion shifts intracellular iron toward a more redox-active state. To directly assess labile Fe^2+^ levels in cells, we used the small-molecule ferrous iron probe RhoNox-1^64^. RhoNox-1 fluorescence increased upon polyamine depletion, which was partially suppressed by DFO (Figure S5H-I), consistent with elevated labile Fe^2+^ in polyamine-deficient cells. Likewise, RhoNox-M^65^ confirmed elevated ferrous iron upon polyamine depletion, similar to that observed upon FAS supplementation (Figure S5J). However, both probes exhibited substantial cell-to-cell variability, likely due to differences in probe uptake, leakage, and their preferential localization in acidic compartments^66^. More broadly, existing small-molecule iron probes are sensitive to local pH, ionic environment, and interference from other metals including zinc and copper. Their variable uptake, poor photostability, and rapid clearance constrain their utility for quantitative single-cell analysis, long-term imaging, or use in complex tissues and organoids.

To overcome these limitations, we developed a genetically encoded reporter that quantifies redox-active iron in living cells with single-cell resolution. We harnessed the endogenous iron-responsive element (IRE)/iron regulatory protein (IRP) system: an evolutionarily conserved mechanism that monitors and regulates the labile iron pool^67^. IRPs bind to IRE hairpins in mRNAs encoding key iron regulatory proteins. When iron is low, IRP binding to 5′ UTR IREs (such as in ferritin and ferroportin mRNAs) blocks their translation, thus inhibiting iron sequestration and export^68^. Conversely, IRP binding to IREs in 3’UTRs (such as in transferrin receptor mRNA) enhances transcript stability and translation, thereby promoting iron uptake^69^. When labile iron concentrations rise, iron-sulfur clusters assemble on IRP1 while IRP2 undergoes degradation, releasing both proteins from their RNA targets.

We exploited this iron-sensing switch to engineer a genetically encoded fluorescent reporter (Figure 5A). We incorporated an optimized IRE configuration (Supplementary Note 1, Figure S5K-M) upstream of EBFP2. Translation initiation of EBFP2 depends on IRP release, providing a sensitive indicator of cellular redox-active iron. We also included an internal ribosome entry site (IRES)-mediated expression of EGFP downstream of EBFP2 as an internal normalization control, effectively accounting for cell-to-cell variability in mRNA abundance, transduction efficiency, and general translational status (Figure 5B). Sensor expression was further refined by placement under a doxycycline-inducible promoter, enabling tight temporal control and preventing saturation of the sensor’s dynamic range with prolonged expression. Importantly, reporter expression did not perturb endogenous iron metabolism, as evidenced by unchanged protein levels of iron metabolism enzymes, including FTH1, TFR1 and SLC40A1 (Figure S5N).

**Figure 5:**
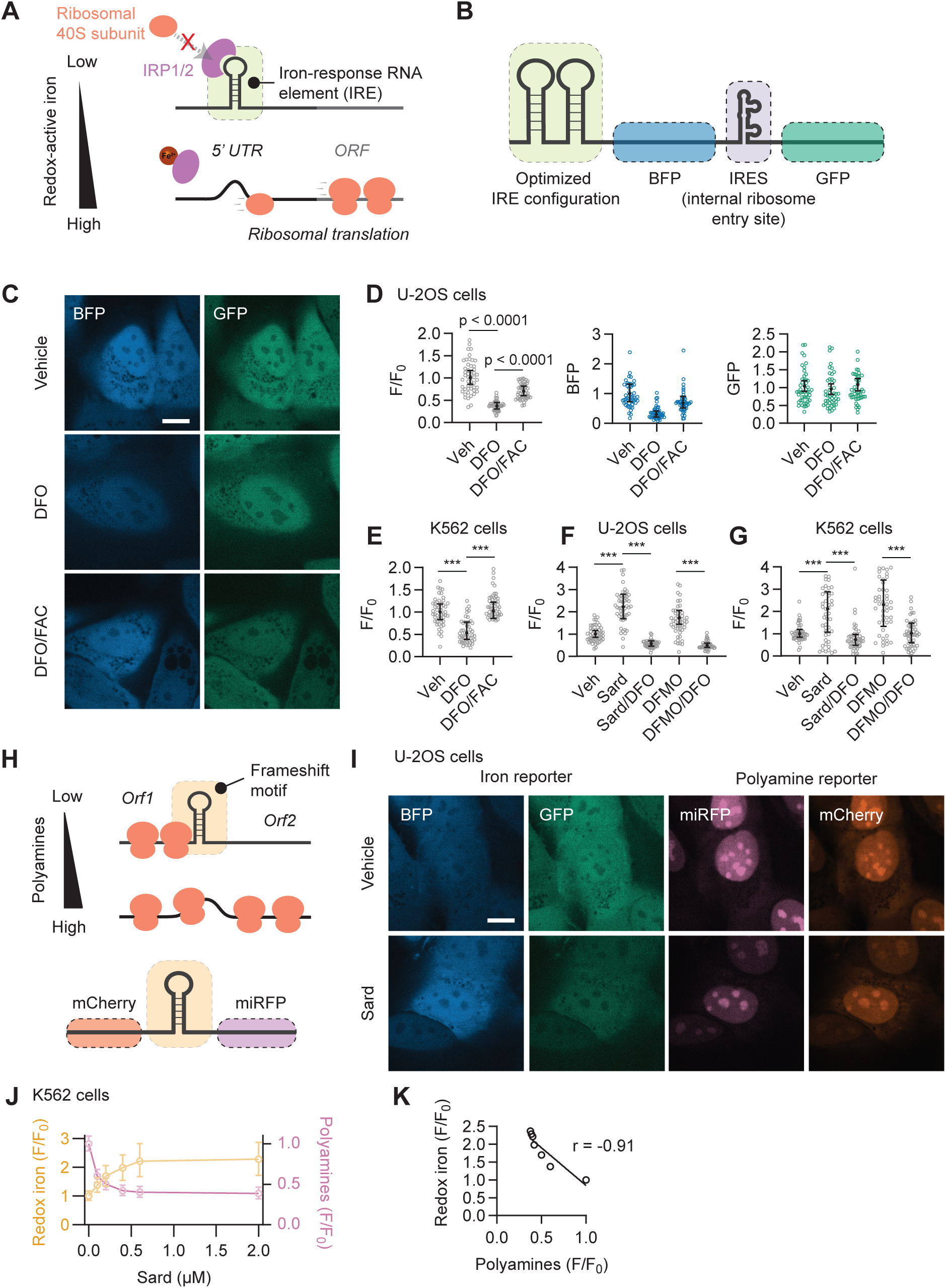
Genetically encoded sensor for redox-active iron. (A) Schematic of iron-responsive translation initiation of FTH mRNA. (B) Design of the redox-active iron sensor. (C-D) Representative fluorescence micrographs with BFP and GFP signals shown separately (C) and flow cytometry quantification (D) of translation initiation efficiency (F = BFP/GFP, F_0_ = F in vehicle-treated cells) in U-2OS cells expressing the iron sensor; DFO (100 μM, 48 h) with or without FAC (10 mM, 24 h). (E) As in (D), for K562 cells; DFO (50 μM, 48 h) with or without FAC (10 mM, 24 h). (F-G) As in (E), for U-2OS (F) and K562 (G) cells; sardomozide (5 μM, 48 h) or DFMO (0.5 mM, 48 h), each with or without DFO (100 μM, 48 h). (H) Design of the polyamine sensor. (I) Representative micrographs of U-2OS cells co-expressing polyamine and iron sensors under indicated treatments; sardomozide (5 μM, 48 h). (J) Redox-active iron (yellow) and polyamine (purple) reporter outputs across a sardomozide titration in K562 cells. (K) Linear regression of redox-active iron versus polyamine reporter outputs. Scale bars, 10 μm (C, I). For (D–G), median ± interquartile range from ≥100 cells; 50 cells shown; ≥2 independent experiments. For (J), median ± interquartile range from ≥100 cells. ***p < 0.0001.

To validate the reporter, we measured changes in the fluorescence ratio (F = EBFP2/EGFP), normalized to untreated cells (F_0_), to correct for imaging variations, as a function of redox-active iron. DFO treatment (100 μM, 48 h) markedly reduced F/F_0_ in U-2OS cells by ∼2.8 fold (0.36 ± 0.10 compared to the baseline of 1.00 ± 0.23), consistent with reduced intracellular redox-active iron. FAC supplementation (10 mM, 24 h) partially restored the signal (F/F_0_ = 0.70 ± 0.16) (Figure 5C-D), confirming the sensor’s responsiveness to iron availability. Similar trends were observed in K562 cells (Figure 5E), indicating that the sensor functions across cell types. We next leveraged our reporter to investigate the relationship between polyamine levels and redox-active iron. Treatment with sardomozide (5 μM for 48 h) or DFMO (0.5 mM for 48 h) increased F/F_0_ from 1.00 ± 0.23 to 2.22 ± 0.81 and 1.74 ± 0.45, respectively (Figure 5F), indicating increased redox-active iron. Co-treatment with DFO (100 μM for 48h) reversed this elevation, confirming that the signal reflected iron-dependent changes (Figure 5F). Similar iron accumulation upon polyamine depletion was observed in K562 cells (Figure 5G).

To definitively link these two metabolic pools at a single-cell level, we co-expressed our iron sensor alongside a genetically encoded polyamine reporter^62^. This sensor employs a polyamine-responsive ribosomal frameshifting region derived from *OAZ1* cloned between mCherry and miRFP670-2 (Figure 5H). In this design, miRFP670-2 is produced only upon polyamine-stimulated frameshifting, providing a readout of intracellular polyamine concentrations, while mCherry serves as a normalization control. This dual-reporter approach enabled simultaneous and quantitative measurement of both pathways within individual cells. We treated cells with a titration of sardomozide to induce a range of intracellular polyamine concentrations, and then captured both reporters in the same cells (Figure 5I-J). Strikingly, we observed a robust inverse relationship between the two reporter readouts (Pearson’s r, between (F/F_0_)_iron_ and (F/F_0_)_polyamine_ = -0.91), indicating that polyamine depletion is proportionally coupled to an increase in redox-active iron (Figure 5K). Altogether, these data demonstrate a tight, inverse coupling between polyamine and labile iron levels at the single-cell level, and establish our genetically encoded reporter as a robust tool for the quantitative dissection of iron biology.

### Polyamines coordinate Fe^2+^ and suppress iron reactivity

Why does polyamine depletion lead to an increase in redox-active iron? One hypothesis is that polyamines normally buffer intracellular iron through direct coordination. Their amine groups bear lone electron pairs capable of coordinating vacant orbitals in metal ions, forming stable Werner-type coordination complexes^70^. At physiological pH, polyamines are expected to be predominantly protonated and lack available lone electron pairs for metal coordination^71^. However, in cells, polyamines are extensively complexed with negatively charged biomolecules, including nucleic acids, ATP, and acidic phospholipids^2^. These electrostatic interactions may shift the local protonation equilibria, partially unmasking nitrogen lone pairs and enabling metal ion coordination^72,73^.

We therefore tested whether polyamines can coordinate ferrous iron under defined cell-free conditions. We first used FerroOrange as an *in vitro* readout of labile Fe^2+^. FerroOrange fluoresces upon irreversible reaction with Fe^2+^, and reduced signal indicates less Fe^2+^ accessible to the probe^74^. Both spermine and spermidine significantly reduced FerroOrange fluorescence in a concentration-dependent manner (Figure 6A), consistent with polyamine-dependent sequestration of Fe^2+^. As a positive control, DFO abolished the FerroOrange signal, confirming probe specificity (Figure 6A). Notably, increasing the solution pH to 10.5 to favor deprotonation enhanced the Fe^2+^-sequestering activity of both spermidine and spermine (Figure S6A), consistent with an amine-mediated coordination interaction.

**Figure 6:**
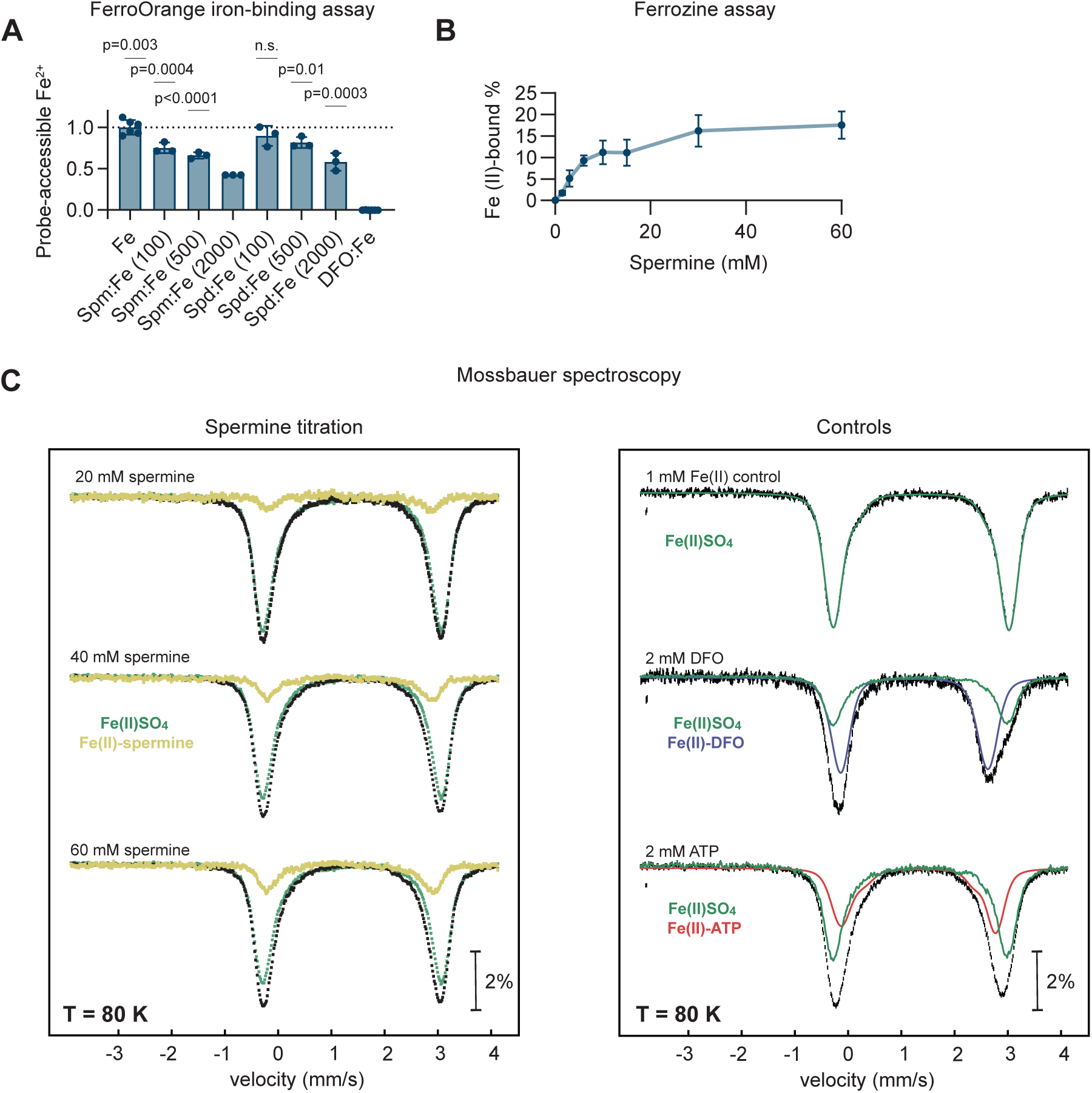
Polyamines can chelate metal ions. (A) Relative FerroOrange fluorescence, reporting probe-accessible Fe^2+^ (free-iron) as a function of increasing concentrations of the indicated polyamines; DFO, positive control (n ≥ 3). (B) Effect of increasing spermine concentrations on formation of ferrozine-Fe^2+^ complex (562 nm); fraction of Fe^2+^ displaced from ferrozine plotted as a function of spermine concentration (n = 3). (C) Effects of spermine and other chelators on the ^57^Fe^2+^ Mössbauer spectra (80 K, zero field; 1 mM ^57^Fe^2+^, 50 mM HEPES pH 7.5). Experimental spectra are shown as black vertical bars. Increasing spermine shifts spectra from unbound high-spin ^57^Fe^2+^ (δ = 1.38 mm/s, ΔE_Q_ = 3.31 mm/s; green) toward spermine-bound Fe²□ (yellow); DFO- and ATP-bound Fe²□ (blue, red) shown for comparison (2 mM each). Representative of 2 independent preparations.

In a complementary approach, we performed a ferrozine-based competition assay. Ferrozine forms a chromogenic complex with Fe^2+^ and competition by an alternative chelator would reduce this signal. Addition of spermine decreased ferrozine-Fe^2+^ complex formation in a concentration-dependent manner (Figure 6B). Because ferrozine is a substantially stronger Fe^2+^ chelator, spermine did not fully displace it even at large molar excess (Figure 6B). Nevertheless, the concentration-dependent reduction in signal demonstrates that polyamines can compete with ferrozine for Fe^2+^ binding.

To further characterize polyamine-iron binding, we performed Mössbauer spectroscopy, which provides a direct readout of iron redox states and the local coordination environment^75^. Perturbations in the Fe chemical environment by polyamine binding will be reflected as changes in the isomer shift (δ) and quadrupole splitting (ΔE_Q_) parameters, while the extent of binding can be quantified by the relative populations of distinct Fe species. In the presence of polyamines, we observed spectral signatures consistent with formation of a specific Fe^2+^ coordination complex (Figure 6C; left). Addition of spermine at a 20-fold molar excess relative to Fe^2+^ produced small but reproducible spectral shifts corresponding to ∼3% of total iron in spermine-Fe^2+^ complex. Increasing spermine to 60-fold molar excess further enhanced these shifts, corresponding to ∼15% of the total iron in spermine-Fe^2+^ complex, demonstrating concentration-dependent chelation of ferrous iron by spermine. Upon spermine binding, the Mössbauer parameters shifted from those of unbound high-spin Fe^2+^ (δ = 1.38 mm/s; ΔE_Q_ = 3.31 mm/s) to lower values (δ = 1.34 mm/s, ΔE_Q_ = 3.11 mm/s). These changes indicate that Fe^2+^ remains high-spin, while the decreases in δ and ΔE_Q_ reflect increased s-electron density at the nucleus and a more symmetric ligand field, consistent with coordination by a weak-field ligand^76^. These results provide direct evidence for polyamine-Fe^2+^ complex formation and the spectral shifts are consistent with the planar nitrogen-based chelation geometry observed in crystal structures for polyamine-metal complexes^77^. Control ligands (ATP and DFO) produced distinct reference spectra with reduced parameters without altering spin state (Figure 6C; right), further supporting that the observed shifts reflect specific polyamine-dependent Fe^2+^ coordination rather than oxidation or nonspecific perturbation. Because labile iron in cells is thought to exist as a heterogeneous pool of weakly bound species, including complexes with abundant metabolites such as ATP and citrate^58^, we asked whether spermine-dependent Fe^2+^ coordination could still be detected in the presence of ATP. In samples containing equimolar Fe^2+^ and ATP, addition of spermine produced a distinct spermine-dependent spectral component (Figure S6B), indicating that ATP does not abolish this interaction. To test whether this interaction depends on polyamine structure, we compared tetraamine spermine with diamine putrescine under matched conditions. At pH 7.5, spermine produced a stronger perturbation of the Fe^2+^ spectrum than putrescine (Figure S6C). This distinction became more pronounced at pH 10, where amine deprotonation is favored: spermine produced a substantial Fe^2+^-bound spectral component, whereas putrescine binding remained negligible (Figure S6D).

The iron concentrations in these assays (micromolar for the optical measurements and millimolar ^57^Fe^2+^ for Mössbauer spectroscopy) were set by the requirements for robust competition and spectroscopic detection, and are not intended to model the absolute concentration of the cellular labile iron pool. Under simple ligand-excess equilibria, the fraction of Fe^2+^ coordinated depends primarily on free ligand concentration and apparent affinity rather than on total iron. Thus, the fractional binding observed here should be preserved at the sub-micromolar iron and millimolar polyamine concentrations present in cells. Collectively, our data support a model where polyamines contribute to low-affinity, high-abundance buffering of the labile, redox-active iron pool, thereby limiting its accessibility and iron-catalyzed lipid peroxidation (Figure 7). Disruption of this buffering system through polyamine depletion liberates this iron, sensitizing cells to ferroptotic death. More broadly, these findings uncover an unexpected layer of iron regulation by polyamines, linking two fundamental metabolic networks with relevance to redox stress, cell death, and disease.

**Figure 7:**
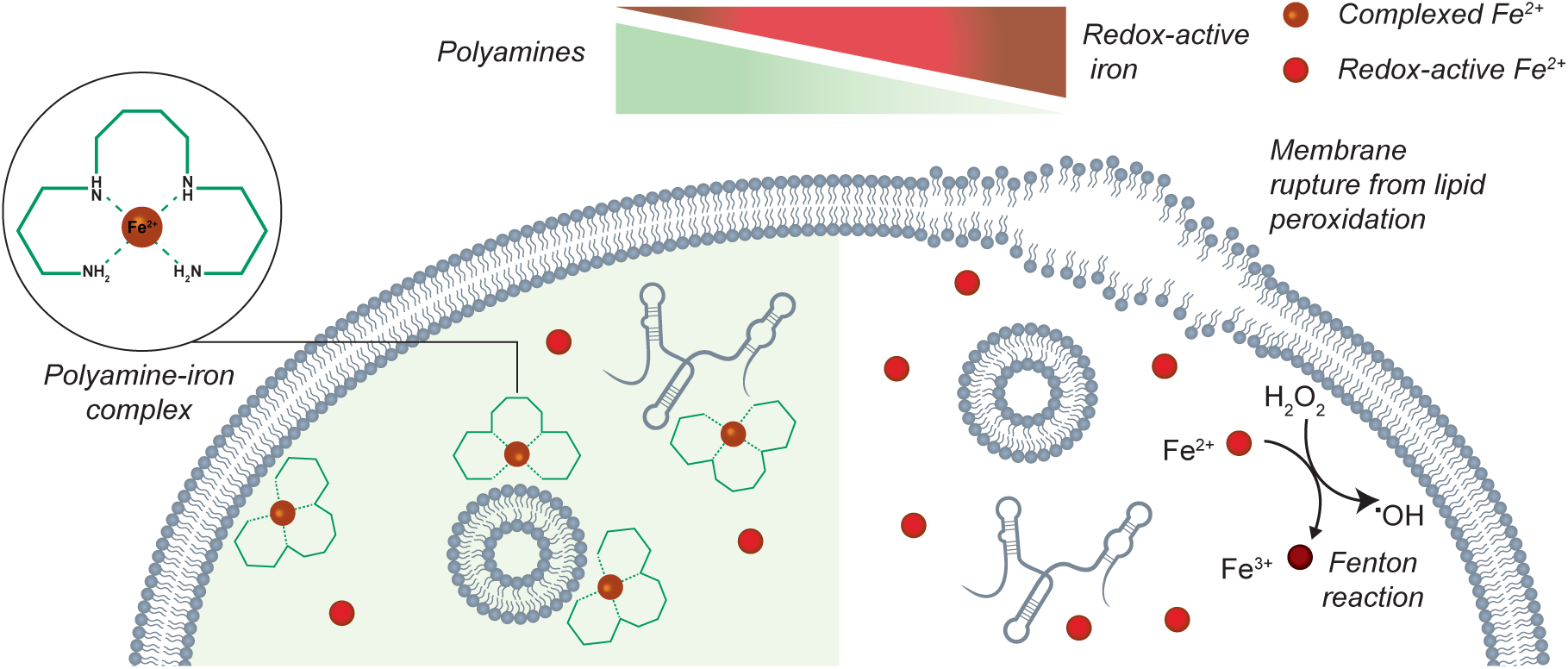
Model of polyamine-mediated iron buffering. Polyamine depletion reduces Fe^2+^ buffering, increasing accessible, redox-active iron that drives Fenton chemistry, lipid peroxidation, membrane rupture, and ferroptosis.

## Discussion

Our finding that polyamines regulate iron homeostasis and ferroptosis sensitivity is relevant to ongoing efforts to target polyamine metabolism in cancer and other diseases (NCT06892678, Ewing sarcoma; NCT05594563, type 1 diabetes; NCT04301843, neuroblastoma; NCT04696029, medulloblastoma; NCT06976424, hirsutism). Elevated polyamine biosynthesis is a long-recognized hallmark of proliferating cancer cells: comparative RNA-seq^78^ shows upregulation of biosynthesis enzymes (e.g., ODC1, SRM) in tumors compared to normal tissues (Figure S7A-B, Table S3), driven by oncogenic signaling pathways (MYC^79^, RAS-RAF-MEK^80,81^, AKT^82^, and mTORC1^83^) and required for the tumorigenic activity of several of these oncogenes^84,85^. Polyamines are also critical for metastatic dissemination^86–90^ and immune evasion^91^. DFMO has shown promise in neuroblastoma maintenance therapy^92^ and chemoprevention^93^, but standalone efficacy has been limited^94^. Our results identify several actionable targets (GPX4, BCL2L1 and ATP13A3), providing a foundation for combination therapies to enhance the efficacy of polyamine depletion in cancer treatment. While this manuscript was in preparation, one study reported that DFMO prevents metastasis in Ewing sarcoma by inducing ferroptosis signatures^95^ and Li et al. independently identified spermine as an endogenous iron chelator that suppresses ferroptosis^96^, showing that inhibiting spermine production promotes ferroptosis and impairs hepatocarcinogenesis, while spermine supplementation protects against ferroptosis-associated ischemia-reperfusion injury.

This model also raises the question of whether polyamine metabolism is itself regulated by cellular iron status. Previous studies have shown that iron depletion using chelators such as DFO reduces cellular spermidine and spermine levels^97^. Mechanistically, iron depletion impairs the methionine salvage pathway, reducing S-adenosylmethionine (AdoMet), which provides the precursor for the aminopropyl donor required to synthesize spermidine and spermine; it also alters multiple polyamine pathway components, decreasing ADI1 and MAT2α and increasing SAT1, and reduces spermidine uptake^97^.

Conversely, elevated iron has been linked to increased polyamine biosynthesis: iron overload induces ODC1 through WNT/MYC signaling^35^, and loss of Nrf2 or iron supplementation upregulates Odc1^98^. Together with our finding that polyamine depletion increases redox-active iron and induces ferritin, these studies indicate that iron and polyamine metabolism are intricately coupled within a homeostatic loop that tunes polyamine biosynthesis, catabolism, and import to iron status. Defining the molecular wiring and physiological consequences of this bidirectional regulation is an important direction for future work.

The polyamine-iron partnership may be ancient: as early life forms evolved to exploit iron’s catalytic power, polyamines could have been co-opted to stabilize redox-active iron and suppress its toxicity. This dual role of polyamines in supporting macromolecular synthesis while buffering metal-induced damage explains the extraordinary investment in maintaining polyamines at millimolar levels, safeguarded by tightly regulated biosynthetic, catabolic, and transport pathways. By revealing polyamines as redox buffers, our work recasts these elaborate regulatory mechanisms as rapid-response systems for managing cellular iron, with implications spanning bacterial pathogenesis and cold tolerance^99^ to cancer therapy.

This framework is also consistent with clinical and genetic observations linking polyamine metabolism to metal homeostasis. Polyamines coordinate transition metals including copper, nickel, cobalt, lead, and zinc^100–103^ with nitrogen donors arranged in a square-planar geometry around the central metal ion^77^. This structure closely resembles the metal-binding architecture observed in other nitrogen-containing biological macrocycles such as porphyrins^104^ and is consistent with our Mössbauer spectroscopy showing nitrogen-based coordination in Fe^2+^-spermine complexes. Trientine^105^, a tetraamine structurally similar to spermine (Figure S7C), is a first-line treatment for copper toxicity in Wilson’s disease. Patients treated with trientine often exhibit side effects associated with iron deficiency, suggesting that it has broader metal-chelating properties beyond copper^106,107^. Conversely, polyamine depletion using DFMO frequently causes iron dysregulation side effects, including anemia and thrombocytopenia^108^. Loss of function mutations in ATP13A2, a polyamine transporter linked to early-onset Parkinson’s disease, lead to iron accumulation in basal ganglia and cultured dopaminergic neurons^109–112^. Notably, before ATP13A2 was recognized as a polyamine transporter, it was shown to protect cells against broad metal-induced toxicity, including from Fe^3+^, Mn^2+^, and Zn^2+113–117^. Similarly, loss-of-function mutations in ATP13A3, the primary polyamine transporter in non-neuronal cells^62^, are associated with pulmonary arterial hypertension^118^, a disease often linked to altered iron metabolism^119^.

Beyond these genetic disorders, the polyamine-iron axis may also influence age-related physiological decline. Polyamine levels decline with age which may result in a progressive reduction in iron buffering, contributing to increased oxidative stress and ferroptosis vulnerability^120^. Consistent with this possibility, aged alveolar type 2 (AT2) cells in vivo exhibit elevated labile iron levels, impacting stemness and tumorigenesis^121^, although a direct role of age-related polyamine loss remains to be tested. Conversely, elevated polyamine synthesis in cancer cells might serve a dual purpose: supporting rapid proliferation while simultaneously protecting against oxidative stress arising from dysregulated metabolism. Notably, both polyamine^122^ and iron metabolism^123–125^ are upregulated in cancer stem cells.

One outstanding question is how polyamines coordinate iron given their predominantly protonated state at physiological pH. Our experiments at alkaline pH demonstrate that deprotonation substantially enhances iron chelation, while Mössbauer spectroscopy shows that spermine-dependent Fe^2+^ coordination is still detectable at pH 7.5. At intracellular stoichiometries where polyamines exceed labile iron by orders of magnitude, even a small deprotonated or coordination-competent fraction could provide substantial buffering. Moreover, polyamines associate extensively with anionic biomolecules (RNA, ATP, phospholipids), which may shift local protonation equilibria and facilitate ternary complex formation. We do not propose that polyamines are the sole ligands of the labile iron pool, or that they form a single, discrete Fe^2+^ species. Rather, at millimolar abundance, they likely contribute alongside other endogenous ligands to a heterogeneous, low-affinity/high-abundance ligand environment that restrains redox-active iron. This interpretation is consistent with current models of the labile iron pool as a set of weakly bound species rather than a single ligand class^58,126–128^.

Beyond iron buffering, polyamine depletion perturbs other pathways that could contribute to ferroptosis sensitivity, including reduced expression of SLC7A11 (cystine import); MTHFD2 and SHMT2 (one-carbon metabolism and NADPH production); and PHGDH and PSAT1 (serine biosynthesis) (Figure S3E). These changes could diminish the capacity of polyamine-depleted cells to mount adaptive antioxidant responses. However, iron chelation rescued ferroptosis sensitivity without broadly reversing this transcriptional state, and antioxidant pools and GPX4 protein levels were preserved or increased under polyamine-depleted conditions. Thus, while these transcriptomic and lipidomic changes may modulate ferroptosis sensitivity under polyamine depletion, our data point to redox-active iron as its principal driver.

An important advance in our work is the development of a genetically encoded fluorescent reporter for quantitative measurement of redox-active iron in living cells. Unlike chemical probes, which suffer from variable uptake, retention, and clearance, genetically encoded reporters provide stable expression, eliminate loading artifacts, and enable longitudinal tracking. Its single-cell resolution supports high-throughput screening for regulators of iron homeostasis, and pharmacokinetic studies of iron-targeting therapies, potentially in live animals. We anticipate that this reporter will provide a powerful platform to facilitate future discoveries into iron biology, ferroptosis, and disease mechanisms.

## Limitations of the study

Although iron chelation rescues the ferroptosis sensitivity caused by polyamine depletion, polyamine depletion is also accompanied by lipidomic, transcriptional, and likely additional cellular changes. While our findings support polyamine-mediated buffering of labile/redox-active iron as a key contributor to ferroptosis suppression, we do not exclude parallel or indirect contributions from lipid metabolism, transcriptional adaptation, or other context-dependent pathways. The precise molecular nature of the polyamine-iron species in cells also remains to be defined. We establish polyamine-dependent Fe^2+^ coordination under defined conditions, but do not directly quantify binding of low-micromolar labile Fe^2+^ by millimolar polyamines in physiological ligand mixtures. Our data are most consistent with a heterogeneous, low-affinity/high-abundance ligand environment rather than a single discrete complex; however, the composition, dynamics, subcellular distribution, and physiological contexts in which these species are most consequential remain to be established. Finally, our conclusions are drawn largely from cellular and organoid models. Although polyamine depletion remodeled iron and metal-homeostasis programs in tumors in vivo, this transcriptional signature does not directly measure the labile iron pool. Acute depletion of higher polyamines in vivo is challenging, because inhibition of de novo synthesis can be offset by compensatory uptake from extracellular, dietary, and microbiota-derived sources. Future studies combining tissue-specific polyamine perturbation with genetically encoded reporters of polyamine status and labile Fe^2+^ will help define how this pathway operates across tissues and in response to genetic, pharmacologic, and dietary modulation of polyamine levels.

## Supporting information

Key Resource Table

## RESOURCE AVAILABILITY

### Lead contact

Further information and requests for resources and reagents should be directed to the lead contact, Ankur Jain.

### Materials availability

All unique and stable reagents generated in this study are available from the lead contact upon completion of a Materials Transfer Agreement.

### Data and code availability

CRISPR screen datasets are deposited in GEO under accession GSE300179. Mapping code for aligning reads to the sgRNA library is available at https://github.com/whitehead/barc/tree/main/Perl. RNA-seq data are deposited at PRJNA1467631, and lipidomics data at 10.5281/zenodo.20706003. Further reanalysis information is available from the lead contact upon request.

## Acknowledgments

We thank Matthew Vander Heiden, Squire Booker, Catherine Drennan, Elizabeth Nolan, Naama Kanarek, Daniel Suess, Saleh Khawaled, Ömer Yilmaz, and Lindsey Backman for discussions, Punam Bisht for providing MRC-5 fibroblasts, George Bell and Xinlei Gao for data-analysis help, Alex Joseph for plasmid-preparation help, and Brighton A. Skeel, Ky Lowenhaupt, Mohamed El-Brolosy, Peter Tsevetkov, Reuben Saunders, Eric Smith, Sarah Willis, Walt Massefski, Mackenzie Field, and the Jain lab for feedback. We thank the Whitehead Institute Flow Cytometry Core and MIT Center for Environmental Health Sciences (CEHS) Core, supported by National Institute of Environmental Health Sciences (P30 ES002109). This work was supported by National Institute of General Medical Sciences (NIGMS) grant R35GM151111, the Bumpus Foundation, Pew Charitable Trusts, and the Chan Zuckerberg Initiative (all A.J.); Koch Institute Core Grant P30CA014051 (National Cancer Institute); the Kathy and Curt Marble Cancer Research Fund; the Howard Hughes Medical Institute Freeman Hrabowski Scholars Program (W.S.H.); R35-GM156452 (M.E.P.); the Ligue Contre le Cancer (Equipe Labellisée) and Fondation Charles Defforey–Institut de France (S.M., R.R.); and NIGMS training grant T32GM136540 (J.S., R.P.M.).

## Author contributions

P.S. designed, performed, analyzed, and interpreted most experiments. H.R.K. assisted with CRISPR-Cas9 screen. R.P.M. and J.S. performed experiments. L.G. performed Mössbauer spectroscopy and ferrozine assays with guidance from M.E.P. C.A. helped with image analysis. R.A. helped with RNA-extraction and sequencing. S.M. and I.S.P. helped synthesize critical reagents with guidance from R.R. and P.T.H., respectively. S.I. provided organoid samples. T.K. and M.W. assisted with metabolomics. B.Y. assisted with data analysis. A.D. assisted with lipidomics. L.F. performed the proteomics experiments. A.J. and W.S.H. supervised the project. P.S. wrote the manuscript with edits from A.J. and W.S.H. All authors reviewed and approved the final manuscript.

## Declaration of interests

P.S. and A.J. are inventors on a pending patent application based on this work. The remaining authors declare no competing financial interests.

## STAR Methods

### Experimental model and study participant details

#### For cell lines

U-2OS, K562, and RPE1 (immortalized) cell lines were obtained from ATCC. NOMO, HEL, and MEL cells were generously provided by Naama Kanarek (Boston Children’s Hospital), and MRC-5 fibroblasts by Punam Bisht (Whitehead Institute). Cell lines used were of the following reported sex: K562 (female), U-2OS (female), RPE1 (female), HEK293T (female), MRC-5 (male), NOMO-1 (female) and HEL (male). U-2OS, HEK293T, RPE1, and MRC-5 cells were cultured in DMEM (Gibco, Cat# 11965126), while K562, NOMO, HEL, and MEL suspension cells were maintained in RPMI-1640 (Gibco, Cat# 11875093). All media were supplemented with 10% (v/v) fetal bovine serum (FBS; Gibco, Cat# A56707-01, Lot# U2933097RP) and penicillin-streptomycin-glutamine (Gibco, Cat# 10378016). FBS was thawed at 4°C and used without further processing. For iron reporter assays, all cell types were cultured in RPMI supplemented with tetracycline-free cosmic calf serum (CCS) in place of FBS. All cell lines were maintained at 37°C in a humidified atmosphere containing 5% CO□ and were tested monthly for mycoplasma contamination.

#### For organoids

Mouse organoids were generated as part of a previous study^129^ and expanded for use here: briefly, colon organoids from Kras-LSL-G12D; Trp53 fl/fl; Rosa26-LSL-tdTomato mice were expanded in L-WRN medium. Organoids were transfected with pSECC (Addgene #60820: Cre, Cas9, U6-driven sgRNA cassette) carrying sgApc (5′-GTCTGCCATCCCTTCACGTT-3′), delivered via Lipofectamine 2000 (Thermo Fisher, 11668027) by spinoculation. Edited, Cre-active clones were selected by withdrawing WNT3A/R-spondin-1 and adding 10 μM nutlin-3 (Sigma-Aldrich, N6287). Apc-null organoids were generated by infecting wild-type colon organoids with a lentiviral vector encoding Cas9, GFP, and the sgApc guide, delivered by spinoculation, followed by selection via Wnt3a/R-spondin-1 withdrawal alone.

### For in vivo animal studies

#### Mice

Mouse studies were performed in the Koch Institute for Integrative Cancer Research at MIT. Mice were housed under a 12-h light/12-h dark cycle, temperatures in the range 20–22□ °C and humidity between 30–70%. Mouse experiments performed in this study received approval by the MIT Institutional Animal Care and Use Committee. Tumors were not allowed to exceed 1.5□cm in diameter. Animals showing discomfort or distress were humanely euthanized following the MIT Institutional Animal Care and Use Committee guidelines. All experiments were conducted in female NSG mice (Jackson: 005557), aged 8-10 weeks.

### Method details

#### Chemicals

The following reagents were purchased: spermidine trihydrochloride (S2501, Sigma-Aldrich), spermine tetrahydrochloride (J63060.14, Thermo Fisher Scientific), dimethyl sulfoxide (DMSO; 276855, Sigma-Aldrich), DL-α-difluoromethylornithine (DFMO; 16889, Cayman Chemical), iFSP1 (HY-13607, MedChemExpress), warfarin (1719000, Sigma-Aldrich), FerroOrange (F374, Dojindo), BODIPY™ 581/591 C11 (D3861, Thermo Fisher Scientific), liproxstatin-1 (HY-12726, MedChemExpress), RSL3 (HY-100218-A, MedChemExpress), ML162 (HY-100002, MedChemExpress), ferrostatin-1 (Cayman Chemical), Z-VAD(OH)-FMK (Cayman Chemical), STY-BODIPY (Cayman Chemical), PMC (HY-111024, MedChemExpress), A-1331852 (HY-19741, MedChemExpress), iron(II) chloride (220299, Sigma-Aldrich), ferrozine (82950-1G, Sigma-Aldrich), BODIPY 493/503 (MedChemExpress), deferoxamine mesylate (DFO; Cayman Chemical), ferric ammonium citrate (F5879, Sigma), Ellman’s reagent (22582, Thermo), oleic acid (O3008, Sigma-Aldrich), and sardomozide dihydrochloride (HY-13746B, MedChemExpress).

#### Lentivirus production for CRISPR-Cas9 screen

15 × 10^6^ HEK-293T cells were seeded in T175 cm^2^ flasks in DMEM (Thermo Fisher Scientific #12430054) supplemented with 10% fetal bovine serum (GeminiBio #100-106). After 24 hours, the media was changed to 20 mL viral production medium: IMDM (Thermo Fisher Scientific #1244053) supplemented with 20% inactivated fetal serum (GeminiBio #100-106). At 32 hours post-seeding, cells were transfected with a mix containing 76.8 µL Xtremegene-9 transfection reagent (Sigma Aldrich #06365779001), 3.62 µg pCMV-VSV-G (Addgene plasmid # 8454)^130^, 8.28 µg psPAX2 (a gift from Didier Trono; Addgene plasmid # 12260), and 20 µg sgRNA/Cas9 plasmid and Opti-MEM (Thermo Fisher Scientific #11058021) to a final volume of 1 mL. Media was changed 16 hours later to 55 mL fresh viral production medium. The virus was collected at 48 hours post-transfection and filtered through a 0.45 µm filter, aliquoted, and stored at -80 °C until use.

#### CRISPR-Cas9 screen

A genome-wide lentiviral sgRNA library^26^ in a Cas9-containing vector comprising 97,888 unique sgRNA sequences with ∼5 sgRNAs per target was used to transduce 390 × 10^6^ K562 cells to achieve an MOI < 1 (10-20% transduction efficiency) and ∼500-1000-fold library coverage. Briefly, polybrene (10 µg/mL final concentration) and virus were mixed with cells (2.5 × 10^6^ cells/mL final density) and distributed into individual wells in 6-well plates. Plates were centrifuged at 1126 *g* for 45 minutes at 37 °C and transferred to an incubator. After 8 hours, cells were pelleted, the virus was removed, cells were resuspended in the fresh growth medium, and transferred to T225 cm^2^ flasks (250,000 cells/mL final density). After 36 hours, cells were collected and reseeded in fresh growth medium (200,000 cells/mL final density) and puromycin was added (3 µg/mL final concentration). After 3 days, cells were collected and transduction efficiency was determined by comparison of cell survival of transduced cells relative to control cells (untransduced and unselected). Cells were passaged every 2 days (0.2 × 10^6^ cells/mL) before DFMO treatment (0.5 mM) starting 9-days post-transduction. At the screen endpoint (14 days DFMO treatment), cell pellets were collected from flasks and frozen at –80 °C.

#### Sequencing library preparation

Genomic DNA (gDNA) was extracted from cell pellets of 20 x 10^6^ cells using the Blood genomicPrep Mini Spin Kit (Cytiva # 28904264) or 60 × 10^6^ cells using the QIAamp DNA Blood Maxiprep Kit (Qiagen # 51192) according to manufacturer’s instructions with minor modifications: Cytiva and QIAGEN Protease were replaced with a 10 mg/mL solution of ProteinaseK (MilliporeSigma # 3115879001) in water; cells were lysed overnight; centrifugation steps after Qiagen Buffer AW1 and AW2 were performed for 2 minutes and 5 minutes, respectively; 2 x 30 µL (Cytiva) or 1 mL (Qiagen) of water preheated to 70 °C was used to elute gDNA (1- or 5-minute incubation), followed by centrifugation for 1 or 5 minutes (Cytiva or Qiagen extractions, respectively). gDNA was quantified using the Qubit dsDNA HS Assay kit (Thermo Fisher Scientific #Q32851).

All PCR reactions were performed in 50 µL reactions using ExTaq Polymerase (Takara Bio #RR001B) using the following primers:

Forward: 5’- AATGATACGGCGACCACCGAGATCTACACCCCACTGACGGGCACCGGA - 3’

Reverse: 5’- CAAGCAGAAGACGGCATACGAGATCnnnnnnTTTCTTGGGTAGTTTGCAGTTTT - 3’

Where “nnnnnn” denotes the barcode used for multiplexing.

For all samples, 1, 3, or 6 µg of gDNA was initially amplified for 28 cycles in 50 µL test PCR reactions. Subsequently, an additional 50 µL reactions were performed using 6 µg per reaction (140 µg gDNA).

Reactions were pooled and 100 µL of each sample was purified using HighPrep PCR beads (MagBio Genomics #AC-60005), eluted with 20 µL water, and quantified using the Qubit dsDNA HS Assay kit before sequencing for 26 cycles on an Illumina NovaSeq using the following primers:

Read 1 sequencing primer: 5’- GTTGATAACGGACTAGCCTTATTTAAACTTGCTATGCTGTTTCCAGCATAGCTCTTAAAC - 3’ Index sequencing primer: 5’-TTTCAAGTTACGGTAAGCATATGATAGTCCATTTTAAAACATAATTTTAAAACTGCAAACTACCCAAGAAA - 3’

#### CRISPR screen data analysis

Sequencing reads were mapped to the sgRNA library using the read_count_CRISPR_guides.pl script available in the GitHub repository (https://github.com/whitehead/barc/tree/main/Perl). The differential sgRNA abundance between DFMO-treated and untreated samples was identified using the Robust Rank Aggregation (RRA) algorithm implemented in the “test” command of MAGeCK (v0.5.9.3)^27^.

#### Cell viability assays

K562, NOMO, HEL, and MEL cells were seeded in 6-well plates at defined densities and pre-treated with sardomozide. For 72-hour pre-treatment, 1 × 10^5^ cells per well were plated in 1 mL of complete growth medium. For extended 96-hour pre-treatment, K562 cells were seeded at 2 × 10^5^ cells per well in 1 mL of medium and passaged into fresh sardomozide-containing medium after 48 hours. Following completion of the pre-treatment period, cells were transferred to medium supplemented with either ML162 or RSL3 and incubated for an additional 24 to 48 hours prior to downstream viability assessment. For cumulative growth curves, cells were passaged every 3 days over 12 days. For iron-free culture, cells were washed twice with pre-warmed HBSS and replated in iron-free IMDM (Gibco).

For viability measurements in Figure S1, the CellTiter-Glo Luminescent Cell Viability Assay (Promega) was used according to the manufacturer’s protocol. Luminescence was recorded on a Tecan Infinite 200 PRO plate reader and normalized to the corresponding AAVS1 control condition. For cell death measurements, Sytox Green was used as a fluorescent marker of cell death. Cells were seeded in the presence of 250 nM sytox green and the average integrated sytox green intensity per cell was measured on an Incucyte S3 instrument. Signal intensity was quantified as a measure of cell death. In all viability and cell-death assays, DMSO served as the vehicle control.

#### Lentivirus preparation and transduction

Lentiviral particles were produced by transient transfection of HEK293T cells with 2 µg of lentiviral transfer plasmid, 1 µg of packaging plasmid psPAX2, and 0.5 µg of envelope plasmid pCMV-VSV-G, using 8 µL of Lipofectamine LTX (Invitrogen, 15338-100) in 500 µL Opti-MEM Reduced Serum Medium (Gibco, 31985-070) according to the manufacturer’s instructions. Viral supernatants were harvested 48 hours post-transfection, clarified by filtration through a 0.45-μm filter, and used to transduce target cells in the presence of 10 μg/mL polybrene (Millipore Sigma, TR1003G). Single guide RNA (sgRNA) sequences, Cas9 expression systems, and vector constructs used for CRISPR-Cas9 knockout are detailed in Table S4. Following antibiotic or fluorescence selection, single-cell clones were isolated, expanded, and validated for gene knockout by immunoblot analysis. For GPX4 knockout, single-cell K562 clones were isolated and expanded in complete medium containing liproxstatin-1 to suppress ferroptotic cell death during clone generation. Prior to all functional assays, liproxstatin-1 was removed by washing cells three times with PBS, and cells were replated in liproxstatin-1-free medium for the indicated assay window. Unless otherwise noted, GPX4 KO and AAVS1 control cells were assayed in parallel under the same post-withdrawal conditions. Constructs used for the expression of iron and polyamine sensors are also listed in Table S4.

#### Liproxstatin-1 withdrawal assay

AAVS1 control and GPX4 KO K562 cells were plated in 96-well plates at the indicated densities in 100 μL complete medium per well. GPX4 KO cells were propagated in liproxstatin-1 prior to the assay. Before seeding, liproxstatin-1 was removed by washing cells three times with PBS. Where indicated, liproxstatin-1 was added back at the time of plating. Viability and cell death were measured 72 h after seeding. Viability was measured using CellTiter-Glo (Promega) according to the manufacturer’s protocol, and luminescence was recorded using a Tecan Infinite 200 PRO plate reader. Cell death was measured using Sytox Green (Thermo Fisher Scientific), which was added at 250 nM at the time of seeding, and average integrated Sytox Green intensity per cell was quantified using an Incucyte S3 instrument.

#### Western blot

Cells were washed with ice-cold PBS and lysed in RIPA lysis buffer (25 mM Tris-HCl pH 7.5 (Invitrogen 15567027), 150 mM NaCl (Invitrogen AM9760G), 1% (v/v) NP-40 (Fisher Scientific AAJ19628AP), 1% (w/v) sodium deoxycholate (Sigma-Aldrich D6750), 0.1% (w/v) SDS (Bio-Rad 1610302)) supplemented with 1% (v/v) HALT protease and phosphatase inhibitors (Thermo Scientific 78429) and 125 U/mL Benzonase nuclease (EMD Millipore E1014). Lysates were homogenized and incubated on ice for 30 min with intermittent vortexing every 10 min. Insoluble debris was removed by centrifugation at 21000× g for 10 min at 4°C. Clarified lysates were mixed with 4× Bolt lithium dodecyl sulfate (LDS) sample buffer (Invitrogen B0007) supplemented with 100 mM DTT (Thermo Scientific R0861) and heated at 70°C for 5 min. This boiling step was skipped for membrane proteins (TFRC, FSP1 and SLC40A1) and instead lysate was kept at 37°C for 30 min. Samples were resolved on a Bolt 4-12% Bis-tris polyacrylamide gel (Invitrogen, NW04122) and transferred to PVDF membranes (Invitrogen IB24002) using the iBlot 2 dry blotting system (Invitrogen IB21001).

Membranes were blocked for 1 h at room temperature in 5% (w/v) nonfat dry skim milk (BD Biosciences 232100) prepared in TBST (tris-buffered saline (Fisher Scientific AAJ60764K3) containing 0.1% (v/v) Tween-20 (Fisher Scientific BP337). Primary antibodies were diluted in 5% (w/v) BSA in TBST and incubated overnight at 4°C. After four washes in TBST, membranes were incubated with HRP-conjugated secondary antibodies diluted in 1% (w/v) skim milk in TBST for 1 hour at room temperature. Following four additional TBST washes, chemiluminescent signals were developed using SuperSignal West Femto Maximum Sensitivity Substrate (Thermo Scientific, 34095) and imaged with a ChemiDoc XRS+ system (Bio-Rad). Antibodies were used at the following dilutions: SRM (Proteintech, 19858-1-AP; 1:1000), SMS (Abcam, ab156879; 1:1000), β-actin (Abcam, ab20272; 1:50,000), ACSL4 (Thermo Fisher, PA5-27137; 1:5000), GPX4 (Abcam, ab41787; 1:2000), FSP1 (Cell Signaling Technology, 24972S; 1:1000), Hypusine (Millipore, ABS1064-I; 1:2500), FTH (Abcam, ab75973; 1:2000), TFRC (Invitrogen, 13-6800; 1:5000), SLC40A1 (Novus Biologicals, NBP1-21502; 1:2000), EGFP (Abcam, AB6556; 1:6000), and anti-rabbit HRP-conjugated secondary antibody (Sigma, A0545; 1:5000).

#### BODIPY staining

Cells were pre-treated with sardomozide for 96 h. ML162 was added 2.5 h prior to analysis. 400,000 cells per well were seeded on 6-well dishes for each treatment condition. Cells were incubated with C11-BODIPY (581/591) (5 μM) for 30 min at 37°C. Subsequently, cells were washed once in HBSS (Gibco) and then resuspended in 300 μL of fresh HBSS (Gibco) strained through a 35 μm cell strainer (Falcon tube with cell strainer cap) and analyzed using the FITC and PE-Texas Red filter. At least 10,000 events were analyzed per sample. Data was analyzed using FlowJo Software.

#### FENIX assay

Liposomes were prepared based on methods previously described^39^. Briefly, soy L-α-phosphatidylcholine (PC, Avanti) in chloroform was formed into a thin film using a rotary evaporator (Buchi). The film was hydrated in 3 mL of PBS and sonicated for 3-5 minutes at 65°C to yield a 20 mM lipid solution. The sample then underwent 3 cycles of freeze-thawing followed by extrusion (Avestin Liposofast LF-50) once at 65°C through a 200 nm membrane (Cytiva Nuclepore) then 3X through 100 nm membranes (Cytiva Nuclepore). The lipid suspension was stored at 4°C. The experiment was performed as previously described^51^. In brief, 0.1□mM liposome was mixed with 2□μM STY-BODIPY (Cayman) and 275□µl of the mixture was added to a black 96-well plate, and compounds (putrescine, spermine or PMC) were added to the indicated concentration, and the samples were incubated for 30□min at 37□°C and initiated by 5□µl of 12□mM DTUN (Cayman) in ethanol, and mixed using pipette to start autoxidation. Data were acquired by STY-BODIPY excitation at 488□nm and emission was measured at 518□nm using a microplate reader.

#### RhoNoxM and RhoNox-1 staining

##### RhoNoxM

Cells were pre-treated with sardomozide (72 h, 5 μM), washed twice with HBSS to remove extracellular iron, resuspended in IMDM, and counted. 0.5 ×□10 cells/well were seeded into 6-well plates with 1 mL iron-free IMDM (Gibco); sardomozide was reintroduced. For iron detection, cells were incubated with RhoNox-M^65^ (2 μM, synthesized in-house) for 5 h at 37°C, with ferrous ammonium sulfate added during the final hour. Cells were washed three times with pre-warmed HBSS and resuspended in IMDM. For lysosomal content, cells were stained with LysoTracker (20 nM, Thermo Fisher) for 30 min at 37°C in the dark and washed. Flow cytometry was performed using PE (570 nm, 580/14 nm) and APC (638 nm, 660/20 nm) channels. ≥5,000 events/sample were acquired and analyzed using FlowJo.

##### RhoNox1

U-2OS cells were treated with sardomozide (5 μM, 48 h) and DFO (100 μM, 48 h). Cells were washed three times with pre-warmed HBSS and then incubated with RhoNox-1 (5 μM in IMDM, synthesized in-house) for 4 h at 37°C. Sardomozide and DFO were reintroduced during this incubation. Finally, cells were again washed three times with pre-warmed HBSS and trypsinized. Flow cytometry was performed using PE (570 nm, 580/14 nm) channel. ≥5,000 events/sample were acquired and analyzed using FlowJo.

#### In-vivo studies

Female NSG mice (Jackson: 005557) were injected subcutaneously with 3x10^6^ K562 cells harboring our polyamine reporter construct. One day after injection, select mice were treated continuously with 1% DFMO (Eflornithine hydrochloride hydrate; MedChemExpress: HY-B0744B) in drinking water. Water bottles were exchanged every 5 days throughout the treatment window. Select mice were also treated daily with (3mg/kg) AMXT-1501 tetrahydrochloride (MedChemExpress: HY-124617A) formulated in sterile PBS and delivered intraperitoneally. All other experimental groups received daily injections of PBS as a vehicle control. Mice were treated for 14 days before doxycycline treatment to induce the polyamine sensor. Doxycycline hyclate (2mg/mL) was delivered in drinking water for 48 hours before tumors were harvested for flow cytometry and immunoblot analysis.

#### Flow cytometry for tumors

A small portion of each tumor was minced, and enzymatically dissociated in collagenase-containing RPMI under gentle agitation for 45 minutes at 37°C. Samples were then passed through 70 µm mesh filters, pelleted by centrifugation at 300xg for 5 minutes, and resuspended in PBS before passing through 40um mesh filters. Samples were then pelleted again, supernatant was removed, and 500 µL of 4% PFA was added for fixation of the isolated single cells. Cells were analyzed on a BD LSR II flow cytometer (BD FACSDiva) with the following laser set up: GFP (excitation: 488□nm, 530/30□nm bandpass filter), mCherry (excitation: 561□nm, 610/20□nm bandpass filter). At least 10,000 events were analyzed per sample. Data was analyzed using FlowJo Software.

#### Iron measurements using ICP-MS

Cells were washed twice with HBSS, counted, and pelleted at 4 × 10□cells per experimental condition. Cell pellets were snap-frozen in liquid nitrogen and stored briefly at –80°C. For analysis, pellets were thawed and resuspended in 100 μL of trace metal–free water (VWR, 87003-236), transferred to metal-free perfluoroalkoxy (PFA) tubes, and lyophilized overnight. Dried pellets were resuspended in 100 μL of 67% trace metal–free nitric acid (VWR, 87003-226) and digested at 80°C for 20 hours. Digested samples were diluted with trace metal–free water to a final nitric acid concentration of 2%. Blank samples underwent identical processing. An Fe standard curve (0–100,000 ppb; Agilent, 5183-4688) was prepared, and a terbium internal standard (1 ppb final concentration; Agilent, 5190-8590) was added to all samples and standards. Total Fe was quantified using an Agilent 7900 inductively coupled plasma mass spectrometer (ICP-MS) operated with a helium collision cell to minimize polyatomic interferences. Data acquisition and quantification were performed using Agilent MassHunter Workstation software. Sample Fe concentrations were determined by interpolation from the standard curve, blank-subtracted, and normalized to cell number. Statistical analyses were conducted in GraphPad Prism.

#### Untargeted metabolomics

All tubes were pre-chilled, and steps were performed on ice unless otherwise noted. Cells were pelleted by centrifugation at 4°C, washed once with 1.5 mL of ice-cold 0.9% NaCl, and pelleted again to remove the wash solution. Cell pellets were immediately snap-frozen on dry ice. Cells were resuspended in 1 mL of 80% methanol (pre-chilled to 20°C). Tubes were vortexed for 10 minutes in a cold room. Samples were then dried using a SpeedVac concentrator. Dried extracts were stored at –80°C prior to LC-MS. Polar metabolites were analyzed using a QExactive Orbitrap MS (Thermo) with an Ion Max source and HESI II probe, coupled to a Dionex UltiMate 3000 UHPLC. External calibration was performed weekly. Samples (5 µL) were injected onto a SeQuant ZIC-pHILIC column (2.1 × 150 mm, 5 µm) with a matching guard column. Column and autosampler were maintained at 25°C and 4°C, respectively. Mobile phase A: 20 mM ammonium carbonate, 0.1% ammonium hydroxide; B: acetonitrile. Flow rate: 0.15 mL/min. Gradient: 0–20 min, 80–20% B; 20–20.5 min, 20–80% B; 20.5–28 min, 80% B hold. The MS operated in full-scan, polarity-switching mode (m/z 70–1000), 70,000 resolution, 3.0 kV spray voltage, 275°C capillary, 350°C HESI probe, sheath/aux/sweep gas: 40/15/1. AGC target was 1×10□, max IT 20 ms. For untargeted metabolomics, ddMS² was acquired on pooled samples (Top 10 method, 15/30/45 V stepped collision energies, 17,500 resolution, AGC 2×10□, max IT 100 ms, isolation window 1.0 m/z). Data were processed in Compound Discoverer 3.1 using an in-house mass list; P-values were Benjamini-Hochberg corrected.

#### Thiol metabolomics

Cysteine and GSH levels (Figure S1K) were measured using a modified Ellman labeling protocol^131^. Briefly, 2 × 10□ cells were pelleted, washed with ice-cold 0.9% NaCl, and extracted with 400 µL ice-cold 80% methanol. Internal standard (10 µL of 4.4 µM 13C,15N-cysteine; HY-Y03375) and 440 µL Ellman’s reagent (10 mM in 80% methanol) were added. Samples were incubated on ice for 1 h, centrifuged (3,000 × g, 5 min), and supernatants stored at –80°C until analysis. Samples were run as above with a targeted selected ion monitoring scan (tSIM) in positive mode centred on m/z 319.00530 and m/z 323.01240 to increase signal for the cysteine-TNB (2-nitro-5-thiobenzoic acid) mixed disulfide and the [13C, 15N]cysteine–TNB mixed disulfide, respectively.

#### RNA-sequencing

Total RNA was extracted with PureLink RNA Mini Kit (Invitrogen) per the manufacturer’s instruction. PolyA selection and RNA library construction were performed by Novogene with sequencing as 150 × 150 base paired-end libraries using an Illumina NovaSeq 6000 (Novogene), to a depth of ≥ 20M reads. Reads were aligned to the human genome (hg38, annotation file GRCh GTF 38.93) using the short-read alignment tool STAR (version 2.7.1a)^132^ with the default options. Expression of each mRNA was calculated and reported as fragments per kilobase of transcript sequence per million base pairs sequenced (FPKM). The change in FPKM was compared between groups (e.g., control versus sardomozide) and differential expression was analyzed using the DESeq2 package. FDR correction was performed using the Benjamini-Hochberg method.

For RNA-seq with DFO treatment: Cells were collected by trypsinization and pellets stored at -80 °C until use. Libraries for differential gene expression were prepared by Plasmidsaurus using BRB-seq^133^, which employs barcoded poly-T primers to capture polyadenylated transcripts, and sequenced to a depth of ≥ 20 million reads per sample. Sequenced reads were trimmed using FastP v0.24.0^134^ with a minimum Phred score of 15 and minimum length of 50 base, then aligned to the human genome (hg38) using STAR 2.7.11^132^ with default options except for --outFilterIntronMotifs RemoveNoncanonical.

Reads were deduplicated using UMICollapse v1.1.0^135^, yielding 14.4-17.6 million unique reads per sample. Gene-level counts were generated using featureCounts 2.0.1^136^ with the GRCh38 114 annotation file. Differential expression analysis was performed using edgeR^137^, with a false discovery rate threshold of 0.05 and log2 fold-change threshold of 1.

For tumors: Tumor samples were processed as above, except reads were aligned to both the human (hg38) and mouse (mm39) genomes. Contaminating mouse reads (3.4-4.9%) were removed with XenofilteR (1.6), yielding 17.3-20.2M human reads per sample.

#### Proteomics

U-2OS cells grown in 6-well plates were harvested at approximately 50% confluency. Culture medium was aspirated, and cells were washed three times with 3 mL PBS. Cells were lysed directly in the wells using 300 μL lysis buffer (50 mM triethylammonium bicarbonate [TEAB], 5% SDS, 2 mM MgCl□) supplemented with benzonase (1 U per mL lysis buffer). Lysates were incubated for 15 min at room temperature with gentle agitation, transferred to low-binding tubes, and clarified by centrifugation at maximum speed for 10 min. The supernatant was transferred to fresh tubes and total protein concentration was determined by BCA assay. Equal amounts of protein (typically 30 μg per sample) were aliquoted, adjusted to equal volumes with lysis buffer, snap-frozen in liquid nitrogen, and stored at −80 °C until submission for mass spectrometry. Sample integrity and loading were verified by SDS-PAGE prior to submission to Whitehead Proteomics Core Facility.

#### Microscopy

Cells were seeded on glass-bottom 96-well plates (Brooks, MGB096-1-2-LG-L) and imaged on a Dragonfly 505 spinning-disk confocal microscope (Andor Technologies) equipped with a piezo Z-stage (ASI) and an iXon Ultra 888 EMCCD camera. Imaging was performed using a 60× oil immersion objective with a pinhole size of 40 μm. Z-stacks were acquired with a step size of 0.3–1 μm, depending on sample thickness (pixel size 121 nm × 121 nm). Cells were maintained at 37 °C and 5% CO□ in a humidified chamber (OKO Labs) during imaging. The following excitation and emission settings were used: YFP (488 nm, 521/38 nm), mCherry (561 nm, 594/43 nm), and miRFP670-2 (637 nm, 620/60 nm). For each condition, at least 40 randomly selected cells were imaged across at least two independent experiments. Representative images were generated as average projections of three z-slices centered on the focal plane.

#### Enzymatic polyamine assay

Measurements were performed using the fluorescent quantification assays (abcam239728) as per the manufacturer’s instructions. The polyamine assay buffer (no added polyamine standard) was used as the background.

#### C8-pos lipidomics

Cell pellets (4 × 10□) were reconstituted in 1 mL isopropanol containing 1,2-didodecanoyl-sn-glycero-3-phosphocholine (Avanti) as internal standard, vortexed, and centrifuged (10,000 × g, 10 min).

Supernatants (10 µL) were injected onto a 100 × 2.1 mm, 1.7 µm ACQUITY BEH C8 column (Waters). Lipidomic profiling was performed on a Shimadzu Nexera X2 U-HPLC coupled to an Exactive Plus Orbitrap MS (Thermo), operated in positive ion mode (full scan, m/z 220–1100, 70,000 resolution, 3 Hz acquisition). MS parameters: sheath gas 50, sweep gas 5, in-source CID 5 eV, spray voltage 3 kV, capillary/heater temperatures 300°C, S-lens RF 60, AGC target 1 × 10□, max ion time 100 ms, 1 microscan. The column was eluted with 80% mobile phase A (95:5:0.1, 10 mM ammonium acetate/methanol/formic acid) for 1 min, ramped to 80% mobile phase B (99.9:0.1 methanol/formic acid) over 2 min, then to 100% B over 7 min, and held at 100% B for 3 min. Raw data were processed using TraceFinder software (Thermo Fisher Scientific) for targeted peak integration and manual review of a subset of identified lipids and using Progenesis QI (Nonlinear Dynamics) for peak detection and integration of both lipids of known identify and unknowns. Lipid identities were determined based on comparison to reference plasma extracts and are denoted by total number of carbons in the lipid acyl chain(s) and total number of double bonds in the lipid acyl chain(s).

For lipidomic analysis, only lipid species annotated by Progenesis QI were included; unannotated features were excluded from downstream analysis. To account for differences in total lipid signal arising from variation in sample input or extraction efficiency, the intensity of each annotated lipid species within a sample was normalized to the summed intensity of all annotated lipid species in that sample. Lipid subclass abundances such as O-PC, TG, and others were calculated by summing normalized intensities of species assigned to the same subclass. Within each lipid class such as PC or PE, abundances were further aggregated based on the number of double bonds to assess changes in unsaturation. Because normalization to annotated species can be influenced by incomplete lipid annotation, we additionally analyzed changes at the level of individual lipid species using Z-score normalization across samples. These analyses yielded trends consistent with those observed after abundance normalization, supporting the robustness of the results.

#### Mössbauer spectroscopy

All samples were prepared under oxygen-free conditions in an anaerobic glovebox (Coylab). Unless otherwise indicated, samples were prepared in 50 mM HEPES, pH 7.5. For spermine titrations, 1 mM ^57^Fe^2+^ was combined with 20, 40, or 60 mM spermine; control samples contained 1 mM ^57^Fe^2+^ with 2 mM DFO or 2 mM ATP. All spectra were recorded at 80 K in zero applied magnetic field. For ATP competition experiments, 1 mM ^57^Fe^2+^ was analyzed alone, with 1 mM ATP, or with 1 mM ATP plus 15 mM spermine in 50 mM HEPES, pH 7.5. For putrescine/spermine comparisons at pH 7.5, 1 mM ^57^Fe^2+^ was combined with 40 mM putrescine or spermine in 200 mM HEPES, pH 7.5. For alkaline comparisons, 1 mM ^57^Fe^2+^ was combined with 40 mM putrescine or spermine in 50 mM CHES, pH 10. Mössbauer spectra were collected using a WEB Research spectrometer (Edina, MN) equipped with a Janis SVT-400 variable-temperature cryostat. Isomer shifts are reported relative to the centroid of an α-iron metal spectrum recorded at room temperature. Spectra were fitted and analyzed using WMOSS software (WEB Research).

#### Ferrozine competition assay

All solutions were prepared in 50 mM HEPES (pH 7.5) under oxygen-free conditions in an anaerobic glovebox (Coylab). Fe^2+^ was supplied as (NH_4_)_2_ FeSO_4_ (200 μM). Spermine was added at the indicated concentrations and allowed to equilibrate for 5 min, after which ferrozine was added to a final concentration of 15 μM. Absorbance spectra were recorded using a Cary UV–Vis spectrophotometer (Agilent), and formation of the Fe^2+^–ferrozine complex was monitored at 562 nm.

#### FerroOrange assay

All buffers and salts were prepared aerobically in argon-degassed Milli-Q water and adjusted to the desired pH with NaOH. The indicated polyamine was mixed with FerroOrange-containing buffer, and Fe^2+^ was added last to minimize oxidation. Samples were incubated for 30 min in the dark, and fluorescence was measured with excitation at 543 nm and emission at 580 nm. Reactions contained 200 mM HEPES (pH 7.5), 50 μM FeCl_2_, and 10 μM FerroOrange, with spermidine or spermine added at 5–100 mM (100–2000-fold molar excess over Fe^2+^). DFO (1 mM) was used as a positive control.

#### Quantification and statistical analysis

Data are presented as mean ± SD unless otherwise specified. For dot plots, box boundaries indicate the interquartile range, and the line indicates the median. Unless otherwise noted in a specific legend, statistical significance was assessed by two-tailed Student’s t-test for pairwise comparisons; the specific test used for any panel employing a different method (e.g., ANOVA, nonparametric tests) is stated in that figure’s legend. A p value < 0.05 was considered statistically significant, and exact p values are reported in the figures where applicable. Unless otherwise noted, graphs were generated using GraphPad Prism.

## Supplemental table legends

**Table S1. Genome-wide CRISPR-Cas9 screen results under polyamine depletion, related to Figure 1.**

**Table S2. C8-positive lipidomics dataset, related to Figures 3 and 4.**

**Table S3. Tumor versus normal expression of ODC1 and SRM across cancer types, related to Figure 7 and Discussion.**

**Table S4. Plasmid and construct sequences used in this study, related to STAR Methods.**

## Notes

### Competing Interest Statement

A.J. and P.S. are inventors on a pending patent application based on this work. The remaining authors declare no competing financial interests.

### Summary of Updates

Minor changes to the text and figures for clarity throughout; no major changes to data or conclusions.

## References

1. Pegg, A.E. (2016). Functions of Polyamines in Mammals*. J. Biol. Chem. 291, 14904–14912. 10.1074/jbc.R116.731661.

2. Watanabe, S., Kusama-Eguchi, K., Kobayashi, H., and Igarashi, K. (1991). Estimation of polyamine binding to macromolecules and ATP in bovine lymphocytes and rat liver. J. Biol. Chem. 266, 20803–20809.

3. Matsufuji, S., Matsufuji, T., Miyazaki, Y., Murakami, Y., Atkins, J.F., Gesteland, R.F., and Hayashi, S. (1995). Autoregulatory frameshifting in decoding mammalian ornithine decarboxylase antizyme. Cell 80, 51–60. 10.1016/0092-8674(95)90450-6.

4. Rom, E., and Kahana, C. (1994). Polyamines regulate the expression of ornithine decarboxylase antizyme in vitro by inducing ribosomal frame-shifting. Proc. Natl. Acad. Sci. U. S. A. 91, 3959–3963. 10.1073/pnas.91.9.3959.

5. Pegg, A.E. (2008). Spermidine/spermine-N(1)-acetyltransferase: a key metabolic regulator. Am. J. Physiol. Endocrinol. Metab. 294, E995–1010. 10.1152/ajpendo.90217.2008.

6. Mitchell, J.L., Judd, G.G., Bareyal-Leyser, A., and Ling, S.Y. (1994). Feedback repression of polyamine transport is mediated by antizyme in mammalian tissue-culture cells. Biochem. J. 299 ( *Pt 1*), 19–22. 10.1042/bj2990019.

7. Park, M.H., and Wolff, E.C. (2018). Hypusine, a polyamine-derived amino acid critical for eukaryotic translation. J. Biol. Chem. 293, 18710–18718. 10.1074/jbc.TM118.003341.

8. Chattopadhyay, M.K., Park, M.H., and Tabor, H. (2008). Hypusine modification for growth is the major function of spermidine in Saccharomyces cerevisiae polyamine auxotrophs grown in limiting spermidine. Proc. Natl. Acad. Sci. U. S. A. 105, 6554–6559. 10.1073/pnas.0710970105.

9. Cho, N.H., Cheveralls, K.C., Brunner, A.-D., Kim, K., Michaelis, A.C., Raghavan, P., Kobayashi, H., Savy, L., Li, J.Y., Canaj, H., et al. (2022). OpenCell: Endogenous tagging for the cartography of human cellular organization. Science 375, eabi6983. 10.1126/science.abi6983.

10. Park, S., Athreya, A., Carrizo, G.E., Benning, N.A., Mitchener, M.M., Bhanu, N.V., Garcia, B.A., Zhang, B., Muir, T.W., Pearce, E.L., et al. (2023). Electrostatic encoding of genome organization principles within single native nucleosomes. BioRxiv Prepr. Serv. Biol., 2023.12.08.570828. 10.1101/2023.12.08.570828.

11. Zhang, D., Zhao, T., Ang, H.S., Chong, P., Saiki, R., Igarashi, K., Yang, H., and Vardy, L.A. (2012). AMD1 is essential for ESC self-renewal and is translationally down-regulated on differentiation to neural precursor cells. Genes Dev. 26, 461–473. 10.1101/gad.182998.111.

12. Minetti, A., Omrani, O., Brenner, C., Allies, G., Imada, S., Rösler, J., Khawaled, S., Cansiz, F., Meckelmann, S.W., Gebert, N., et al. (2024). Polyamines sustain epithelial regeneration in aged intestines by modulating protein homeostasis. Preprint at bioRxiv, 10.1101/2024.07.26.605278 https://doi.org/10.1101/2024.07.26.605278.

13. Hofer, S.J., Simon, A.K., Bergmann, M., Eisenberg, T., Kroemer, G., and Madeo, F. (2022). Mechanisms of spermidine-induced autophagy and geroprotection. Nat. Aging 2, 1112–1129. 10.1038/s43587-022-00322-9.

14. Puleston, D.J., Baixauli, F., Sanin, D.E., Edwards-Hicks, J., Villa, M., Kabat, A.M., Kamiński, M.M., Stanckzak, M., Weiss, H.J., Grzes, K.M., et al. (2021). Polyamine metabolism is a central determinant of helper T cell lineage fidelity. Cell 184, 4186–4202.e20. 10.1016/j.cell.2021.06.007.

15. Gupta, V.K., Scheunemann, L., Eisenberg, T., Mertel, S., Bhukel, A., Koemans, T.S., Kramer, J.M., Liu, K.S.Y., Schroeder, S., Stunnenberg, H.G., et al. (2013). Restoring polyamines protects from age-induced memory impairment in an autophagy-dependent manner. Nat. Neurosci. 16, 1453–1460. 10.1038/nn.3512.

16. Eisenberg, T., Abdellatif, M., Schroeder, S., Primessnig, U., Stekovic, S., Pendl, T., Harger, A., Schipke, J., Zimmermann, A., Schmidt, A., et al. (2016). Cardioprotection and lifespan extension by the natural polyamine spermidine. Nat. Med. 22, 1428–1438. 10.1038/nm.4222.

17. Eisenberg, T., Knauer, H., Schauer, A., Büttner, S., Ruckenstuhl, C., Carmona-Gutierrez, D., Ring, J., Schroeder, S., Magnes, C., Antonacci, L., et al. (2009). Induction of autophagy by spermidine promotes longevity. Nat. Cell Biol. 11, 1305–1314. 10.1038/ncb1975.

18. van Veen, S., Martin, S., Van den Haute, C., Benoy, V., Lyons, J., Vanhoutte, R., Kahler, J.P., Decuypere, J.-P., Gelders, G., Lambie, E., et al. (2020). ATP13A2 deficiency disrupts lysosomal polyamine export. Nature 578, 419–424. 10.1038/s41586-020-1968-7.

19. Schwartz, C.E., Wang, X., Stevenson, R.E., and Pegg, A.E. (2011). Spermine synthase deficiency resulting in X-linked intellectual disability (Snyder-Robinson syndrome). Methods Mol. Biol. Clifton NJ 720, 437–445. 10.1007/978-1-61779-034-8_28.

20. Bachmann, A.S., VanSickle, E.A., Michael, J., Vipond, M., and Bupp, C.P. (2024). Bachmann-Bupp syndrome and treatment. Dev. Med. Child Neurol. 66, 445–455. 10.1111/dmcn.15687.

21. Casero, R.A., Murray Stewart, T., and Pegg, A.E. (2018). Polyamine metabolism and cancer: treatments, challenges and opportunities. Nat. Rev. Cancer 18, 681–695. 10.1038/s41568-018-0050-3.

22. Shalem, O., Sanjana, N.E., Hartenian, E., Shi, X., Scott, D.A., Mikkelson, T., Heckl, D., Ebert, B.L., Root, D.E., Doench, J.G., et al. (2014). Genome-scale CRISPR-Cas9 knockout screening in human cells. Science 343, 84–87. 10.1126/science.1247005.

23. Wang, T., Wei, J.J., Sabatini, D.M., and Lander, E.S. (2014). Genetic screens in human cells using the CRISPR-Cas9 system. Science 343, 80–84. 10.1126/science.1246981.

24. Yang, W.S., SriRamaratnam, R., Welsch, M.E., Shimada, K., Skouta, R., Viswanathan, V.S., Cheah, J.H., Clemons, P.A., Shamji, A.F., Clish, C.B., et al. (2014). Regulation of Ferroptotic Cancer Cell Death by GPX4. Cell 156, 317–331. 10.1016/j.cell.2013.12.010.

25. Li, Z., Lange, M., Dixon, S.J., and Olzmann, J.A. (2024). Lipid Quality Control and Ferroptosis: From Concept to Mechanism. Annu. Rev. Biochem. 93, 499–528. 10.1146/annurev-biochem-052521-033527.

26. Inglis, A.J., Guna, A., Gálvez-Merchán, Á., Pal, A., Esantsi, T.K., Keys, H.R., Frenkel, E.M., Oania, R., Weissman, J.S., and Voorhees, R.M. (2023). Coupled protein quality control during nonsense-mediated mRNA decay. J. Cell Sci. 136, jcs261216. 10.1242/jcs.261216.

27. Li, W., Xu, H., Xiao, T., Cong, L., Love, M.I., Zhang, F., Irizarry, R.A., Liu, J.S., Brown, M., and Liu, X.S. (2014). MAGeCK enables robust identification of essential genes from genome-scale CRISPR/Cas9 knockout screens. Genome Biol. 15, 554. 10.1186/s13059-014-0554-4.

28. Uemura, T., Yerushalmi, H.F., Tsaprailis, G., Stringer, D.E., Pastorian, K.E., Hawel, L., Byus, C.V., and Gerner, E.W. (2008). Identification and Characterization of a Diamine Exporter in Colon Epithelial Cells. J. Biol. Chem. 283, 26428–26435. 10.1074/jbc.M804714200.

29. Goldmann, U., and Sedlyarov, V. (2019). RESOLUTE database.

30. Dixon, S.J., Lemberg, K.M., Lamprecht, M.R., Skouta, R., Zaitsev, E.M., Gleason, C.E., Patel, D.N., Bauer, A.J., Cantley, A.M., Yang, W.S., et al. (2012). Ferroptosis: An Iron-Dependent Form of Non-Apoptotic Cell Death. Cell 149, 1060–1072. 10.1016/j.cell.2012.03.042.

31. Hamouda, N.N., Van den Haute, C., Vanhoutte, R., Sannerud, R., Azfar, M., Mayer, R., Cortés Calabuig, Á., Swinnen, J.V., Agostinis, P., Baekelandt, V., et al. (2021). ATP13A3 is a major component of the enigmatic mammalian polyamine transport system. J. Biol. Chem. 296, 100182. 10.1074/jbc.RA120.013908.

32. Vander Heiden, M.G., Li, X.X., Gottleib, E., Hill, R.B., Thompson, C.B., and Colombini, M. (2001). Bcl-xL promotes the open configuration of the voltage-dependent anion channel and metabolite passage through the outer mitochondrial membrane. J. Biol. Chem. 276, 19414–19419. 10.1074/jbc.M101590200.

33. Wang, L., Doherty, G.A., Judd, A.S., Tao, Z.-F., Hansen, T.M., Frey, R.R., Song, X., Bruncko, M., Kunzer, A.R., Wang, X., et al. (2020). Discovery of A-1331852, a First-in-Class, Potent, and Orally-Bioavailable BCL-XL Inhibitor. ACS Med. Chem. Lett. 11, 1829–1836. 10.1021/acsmedchemlett.9b00568.

34. Doll, S., Proneth, B., Tyurina, Y.Y., Panzilius, E., Kobayashi, S., Ingold, I., Irmler, M., Beckers, J., Aichler, M., Walch, A., et al. (2017). ACSL4 dictates ferroptosis sensitivity by shaping cellular lipid composition. Nat. Chem. Biol. 13, 91–98. 10.1038/nchembio.2239.

35. Bi, G., Liang, J., Bian, Y., Shan, G., Huang, Y., Lu, T., Zhang, H., Jin, X., Chen, Z., Zhao, M., et al. (2024). Polyamine-mediated ferroptosis amplification acts as a targetable vulnerability in cancer. Nat. Commun. 15, 2461. 10.1038/s41467-024-46776-w.

36. Murray Stewart, T., Dunston, T.T., Woster, P.M., and Casero, R.A. (2018). Polyamine catabolism and oxidative damage. J. Biol. Chem. 293, 18736–18745. 10.1074/jbc.TM118.003337.

37. Zilka, O., Shah, R., Li, B., Friedmann Angeli, J.P., Griesser, M., Conrad, M., and Pratt, D.A. (2017). On the Mechanism of Cytoprotection by Ferrostatin-1 and Liproxstatin-1 and the Role of Lipid Peroxidation in Ferroptotic Cell Death. ACS Cent. Sci. 3, 232–243. 10.1021/acscentsci.7b00028.

38. Cañeque, T., Baron, L., Müller, S., Carmona, A., Colombeau, L., Versini, A., Solier, S., Gaillet, C., Sindikubwabo, F., Sampaio, J.L., et al. (2025). Activation of lysosomal iron triggers ferroptosis in cancer. Nature 642, 492–500. 10.1038/s41586-025-08974-4.

39. Li, Y., Ran, Q., Duan, Q., Jin, J., Wang, Y., Yu, L., Wang, C., Zhu, Z., Chen, X., Weng, L., et al. (2024). 7-Dehydrocholesterol dictates ferroptosis sensitivity. Nature 626, 411–418. 10.1038/s41586-023-06983-9.

40. Tsvetkov, P., Coy, S., Petrova, B., Dreishpoon, M., Verma, A., Abdusamad, M., Rossen, J., Joesch-Cohen, L., Humeidi, R., Spangler, R.D., et al. (2022). Copper induces cell death by targeting lipoylated TCA cycle proteins. Science 375, 1254–1261. 10.1126/science.abf0529.

41. Alvarez, S.W., Sviderskiy, V.O., Terzi, E.M., Papagiannakopoulos, T., Moreira, A.L., Adams, S., Sabatini, D.M., Birsoy, K., and Possemato, R. (2017). NFS1 undergoes positive selection in lung tumours and protects cells from ferroptosis. Nature 551, 639–643. 10.1038/nature24637.

42. Tada, K., Nishizawa, H., Shima, H., Muto, A., Wada, M., and Igarashi, K. (2024). Cysteine restriction induces ferroptosis depending on the polyamine biosynthetic pathway in hepatic cancer cells. Preprint at bioRxiv, 10.1101/2024.02.29.582667 https://doi.org/10.1101/2024.02.29.582667.

43. Sanayama, H., Ito, K., Ookawara, S., Uemura, T., Sakiyama, Y., Sugawara, H., Tabei, K., Igarashi, K., and Soda, K. (2023). Whole Blood Spermine/Spermidine Ratio as a New Indicator of Sarcopenia Status in Older Adults. Biomedicines 11, 1403. 10.3390/biomedicines11051403.

44. Schuller, A.P., Wu, C.C.-C., Dever, T.E., Buskirk, A.R., and Green, R. (2017). eIF5A Functions Globally in Translation Elongation and Termination. Mol. Cell 66, 194–205.e5. 10.1016/j.molcel.2017.03.003.

45. Manjunath, H., Zhang, H., Rehfeld, F., Han, J., Chang, T.-C., and Mendell, J.T. (2019). Suppression of Ribosomal Pausing by eIF5A Is Necessary to Maintain the Fidelity of Start Codon Selection. Cell Rep. 29, 3134–3146.e6. 10.1016/j.celrep.2019.10.129.

46. Ray, R.M., Zimmerman, B.J., McCormack, S.A., Patel, T.B., and Johnson, L.R. (1999). Polyamine depletion arrests cell cycle and induces inhibitors p21(Waf1/Cip1), p27(Kip1), and p53 in IEC-6 cells. Am. J. Physiol. 276, C684–691. 10.1152/ajpcell.1999.276.3.C684.

47. Rodencal, J., Kim, N., He, A., Li, V.L., Lange, M., He, J., Tarangelo, A., Schafer, Z.T., Olzmann, J.A., Long, J.Z., et al. (2024). Sensitization of cancer cells to ferroptosis coincident with cell cycle arrest. Cell Chem. Biol. 31, 234–248.e13. 10.1016/j.chembiol.2023.10.011.

48. Doll, S., Freitas, F.P., Shah, R., Aldrovandi, M., da Silva, M.C., Ingold, I., Goya Grocin, A., Xavier da Silva, T.N., Panzilius, E., Scheel, C.H., et al. (2019). FSP1 is a glutathione-independent ferroptosis suppressor. Nature 575, 693–698. 10.1038/s41586-019-1707-0.

49. Bersuker, K., Hendricks, J.M., Li, Z., Magtanong, L., Ford, B., Tang, P.H., Roberts, M.A., Tong, B., Maimone, T.J., Zoncu, R., et al. (2019). The CoQ oxidoreductase FSP1 acts parallel to GPX4 to inhibit ferroptosis. Nature 575, 688–692. 10.1038/s41586-019-1705-2.

50. Bellé, N.A.V., Dalmolin, G.D., Fonini, G., Rubin, M.A., and Rocha, J.B.T. (2004). Polyamines reduces lipid peroxidation induced by different pro-oxidant agents. Brain Res. 1008, 245–251. 10.1016/j.brainres.2004.02.036.

51. Shah, R., Farmer, L.A., Zilka, O., Van Kessel, A.T.M., and Pratt, D.A. (2019). Beyond DPPH: Use of Fluorescence-Enabled Inhibited Autoxidation to Predict Oxidative Cell Death Rescue. Cell Chem. Biol. 26, 1594–1607.e7. 10.1016/j.chembiol.2019.09.007.

52. Tie, J.-K., Jin, D.-Y., Straight, D.L., and Stafford, D.W. (2011). Functional study of the vitamin K cycle in mammalian cells. Blood 117, 2967–2974. 10.1182/blood-2010-08-304303.

53. Mishima, E., Ito, J., Wu, Z., Nakamura, T., Wahida, A., Doll, S., Tonnus, W., Nepachalovich, P., Eggenhofer, E., Aldrovandi, M., et al. (2022). A non-canonical vitamin K cycle is a potent ferroptosis suppressor. Nature 608, 778–783. 10.1038/s41586-022-05022-3.

54. TeSlaa, T., Ralser, M., Fan, J., and Rabinowitz, J.D. (2023). The pentose phosphate pathway in health and disease. Nat. Metab. 5, 1275–1289. 10.1038/s42255-023-00863-2.

55. Plays, M., Müller, S., and Rodriguez, R. (2021). Chemistry and biology of ferritin. Met. Integr. Biometal Sci. 13, mfab021. 10.1093/mtomcs/mfab021.

56. Rodriguez, R., Müller, S., Colombeau, L., Solier, S., Sindikubwabo, F., and Cañeque, T. (2025). Metal Ion Signaling in Biomedicine. Chem. Rev. 125, 660–744. 10.1021/acs.chemrev.4c00577.

57. Galy, B., Conrad, M., and Muckenthaler, M. (2023). Mechanisms controlling cellular and systemic iron homeostasis. Nat. Rev. Mol. Cell Biol., 1–23. 10.1038/s41580-023-00648-1.

58. Brawley, H.N., Kreinbrink, A.C., Hierholzer, J.D., Vali, S.W., and Lindahl, P.A. (2023). Labile Iron Pool of Isolated Escherichia coli Cytosol Likely Includes Fe-ATP and Fe-Citrate but not Fe-Glutathione or Aqueous Fe. J. Am. Chem. Soc. 145, 2104–2117. 10.1021/jacs.2c06625.

59. Cancer Genome Atlas Network (2012). Comprehensive molecular characterization of human colon and rectal cancer. Nature 487, 330–337. 10.1038/nature11252.

60. Goto, N., Westcott, P.M.K., Goto, S., Imada, S., Taylor, M.S., Eng, G., Braverman, J., Deshpande, V., Jacks, T., Agudo, J., et al. (2024). SOX17 enables immune evasion of early colorectal adenomas and cancers. Nature 627, 636–645. 10.1038/s41586-024-07135-3.

61. Tsherniak, A., Vazquez, F., Montgomery, P.G., Weir, B.A., Kryukov, G., Cowley, G.S., Gill, S., Harrington, W.F., Pantel, S., Krill-Burger, J.M., et al. (2017). Defining a Cancer Dependency Map. Cell 170, 564–576.e16. 10.1016/j.cell.2017.06.010.

62. Sharma, P., Kim, C.Y., Keys, H.R., Imada, S., Joseph, A.B., Ferro, L., Kunchok, T., Anderson, R., Yilmaz, O., Weng, J.-K., et al. (2024). Genetically encoded fluorescent reporter for polyamines. BioRxiv Prepr. Serv. Biol., 2024.08.24.609500. 10.1101/2024.08.24.609500.

63. Gamble, L.D., Purgato, S., Murray, J., Xiao, L., Yu, D.M.T., Hanssen, K.M., Giorgi, F.M., Carter, D.R., Gifford, A.J., Valli, E., et al. (2019). Inhibition of polyamine synthesis and uptake reduces tumor progression and prolongs survival in mouse models of neuroblastoma. Sci. Transl. Med. 11, eaau1099. 10.1126/scitranslmed.aau1099.

64. Hirayama, T., Okuda, K., and Nagasawa, H. (2013). A highly selective turn-on fluorescent probe for iron(II) to visualize labile iron in living cells. Chem. Sci. 4, 1250–1256. 10.1039/C2SC21649C.

65. Niwa, M., Hirayama, T., Okuda, K., and Nagasawa, H. (2014). A new class of high-contrast Fe(II) selective fluorescent probes based on spirocyclized scaffolds for visualization of intracellular labile iron delivered by transferrin. Org. Biomol. Chem. 12, 6590–6597. 10.1039/C4OB00935E.

66. Wang, S., Guo, W., Wan, B.-W., Wang, K.-M., Dong, B., Jiang, C.-S., and Zhang, J. (2026). Recent advances in Golgi-targeted fluorescent probes in ferroptosis research. Analyst. 10.1039/D5AN01240F.

67. Hentze, M.W., Muckenthaler, M.U., and Andrews, N.C. (2004). Balancing acts: molecular control of mammalian iron metabolism. Cell 117, 285–297. 10.1016/s0092-8674(04)00343-5.

68. Eisenstein, R.S., and Blemings, K.P. (1998). Iron regulatory proteins, iron responsive elements and iron homeostasis. J. Nutr. 128, 2295–2298. 10.1093/jn/128.12.2295.

69. Connell, G.J., Abasiri, I.M., and Chaney, E.H. (2023). A temporal difference in the stabilization of two mRNAs with a 3’ iron-responsive element during iron deficiency. RNA N. Y. N 29, 1117–1125. 10.1261/rna.079665.123.

70. Ueda, H., Suzuki, M., Kuroda, R., Tanaka, T., and Aoki, S. (2021). Design, Synthesis, and Biological Evaluation of Boron-Containing Macrocyclic Polyamines and Their Zinc(II) Complexes for Boron Neutron Capture Therapy. J. Med. Chem. 64, 8523–8544. 10.1021/acs.jmedchem.1c00445.

71. 24.3: Basicity of Amines (2015). Chem. Libr. https://chem.libretexts.org/Bookshelves/Organic_Chemistry/Organic_Chemistry_(Morsch_et_al.)/24%3A_Amines_and_Heterocycles/24.03%3A_Basicity_of_Amines.

72. Kontoghiorghes, G.J., and Kontoghiorghe, C.N. (2020). Iron and Chelation in Biochemistry and Medicine: New Approaches to Controlling Iron Metabolism and Treating Related Diseases. Cells 9, 1456. 10.3390/cells9061456.

73. Zhang, Y., Zhao, X., Qin, Y., Li, X., Chang, Y., Shi, Z., Song, M., Sun, W., Xiao, J., Li, Z., et al. (2023). Order-order assembly transition-driven polyamines detection based on iron-sulfur complexes. Commun. Chem. 6, 146. 10.1038/s42004-023-00942-1.

74. Grubwieser, P., Brigo, N., Seifert, M., Grander, M., Theurl, I., Nairz, M., Weiss, G., and Pfeifhofer-Obermair, C. (2024). Quantification of Macrophage Cellular Ferrous Iron (Fe2+) Content Using a Highly Specific Fluorescent Probe in a Plate Reader. Bio-Protoc. 14, e4929. 10.21769/BioProtoc.4929.

75. Gütlich, P., Bill, E., and Trautwein, A.X. (2010). Mössbauer Spectroscopy and Transition Metal Chemistry: Fundamentals and Applications (Springer Science & Business Media).

76. Greenwood, N.N. (2012). Mössbauer Spectroscopy (Springer Science & Business Media).

77. Boggs, R., and Donohue, J. (1975). Spermine copper(II) perchlorate. Acta Crystallogr. B 31, 320–322. 10.1107/S0567740875002658.

78. Tang, Z., Li, C., Kang, B., Gao, G., Li, C., and Zhang, Z. (2017). GEPIA: a web server for cancer and normal gene expression profiling and interactive analyses. Nucleic Acids Res. 45, W98–W102. 10.1093/nar/gkx247.

79. Bello-Fernandez, C., Packham, G., and Cleveland, J.L. (1993). The ornithine decarboxylase gene is a transcriptional target of c-Myc. Proc. Natl. Acad. Sci. U. S. A. 90, 7804–7808. 10.1073/pnas.90.16.7804.

80. Origanti, S., and Shantz, L.M. (2007). Ras transformation of RIE-1 cells activates cap-independent translation of ornithine decarboxylase: regulation by the Raf/MEK/ERK and phosphatidylinositol 3-kinase pathways. Cancer Res. 67, 4834–4842. 10.1158/0008-5472.CAN-06-4627.

81. Roy, U.K.B., Rial, N.S., Kachel, K.L., and Gerner, E.W. (2008). Activated K-RAS increases polyamine uptake in human colon cancer cells through modulation of caveolar endocytosis. Mol. Carcinog. 47, 538–553. 10.1002/mc.20414.

82. Kucharzewska, P., Welch, J.E., Svensson, K.J., and Belting, M. (2009). The polyamines regulate endothelial cell survival during hypoxic stress through PI3K/AKT and MCL-1. Biochem. Biophys. Res. Commun. 380, 413–418. 10.1016/j.bbrc.2009.01.097.

83. Zabala-Letona, A., Arruabarrena-Aristorena, A., Martín-Martín, N., Fernandez-Ruiz, S., Sutherland, J.D., Clasquin, M., Tomas-Cortazar, J., Jimenez, J., Torres, I., Quang, P., et al. (2017). mTORC1-dependent AMD1 regulation sustains polyamine metabolism in prostate cancer. Nature 547, 109– 113. 10.1038/nature22964.

84. Hogarty, M.D., Norris, M.D., Davis, K., Liu, X., Evageliou, N.F., Hayes, C.S., Pawel, B., Guo, R., Zhao, H., Sekyere, E., et al. (2008). ODC1 is a critical determinant of MYCN oncogenesis and a therapeutic target in neuroblastoma. Cancer Res. 68, 9735–9745. 10.1158/0008-5472.CAN-07-6866.

85. Rounbehler, R.J., Li, W., Hall, M.A., Yang, C., Fallahi, M., and Cleveland, J.L. (2009). Targeting ornithine decarboxylase impairs development of MYCN-amplified neuroblastoma. Cancer Res. 69, 547–553. 10.1158/0008-5472.CAN-08-2968.

86. Manni, A., Washington, S., Hu, X., Griffith, J.W., Bruggeman, R., Demers, L.M., Mauger, D., and Verderame, M.F. (2005). Effects of polyamine synthesis inhibitors on primary tumor features and metastatic capacity of human breast cancer cells. Clin. Exp. Metastasis 22, 255–263. 10.1007/s10585-005-8480-1.

87. Klein, S., Miret, J.J., Algranati, I.D., and de Lustig, E.S. (1985). Effect of alpha-difluoromethylornithine in lung metastases before and after surgery of primary adenocarcinoma tumors in mice. Biol. Cell 53, 33–36. 10.1111/j.1768-322x.1985.tb00352.x.

88. Herr, H.W., Kleinert, E.L., Conti, P.S., Burchenal, J.H., and Whitmore, W.F. (1984). Effects of alpha-difluoromethylornithine and methylglyoxal bis(guanylhydrazone) on the growth of experimental renal adenocarcinoma in mice. Cancer Res. 44, 4382–4385.

89. Kubota, S., Ohsawa, N., and Takaku, F. (1987). Effects of DL-alpha-difluoromethylornithine on the growth and metastasis of B16 melanoma in vivo. Int. J. Cancer 39, 244–247. 10.1002/ijc.2910390220.

90. Kubota, S., Kiyosawa, H., Nomura, Y., Yamada, T., and Seyama, Y. (1997). Ornithine decarboxylase overexpression in mouse 10T1/2 fibroblasts: cellular transformation and invasion. J. Natl. Cancer Inst. 89, 567–571. 10.1093/jnci/89.8.567.

91. Hibino, S., Eto, S., Hangai, S., Endo, K., Ashitani, S., Sugaya, M., Osawa, T., Soga, T., Taniguchi, T., and Yanai, H. (2023). Tumor cell-derived spermidine is an oncometabolite that suppresses TCR clustering for intratumoral CD8+ T cell activation. Proc. Natl. Acad. Sci. U. S. A. 120, e2305245120. 10.1073/pnas.2305245120.

92. Sholler, G.L.S., Ferguson, W., Bergendahl, G., Bond, J.P., Neville, K., Eslin, D., Brown, V., Roberts, W., Wada, R.K., Oesterheld, J., et al. (2018). Maintenance DFMO Increases Survival in High Risk Neuroblastoma. Sci. Rep. 8, 14445. 10.1038/s41598-018-32659-w.

93. Simoneau, A.R., Gerner, E.W., Nagle, R., Ziogas, A., Fujikawa-Brooks, S., Yerushalmi, H., Ahlering, T.E., Lieberman, R., McLaren, C.E., Anton-Culver, H., et al. (2008). The Effect of Difluoromethylornithine on Decreasing Prostate Size and Polyamines in Men: Results of a Year-Long Phase IIb Randomized Placebo-Controlled Chemoprevention Trial. Cancer Epidemiol. Biomark. Prev. Publ. Am. Assoc. Cancer Res. Cosponsored Am. Soc. Prev. Oncol. 17, 292–299. 10.1158/1055-9965.EPI-07-0658.

94. Holbert, C.E., Cullen, M.T., Casero, R.A., and Stewart, T.M. (2022). Polyamines in cancer: integrating organismal metabolism and antitumour immunity. Nat. Rev. Cancer 22, 467–480. 10.1038/s41568-022-00473-2.

95. Offenbacher, R., Jackson, K.W., Hayashi, M., Zhang, J., Peng, D., Tan, Y., Stewart, T.M., Ciero, P., Foley, J., Casero, R.A., et al. (2024). Polyamine Depletion by D, L-alpha-difluoromethylornithine Inhibits Ewing Sarcoma Metastasis by Inducing Ferroptosis. BioRxiv Prepr. Serv. Biol., 2024.06.14.599064. 10.1101/2024.06.14.599064.

96. Li, M., Yu, X., Ouyang, S., Chen, X., Yu, H., Liu, Y., Li, Z., Yu, C., Kang, R., Gaillet, C., et al. (2026). Spermine is an endogenous iron chelator that inhibits ferroptosis. Nature. 10.1038/s41586-026-10597-2.

97. Lane, D.J.R., Bae, D.-H., Siafakas, A.R., Suryo Rahmanto, Y., Al-Akra, L., Jansson, P.J., Casero, R.A., and Richardson, D.R. (2018). Coupling of the polyamine and iron metabolism pathways in the regulation of proliferation: Mechanistic links to alterations in key polyamine biosynthetic and catabolic enzymes. Biochim. Biophys. Acta Mol. Basis Dis. 1864, 2793–2813. 10.1016/j.bbadis.2018.05.007.

98. Dong, Y., Kang, H., Peng, R., Liu, Z., Liao, F., Hu, S.-A., Ding, W., Wang, P., Yang, P., Zhu, M., et al. (2024). A clinical-stage Nrf2 activator suppresses osteoclast differentiation via the iron-ornithine axis. Cell Metab. 36, 1679–1695.e6. 10.1016/j.cmet.2024.03.005.

99. Lam, B., Kajderowicz, K.M., Keys, H.R., Roessler, J.M., Frenkel, E.M., Kirkland, A., Bisht, P., El-Brolosy, M.A., Jaenisch, R., Bell, G.W., et al. (2024). Multi-species genome-wide CRISPR screens identify conserved suppressors of cold-induced cell death. BioRxiv Prepr. Serv. Biol., 2024.07.25.605098. 10.1101/2024.07.25.605098.

100. Hares, G.B., Fernelius, W.C., and Douglas, B.E. (1956). Equilibrium Constants for the Formation of Complexes between Metal Ions and Polyamines1,2. J. Am. Chem. Soc. 78, 1816–1818. 10.1021/ja01590a011.

101. Bertsch, C.R., Fernelius, W.C., and Block, B.P. (1958). A Thermodynamic Study of Some Complexes of Metal Ions with Polyamines. J. Phys. Chem. 62, 444–450. 10.1021/j150562a018.

102. Palmer, B.N., and Powell, H.K.J. (1974). Complex formation between 4,9-diazadodecane-1,12-diamine (spermine) and copper(II) ions and protons in aqueous solution. J. Chem. Soc. Dalton Trans., 2086–2089. 10.1039/DT9740002086.

103. Barbucci, R., Campbell, M.J.M., Cannas, M., and Marongiu, G. (1980). Seven membered chelate rings in copper(II) complexes with spermidine. Inorganica Chim. Acta 46, 135–138. 10.1016/S0020-1693(00)84181-X.

104. LØVaas, E. (1996). Antioxidative and Metal-Chelating Effects of Polyamines11Dedicated to the memory of David E. Green: Long gone but not forgotten. In Advances in Pharmacology, H. Sies, ed. (Academic Press), pp. 119–149. 10.1016/S1054-3589(08)60982-5.

105. Kamlin, C.O.F., M. Jenkins, T., L Heise, J., and S. Amin, N. (2024). Trientine Tetrahydrochloride, From Bench to Bedside: A Narrative Review. Drugs 84, 1509–1518. 10.1007/s40265-024-02099-0.

106. Roberts, E.A., Schilsky, M.L., and American Association for Study of Liver Diseases (AASLD) (2008). Diagnosis and treatment of Wilson disease: an update. Hepatol. Baltim. Md 47, 2089–2111. 10.1002/hep.22261.

107. Perry, A.R., Pagliuca, A., Fitzsimons, E.J., Mufti, G.J., and Williams, R. (1996). Acquired sideroblastic anaemia induced by a copper-chelating agent. Int. J. Hematol. 64, 69–72. 10.1016/0925-5710(96)00457-4.

108. Tangella, A.V., Gajre, A.S., Chirumamilla, P.C., and Rathhan, P.V. Difluoromethylornithine (DFMO) and Neuroblastoma: A Review. Cureus 15, e37680. 10.7759/cureus.37680.

109. Ramirez, A., Heimbach, A., Gründemann, J., Stiller, B., Hampshire, D., Cid, L.P., Goebel, I., Mubaidin, A.F., Wriekat, A.-L., Roeper, J., et al. (2006). Hereditary parkinsonism with dementia is caused by mutations in ATP13A2, encoding a lysosomal type 5 P-type ATPase. Nat. Genet. 38, 1184–1191. 10.1038/ng1884.

110. Kırımtay, K., Temizci, B., Gültekin, M., Yapıcı, Z., and Karabay, A. (2021). Novel mutations in ATP13A2 associated with mixed neurological presentations and iron toxicity due to nonsense-mediated decay. Brain Res. 1750, 147167. 10.1016/j.brainres.2020.147167.

111. Schneider, S.A., Paisan-Ruiz, C., Quinn, N.P., Lees, A.J., Houlden, H., Hardy, J., and Bhatia, K.P. (2010). ATP13A2 mutations (PARK9) cause neurodegeneration with brain iron accumulation. Mov. Disord. 25, 979–984. 10.1002/mds.22947.

112. Rajagopalan, S., Rane, A., Chinta, S.J., and Andersen, J.K. (2016). Regulation of ATP13A2 via PHD2-HIF1α Signaling Is Critical for Cellular Iron Homeostasis: Implications for Parkinson’s Disease. J. Neurosci. Off. J. Soc. Neurosci. 36, 1086–1095. 10.1523/JNEUROSCI.3117-15.2016.

113. Gitler, A.D., Chesi, A., Geddie, M.L., Strathearn, K.E., Hamamichi, S., Hill, K.J., Caldwell, K.A., Caldwell, G.A., Cooper, A.A., Rochet, J.-C., et al. (2009). Alpha-synuclein is part of a diverse and highly conserved interaction network that includes PARK9 and manganese toxicity. Nat. Genet. 41, 308–315. 10.1038/ng.300.

114. Schmidt, K., Wolfe, D.M., Stiller, B., and Pearce, D.A. (2009). Cd2+, Mn2+, Ni2+ and Se2+ toxicity to *Saccharomyces cerevisiae* lacking YPK9p the orthologue of human ATP13A2. Biochem. Biophys. Res. Commun. 383, 198–202. 10.1016/j.bbrc.2009.03.151.

115. Kong, S.M.Y., Chan, B.K.K., Park, J.-S., Hill, K.J., Aitken, J.B., Cottle, L., Farghaian, H., Cole, A.R., Lay, P.A., Sue, C.M., et al. (2014). Parkinson’s disease-linked human PARK9/ATP13A2 maintains zinc homeostasis and promotes α-Synuclein externalization via exosomes. Hum. Mol. Genet. 23, 2816–2833. 10.1093/hmg/ddu099.

116. Rinaldi, D.E., Corradi, G.R., Cuesta, L.M., Adamo, H.P., and de Tezanos Pinto, F. (2015). The Parkinson-associated human P5B-ATPase ATP13A2 protects against the iron-induced cytotoxicity. Biochim. Biophys. Acta 1848, 1646–1655. 10.1016/j.bbamem.2015.04.008.

117. Martin, S., van Veen, S., Holemans, T., Demirsoy, S., van den Haute, C., Baekelandt, V., Agostinis, P., Eggermont, J., and Vangheluwe, P. (2016). Protection against Mitochondrial and Metal Toxicity Depends on Functional Lipid Binding Sites in ATP13A2. Park. Dis. 2016, e9531917. 10.1155/2016/9531917.

118. Liu, B., Azfar, M., Legchenko, E., West, J.A., Martin, S., Van den Haute, C., Baekelandt, V., Wharton, J., Howard, L., Wilkins, M.R., et al. (2024). ATP13A3 variants promote pulmonary arterial hypertension by disrupting polyamine transport. Cardiovasc. Res. 120, 756–768. 10.1093/cvr/cvae068.

119. Soon, E., Treacy, C.M., Toshner, M.R., MacKenzie-Ross, R., Manglam, V., Busbridge, M., Sinclair-McGarvie, M., Arnold, J., Sheares, K.K., Morrell, N.W., et al. (2011). Unexplained iron deficiency in idiopathic and heritable pulmonary arterial hypertension. Thorax 66, 326–332. 10.1136/thx.2010.147272.

120. Mazhar, M., Din, A.U., Ali, H., Yang, G., Ren, W., Wang, L., Fan, X., and Yang, S. (2021). Implication of ferroptosis in aging. Cell Death Discov. 7, 149. 10.1038/s41420-021-00553-6.

121. Zhuang, X., Wang, Q., Joost, S., Ferrena, A., Humphreys, D.T., Li, Z., Blum, M., Krause, K., Ding, S., Landais, Y., et al. (2025). Ageing limits stemness and tumorigenesis by reprogramming iron homeostasis. Nature 637, 184–194. 10.1038/s41586-024-08285-0.

122. Tamari, K., Konno, M., Asai, A., Koseki, J., Hayashi, K., Kawamoto, K., Murai, N., Matsufuji, S., Isohashi, F., Satoh, T., et al. (2018). Polyamine flux suppresses histone lysine demethylases and enhances ID1 expression in cancer stem cells. Cell Death Discov. 4, 104. 10.1038/s41420-018-0117-7.

123. Müller, S., Sindikubwabo, F., Cañeque, T., Lafon, A., Versini, A., Lombard, B., Loew, D., Wu, T.-D., Ginestier, C., Charafe-Jauffret, E., et al. (2020). CD44 regulates epigenetic plasticity by mediating iron endocytosis. Nat. Chem. 12, 929–938. 10.1038/s41557-020-0513-5.

124. Mai, T.T., Hamaï, A., Hienzsch, A., Cañeque, T., Müller, S., Wicinski, J., Cabaud, O., Leroy, C., David, A., Acevedo, V., et al. (2017). Salinomycin kills cancer stem cells by sequestering iron in lysosomes. Nat. Chem. 9, 1025–1033. 10.1038/nchem.2778.

125. Schonberg, D.L., Miller, T.E., Wu, Q., Flavahan, W.A., Das, N.K., Hale, J.S., Hubert, C.G., Mack, S.C., Jarrar, A.M., Karl, R.T., et al. (2015). Preferential Iron Trafficking Characterizes Glioblastoma Stem-like Cells. Cancer Cell 28, 441–455. 10.1016/j.ccell.2015.09.002.

126. Kakhlon, O., and Cabantchik, Z.I. (2002). The labile iron pool: characterization, measurement, and participation in cellular processes(1). Free Radic. Biol. Med. 33, 1037–1046. 10.1016/s0891-5849(02)01006-7.

127. Cabantchik, Z.I. (2014). Labile iron in cells and body fluids: physiology, pathology, and pharmacology. Front. Pharmacol. 5, 45. 10.3389/fphar.2014.00045.

128. Philpott, C.C., Patel, S.J., and Protchenko, O. (2020). Management versus miscues in the cytosolic labile iron pool: The varied functions of iron chaperones. Biochim. Biophys. Acta Mol. Cell Res. 1867, 118830. 10.1016/j.bbamcr.2020.118830.

129. Roper, J., Tammela, T., Cetinbas, N.M., Akkad, A., Roghanian, A., Rickelt, S., Almeqdadi, M., Wu, K., Oberli, M.A., Sánchez-Rivera, F.J., et al. (2017). In vivo genome editing and organoid transplantation models of colorectal cancer and metastasis. Nat. Biotechnol. 35, 569–576. 10.1038/nbt.3836.

130. Stewart, S.A., Dykxhoorn, D.M., Palliser, D., Mizuno, H., Yu, E.Y., An, D.S., Sabatini, D.M., Chen, I.S.Y., Hahn, W.C., Sharp, P.A., et al. (2003). Lentivirus-delivered stable gene silencing by RNAi in primary cells. RNA N. Y. N 9, 493–501. 10.1261/rna.2192803.

131. Adelmann, C.H., Traunbauer, A.K., Chen, B., Condon, K.J., Chan, S.H., Kunchok, T., Lewis, C.A., and Sabatini, D.M. (2020). MFSD12 mediates the import of cysteine into melanosomes and lysosomes. Nature 588, 699–704. 10.1038/s41586-020-2937-x.

132. Dobin, A., Davis, C.A., Schlesinger, F., Drenkow, J., Zaleski, C., Jha, S., Batut, P., Chaisson, M., and Gingeras, T.R. (2013). STAR: ultrafast universal RNA-seq aligner. Bioinforma. Oxf. Engl. 29, 15–21. 10.1093/bioinformatics/bts635.

133. Alpern, D., Gardeux, V., Russeil, J., Mangeat, B., Meireles-Filho, A.C.A., Breysse, R., Hacker, D., and Deplancke, B. (2019). BRB-seq: ultra-affordable high-throughput transcriptomics enabled by bulk RNA barcoding and sequencing. Genome Biol. 20, 71. 10.1186/s13059-019-1671-x.

134. Chen, S. (2025). fastp 1.0: An ultra-fast all-round tool for FASTQ data quality control and preprocessing. iMeta 4, e70078. 10.1002/imt2.70078.

135. Liu, D. (2019). Algorithms for efficiently collapsing reads with Unique Molecular Identifiers. PeerJ 7, e8275. 10.7717/peerj.8275.

136. Liao, Y., Smyth, G.K., and Shi, W. (2014). featureCounts: an efficient general purpose program for assigning sequence reads to genomic features. Bioinformatics 30, 923–930. 10.1093/bioinformatics/btt656.

137. Robinson, M.D., McCarthy, D.J., and Smyth, G.K. (2010). edgeR: a Bioconductor package for differential expression analysis of digital gene expression data. Bioinformatics 26, 139–140. 10.1093/bioinformatics/btp616.

