## Supplementary Figures for "Polyamines buffer labile iron to suppress ferroptosis"

**Figure S1. Polyamine depletion sensitizes cells to ferroptosis independently of hypusination, cell cycle arrest, antioxidant depletion, and radical-trapping activity, related to Figure 3**

(A) Enzymatic quantification of polyamines in K562 cells treated with DFMO (0.5 mM, 72 h) (n=3).

(B) Relative live cell count of AAVS1 control and GPX4 knockout cells plated at indicated cell densities with or without 1 μM Liproxstatin-1 (Lip-1) after 72 hours. Live cells were measured by CellTiter-Glo and normalized to AAVS1 knockout control at each seeding density. Data shown as mean +/- SEM. Data is representative of three independent experiments (n=3).

(C) Cell death of AAVS1 control and GPX4 knockout cells plated at the indicated cell densities with or without 1 μM Lip-1 after 72 hours. Cell death was measured by Sytox Green fluorescence intensity. Data shown as mean +/- SEM. Data is representative of three independent experiments (n=3).

(D) Chemical structure of sardomozide.

(E) Polyamine levels in K562 cells treated with sardomozide (5 μM, 72 h) and/or spermidine (10 μM, 72 h) using the previously reported genetically encoded sensor (n>10,000 cells).

(F) Viability of MEL cells treated with ML162 (24 h) following pre-treatment with sardomozide (5 μM, 72 h). DMSO served as vehicle control (n=3).

(G) Viability of K562 cells treated with ML162 (2 μM, 24 h) after pre-treatment with GC7 (40 μM, 96 h). Acetic acid served as vehicle control. Liproxstatin (2.5 μM) was co-administered with ML162 (n=3).

(H) Cumulative population doublings (CPDs) of K562 cells treated with palbociclib (10 μM). Cells were passaged every 2 days (n=3).

(I) Viability of AAVS1 or GPX4 KO K562 cells treated with palbociclib (Pal) (10 μM, 6 days) (n=3).

(J) Fluorescence of STY-BODIPY oxidation (2 μM STY-BODIPY, 0.1 mM liposomal soy PC, 200 μM DTUN) in the presence of putrescine or PMC (positive control) (n=1, representative of 2 independent experiments).

(K) Untargeted metabolomics of K562 cells treated with sardomozide (5 μM, 72 h). Highlighted metabolites are canonical antioxidant/redox metabolites (n=3).

(L-O) RNA-seq on K562 cells treated with sardomozide (5 μM, 72 h) with highlighted pathways (n=3).

Figure S1:

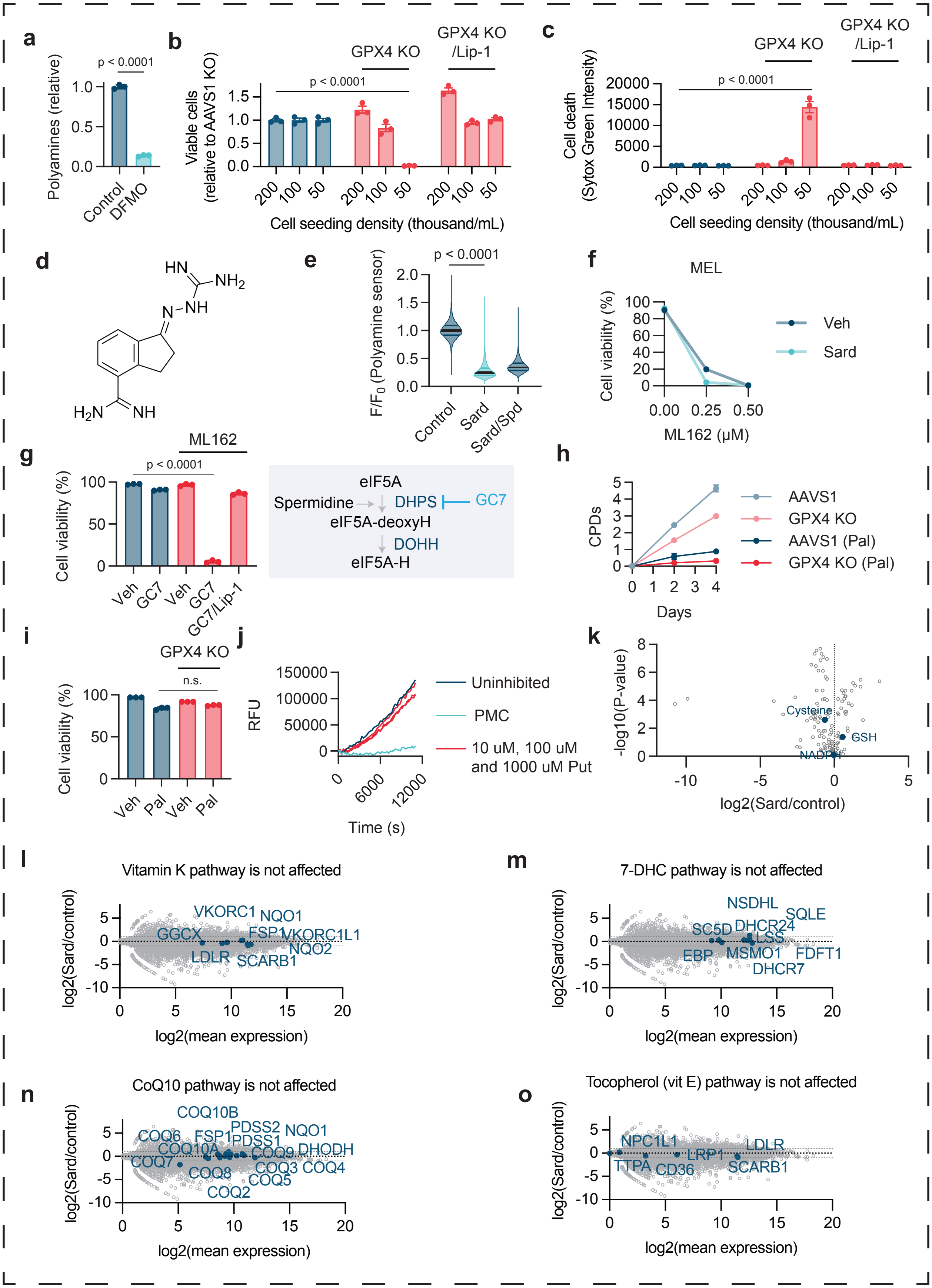

a
Polyamines (relative)
p < 0.0001
Control
DFMO
b
Viable cells (relative to AAVS1 KO)
GPX4 KO
GPX4 KO /Lip-1
p < 0.0001
Cell seeding density (thousand/mL)
c
Cell death (Sytox Green Intensity)
GPX4 KO
GPX4 KO /Lip-1
p < 0.0001
Cell seeding density (thousand/mL)
d
e
F/F0 (Polyamine sensor)
p < 0.0001
Control
Sard
Sard/Spd
f
MEL
Cell viability (%)
Veh
Sard
ML162 (μM)
g
ML162
Cell viability (%)
p < 0.0001
Veh
GC7
GC7/Lip-1
eIF5A
Spermidine
DHPS
GC7
eIF5A-deoxyH
DOHH
eIF5A-H
h
CPDs
AAVS1
GPX4 KO
AAVS1 (Pal)
GPX4 KO (Pal)
Days
i
GPX4 KO
n.s.
Cell viability (%)
Veh
Pal
j
RFU
Uninhibited
PMC
10 uM, 100 uM and 1000 uM Put
Time (s)
k
-log10(P-value)
Cysteine
GSH
NADPH
log2(Sard/control)
l
Vitamin K pathway is not affected
log2(Sard/control)
VKORC1
NQO1
GGCX
FSP1
VKORC1L1
NQO2
LDLR
SCARB1
log2(mean expression)
m
7-DHC pathway is not affected
log2(Sard/control)
NSDHL
SQLE
SC5D
DHCR24
LSS
EBP
MSMO1
FDFT1
DHCR7
log2(mean expression)
n
CoQ10 pathway is not affected
log2(Sard/control)
COQ10B
PDSS2
NQO1
COQ6
FSP1
PDSS1
COQ10A
COQ9
DHODH
COQ7
COQ8
COQ3
COQ4
COQ5
COQ2
log2(mean expression)
o
Tocopherol (vit E) pathway is not affected
log2(Sard/control)
NPC1L1
LDLR
LRP1
TTPA
CD36
SCARB1
log2(mean expression)

**Figure S2. Polyamine depletion induces metabolic and transcriptional changes without expanding PUFA-containing phospholipid substrates, related to Figure 3**

(A) Untargeted metabolomics of K562 cells treated with sardomozide (5 μM, 72 h). Pentose phosphate pathway metabolites are highlighted (n=3).

(B-D) Lipidomic profiling during polyamine depletion and iron chelation. Lipid class composition in vehicle, sardomozide (SARD) and DFO-treated K562 cells. Relative abundances are normalized to total lipid content and grouped by major lipid classes: triacylglycerol (TAG), sphingomyelin (SM), phosphatidylinositol (PI), phosphatidylethanolamine (PE), ether/plasmalogen phosphatidylethanolamine (PE-(O/P)), phosphatidylcholine (PC), ether/plasmalogen phosphatidylcholine (PC-(O/P)), lysophosphatidylcholine (LPC), ether/plasmalogen lysophosphatidylcholine (LPC-(O/P)), lysophosphatidylethanolamine (LPE), ether/plasmalogen lysophosphatidylethanolamine (LPE-(O/P)), hexosylceramide (HexCer), diacylglycerol (DG), ceramide (Cer), and cholesteryl ester (CE). Data is shown as the mean +/-SEM. n=3 technical replicates. Data is representative of 2 independent experiments.

(E-H) Heatmaps showing relative abundances of polyunsaturated lipid species across vehicle, DFO, and SARD-treated cells. Lipid species are PUFA-containing phosphatidylcholine lipids (PUFA PC), PUFA-containing ether/plasmalogen phosphatidylcholines (PUFA PC-O/P), PUFA-containing ether/plasmalogen phosphatidylethanolamines (PUFA PE-O/P), and PUFA-containing phosphatidylethanolamines (PUFA PE). Heatmaps display Z-score–normalized lipid abundances for each lipid species across conditions. Each row represents an individual lipid species and each column represents a replicate. PUFA-containing species were defined as lipids with more than two total acyl-chain double bonds. Z-score scale bars are common between different panels (n=3).

(I) RNA-seq on K562 cells treated with sardomozide (5 μM, 72 h) with highlighted genes (n=3).

Figure S2:

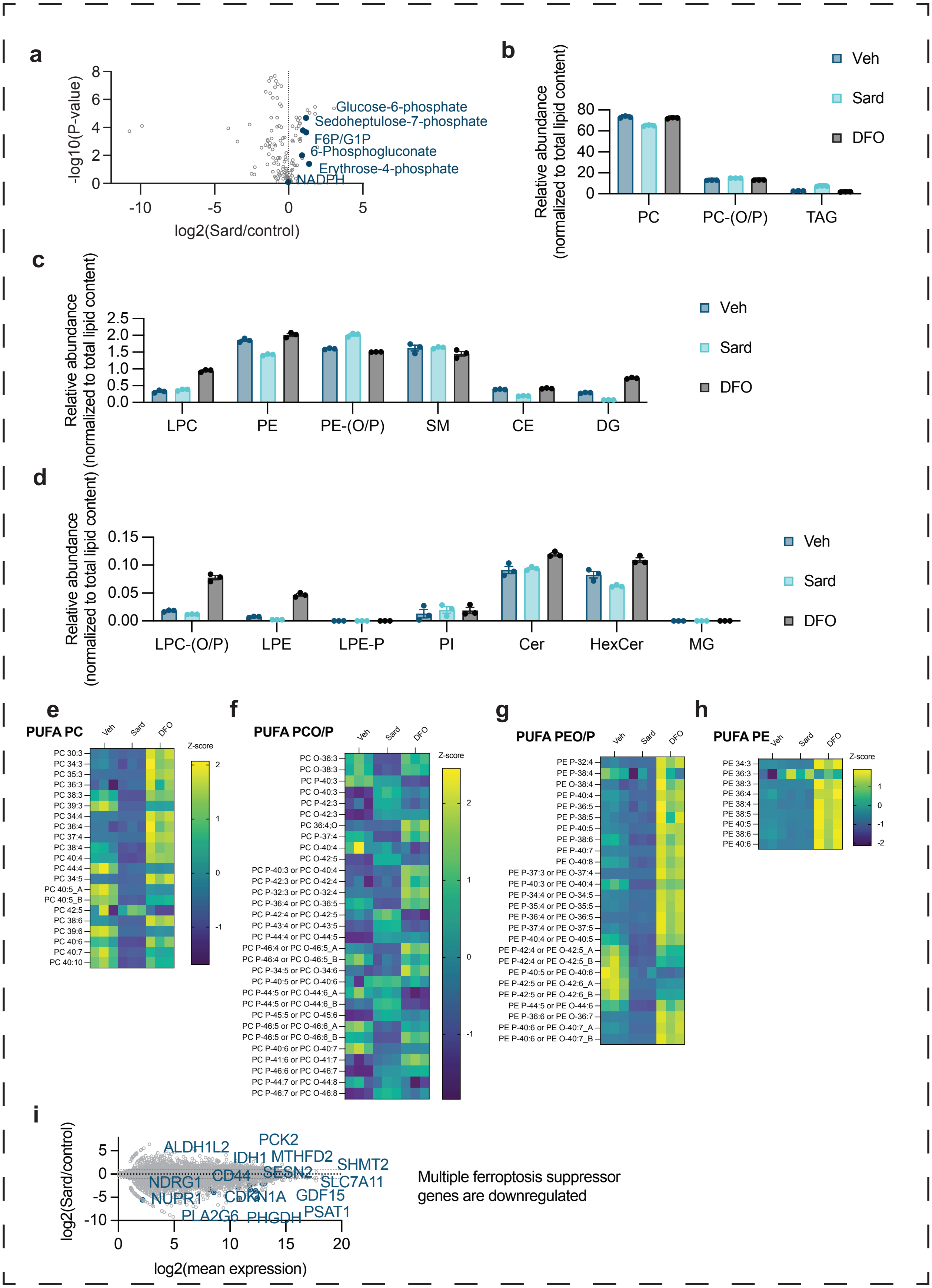
a
-log10(P-value)
log2(Sard/control)
Glucose-6-phosphate
Sedoheptulose-7-phosphate
F6P/G1P
6-Phosphogluconate
Erythrose-4-phosphate
NADPH
b
Relative abundance (normalized to total lipid content)
PC
PC-(O/P)
TAG
Veh
Sard
DFO
c
Relative abundance (normalized to total lipid content)
LPC
PE
PE-(O/P)
SM
CE
DG
Veh
Sard
DFO
d
Relative abundance (normalized to total lipid content)
LPC-(O/P)
LPE
LPE-P
PI
Cer
HexCer
MG
Veh
Sard
DFO
e
PUFA PC
Veh
Sard
DFO
Z-score
PC 30:3
PC 34:3
PC 35:3
PC 36:3
PC 38:3
PC 39:3
PC 34:4
PC 36:4
PC 37:4
PC 38:4
PC 40:4
PC 44:4
PC 34:5
PC 40:5_A
PC 40:5_B
PC 42:5
PC 38:6
PC 39:6
PC 40:6
PC 40:7
PC 40:10
f
PUFA PCO/P
Veh
Sard
DFO
Z-score
PC O-36:3
PC O-38:3
PC P-40:3
PC O-40:3
PC P-42:3
PC O-42:3
PC 36:4;O
PC P-37:4
PC O-40:4
PC O-42:5
PC P-40:3 or PC O-40:4
PC P-42:3 or PC O-42:4
PC P-32:3 or PC O-32:4
PC P-36:4 or PC O-36:5
PC P-42:4 or PC O-42:5
PC P-43:4 or PC O-43:5
PC P-44:4 or PC O-44:5
PC P-46:4 or PC O-46:5_A
PC P-46:4 or PC O-46:5_B
PC P-34:5 or PC O-34:6
PC P-40:5 or PC O-40:6
PC P-44:5 or PC O-44:6_A
PC P-44:5 or PC O-44:6_B
PC P-45:5 or PC O-45:6
PC P-46:5 or PC O-46:6_A
PC P-46:5 or PC O-46:6_B
PC P-40:6 or PC O-40:7
PC P-41:6 or PC O-41:7
PC P-46:6 or PC O-46:7
PC P-44:7 or PC O-44:8
PC P-46:7 or PC O-46:8
g
PUFA PEO/P
Veh
Sard
DFO
PE P-32:4
PE P-38:4
PE O-38:4
PE P-40:4
PE P-36:5
PE P-38:5
PE P-40:5
PE P-38:6
PE P-40:7
PE O-40:8
PE P-37:3 or PE O-37:4
PE P-40:3 or PE O-40:4
PE P-34:4 or PE O-34:5
PE P-35:4 or PE O-35:5
PE P-36:4 or PE O-36:5
PE P-37:4 or PE O-37:5
PE P-40:4 or PE O-40:5
PE P-42:4 or PE O-42:5_A
PE P-42:4 or PE O-42:5_B
PE P-40:5 or PE O-40:6
PE P-42:5 or PE O-42:6_A
PE P-42:5 or PE O-42:6_B
PE P-44:5 or PE O-44:6
PE P-36:6 or PE O-36:7
PE P-40:6 or PE O-40:7_A
PE P-40:6 or PE O-40:7_B
h
PUFA PE
Veh
Sard
DFO
Z-score
PE 34:3
PE 36:3
PE 38:3
PE 36:4
PE 38:4
PE 38:5
PE 40:5
PE 38:6
PE 40:6
i
log2(Sard/control)
log2(mean expression)
ALDH1L2
PCK2
IDH1
MTHFD2
SHMT2
CD44
SESN2
NDRG1
SLC7A11
NUPR1
CDKN1A
GDF15
PLA2G6
PHGDH
PSAT1
Multiple ferroptosis suppressor genes are downregulated

**Figure S3. Iron chelation reverses iron homeostasis changes without broadly altering transcriptional or lipidomic responses to polyamine depletion, related to Figure 4**

(A) Immunoblot analysis of FTH protein levels in K562 cells treated with ferric ammonium citrate (FAC; 50 µg/mL, 12 h) or ferrous ammonium sulfate (FAS; 10 µM, 12 h) (n=1).
(B) Quantitative proteomics of U-2OS cells treated with DFMO (1 mM, 2 days); polyamine metabolism related proteins (controls for the experiment) are highlighted (n=3).
(C) Quantitative proteomics of U-2OS cells treated with DFMO (1 mM, 2 days); iron metabolism-related proteins are highlighted (n=3).

(D-E) RNA-seq on K562 cells treated with sardomozide (5 µM, 72 h) and DFO rescue with highlighted genes (n=3).
(F-G) Lipidomic profiling during polyamine depletion and iron chelation. Lipid class composition in vehicle, sardomozide (SARD), DFO, and SARD + DFO-treated K562 cells. Relative abundances are normalized to total lipid content and grouped by major lipid classes: triacylglycerol (TAG), sphingomyelin (SM), phosphatidylethanolamine (PE), diacylglycerol (DG), cholesteryl ester (CE), ether/plasmalogen phosphatidylethanolamine (PE-(O/P)), phosphatidylcholine (PC), ether/plasmalogen phosphatidylcholine (PC-(O/P)), lysophosphatidylcholine (LPC). Data is shown as the mean +/- SEM. n=3 technical replicates. Data is representative of 2 independent experiments.

Figure S3:

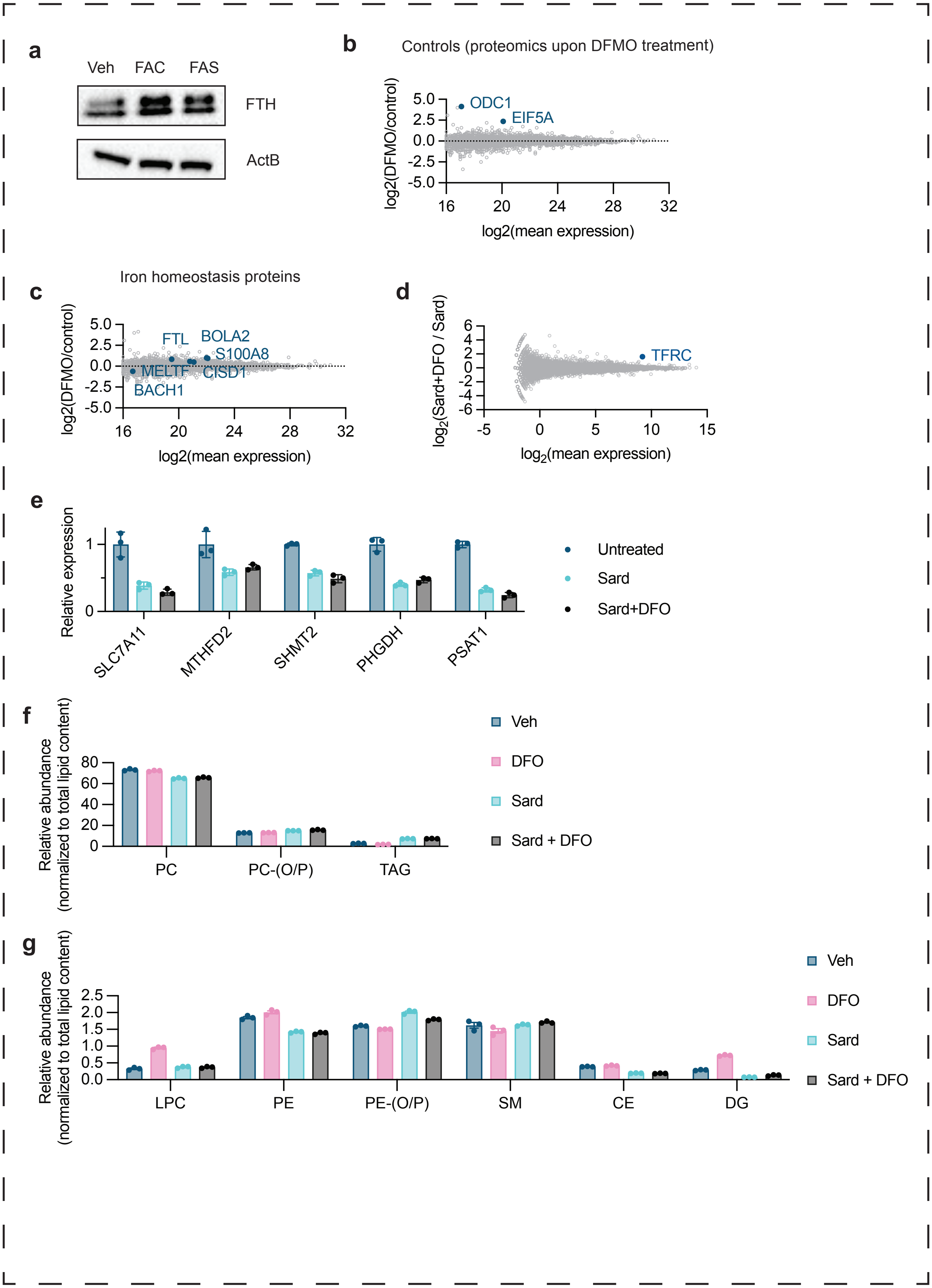
a
Veh
FAC
FAS
FTH
ActB
b
Controls (proteomics upon DFMO treatment)
log2(DFMO/control)
ODC1
EIF5A
log2(mean expression)
c
Iron homeostasis proteins
log2(DFMO/control)
FTL
BOLA2
S100A8
MELTF
CISD1
BACH1
log2(mean expression)
d
log2(Sard+DFO / Sard)
TFRC
log2(mean expression)
e
Relative expression
SLC7A11
MTHFD2
SHMT2
PHGDH
PSAT1
Untreated
Sard
Sard+DFO
f
Relative abundance (normalized to total lipid content)
PC
PC-(O/P)
TAG
Veh
DFO
Sard
Sard + DFO
g
Relative abundance (normalized to total lipid content)
LPC
PE
PE-(O/P)
SM
CE
DG
Veh
DFO
Sard
Sard + DFO

**Figure S4. Polyamine depletion and iron chelation minimally alter global lipid composition and PUFA-containing phospholipid abundance, related to Figure 4**

(A) Lipidomic profiling during polyamine depletion and iron chelation. Lipid class composition in vehicle, sardomozide (SARD), DFO, and SARD + DFO-treated K562 cells. Relative abundances are normalized to total lipid content and grouped by major lipid classes. Data is shown as the mean +/- SEM. n=3 technical replicates. Data is representative of 2 independent experiments.

(B-E) Heatmaps showing relative abundances of polyunsaturated lipid species across vehicle, DFO, SARD, and SARD + DFO-treated cells. Heatmaps display Z-score–normalized lipid abundances for each lipid species across conditions. Each row represents an individual lipid species and each column represents a replicate. PUFA-containing species were defined as lipids with more than two total acyl-chain double bonds. Each heatmap displays Z-score–normalized abundances with the indicated scale bar. n=3 technical replicates. Data is representative of 2 independent experiments.

Figure S4:

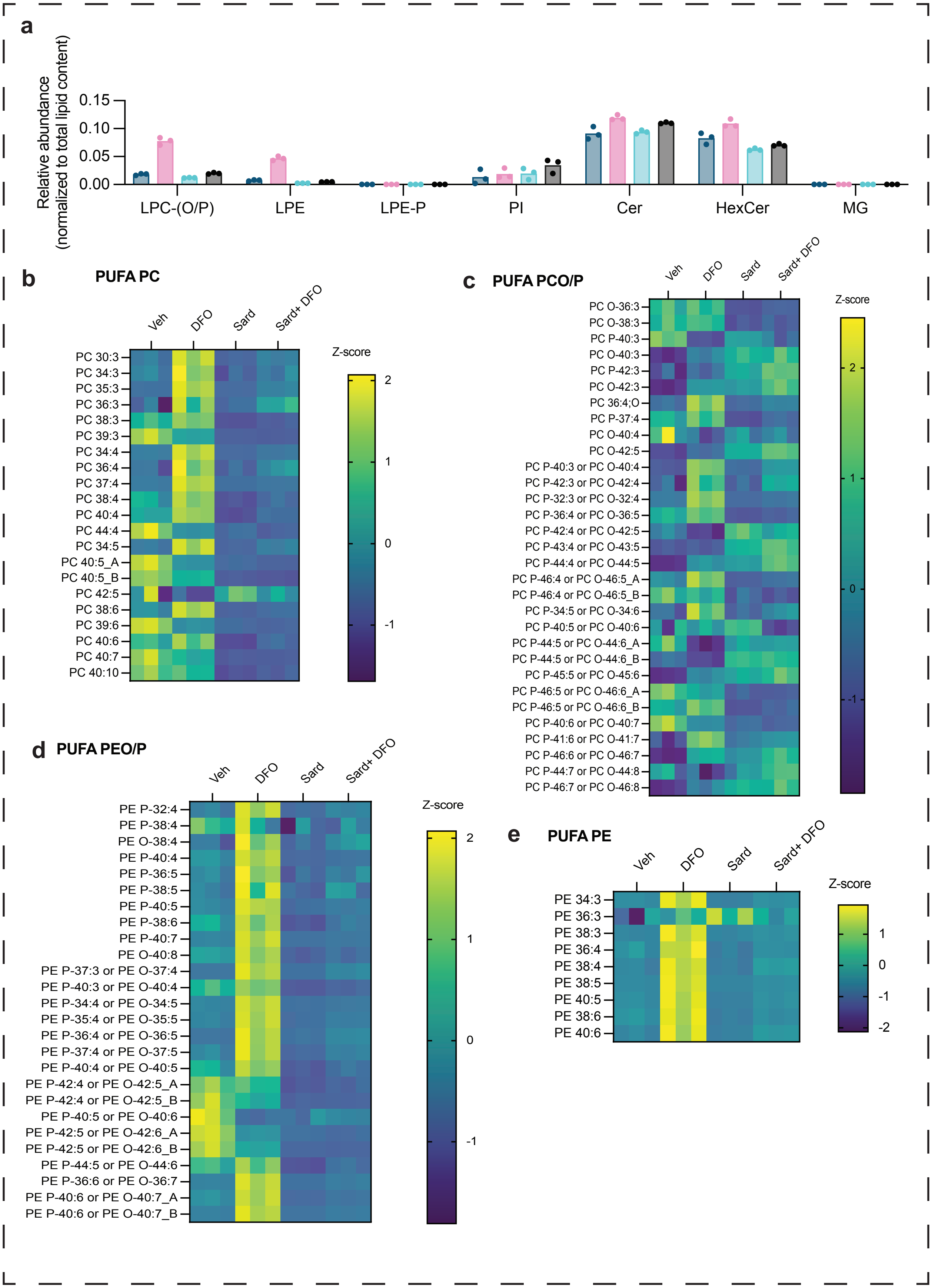

a
Relative abundance
(normalized to total lipid content)
0.15
0.10
0.05
0.00
LPC-(O/P)
LPE
LPE-P
PI
Cer
HexCer
MG
b
PUFA PC
Veh
DFO
Sard
Sard+ DFO
PC 30:3
PC 34:3
PC 35:3
PC 36:3
PC 38:3
PC 39:3
PC 34:4
PC 36:4
PC 37:4
PC 38:4
PC 40:4
PC 44:4
PC 34:5
PC 40:5_A
PC 40:5_B
PC 42:5
PC 38:6
PC 39:6
PC 40:6
PC 40:7
PC 40:10
Z-score
2
1
0
-1
c
PUFA PCO/P
Veh
DFO
Sard
Sard+ DFO
PC O-36:3
PC O-38:3
PC P-40:3
PC O-40:3
PC P-42:3
PC O-42:3
PC 36:4;O
PC P-37:4
PC O-40:4
PC O-42:5
PC P-40:3 or PC O-40:4
PC P-42:3 or PC O-42:4
PC P-32:3 or PC O-32:4
PC P-36:4 or PC O-36:5
PC P-42:4 or PC O-42:5
PC P-43:4 or PC O-43:5
PC P-44:4 or PC O-44:5
PC P-46:4 or PC O-46:5_A
PC P-46:4 or PC O-46:5_B
PC P-34:5 or PC O-34:6
PC P-40:5 or PC O-40:6
PC P-44:5 or PC O-44:6_A
PC P-44:5 or PC O-44:6_B
PC P-45:5 or PC O-45:6
PC P-46:5 or PC O-46:6_A
PC P-46:5 or PC O-46:6_B
PC P-40:6 or PC O-40:7
PC P-41:6 or PC O-41:7
PC P-46:6 or PC O-46:7
PC P-44:7 or PC O-44:8
PC P-46:7 or PC O-46:8
Z-score
2
1
0
-1
d
PUFA PEO/P
Veh
DFO
Sard
Sard+ DFO
PE P-32:4
PE P-38:4
PE O-38:4
PE P-40:4
PE P-36:5
PE P-38:5
PE P-40:5
PE P-38:6
PE P-40:7
PE O-40:8
PE P-37:3 or PE O-37:4
PE P-40:3 or PE O-40:4
PE P-34:4 or PE O-34:5
PE P-35:4 or PE O-35:5
PE P-36:4 or PE O-36:5
PE P-37:4 or PE O-37:5
PE P-40:4 or PE O-40:5
PE P-42:4 or PE O-42:5_A
PE P-42:4 or PE O-42:5_B
PE P-40:5 or PE O-40:6
PE P-42:5 or PE O-42:6_A
PE P-42:5 or PE O-42:6_B
PE P-44:5 or PE O-44:6
PE P-36:6 or PE O-36:7
PE P-40:6 or PE O-40:7_A
PE P-40:6 or PE O-40:7_B
Z-score
2
1
0
-1
e
PUFA PE
Veh
DFO
Sard
Sard+ DFO
PE 34:3
PE 36:3
PE 38:3
PE 36:4
PE 38:4
PE 38:5
PE 40:5
PE 38:6
PE 40:6
Z-score
1
0
-1
-2

**Figure S5. Validation of iron homeostasis measurements and optimization of the genetically encoded iron reporter, related to Figure 4 and 5**

(A-E) Gene essentiality scores for TFRC and AMD1 from the Cancer Cell Line Encyclopedia (DepMap) across different cancer types: (A) prostate, (B) bladder, (C) eye, (D) head and neck, and (E) skin cancer cell lines.

(F) Quantification of the polyamine reporter (eYFP/mCherry ratio) in subcutaneous K562 tumors from mice treated with vehicle (control) or combined DFMO/AMXT-1501 for 14 days prior to reporter induction. Bars show mean ± SD. Combined treatment significantly reduced reporter signal relative to control ($p < 0.05$, unpaired two-tailed t test) (n=5).

(G) MA plot of RNA-seq data from DFMO/AMXT-1501-treated versus control K562 tumors, plotting log2 fold change (DFMO-AMXT/control) against log2 mean expression across all detected transcripts (n=5).

(H-I) Representative fluorescence micrographs (F) and flow cytometry quantification (G) of RhoNox-1 in U-2OS cells under indicated treatments. DFO, 100 µM (48 h) and Sard, 5 µM h). Mean ± standard deviation shown. n: ≥2 independent experiments. Scale bars, 10 µm.

(J) Redox-active $Fe^{2+}$ content measured with RhoNox-M and LysoTracker in cells treated with sardomozide (5 µM, 72 h). FAS (50 µM, 5 h) as positive control (n=3).

(K-M) Flow cytometry quantification of translation initiation efficiency in K562 cells expressing iron sensor with varying number of iron-responsive elements (IREs) upstream of EBFP2 under indicated treatments. Translation efficiency is reported as the ratio of EBFP2 to EGFP fluorescence (F = EBFP2/EGFP), normalized to control cells (F0). Cells were analyzed under the indicated treatments. DFO (50 µM, 48 h) and FAC (10 mM, 24 h). Each panel shows the median ± interquartile range from ≥100 cells, with 50 individual data points plotted. Data represent ≥2 independent experiments. Statistical significance was assessed using a two-tailed Student's t-test.

(N) Immunoblot analysis of iron metabolism proteins in K562 cells upon expression of iron sensor using doxycycline (n=1).

Figure S5:

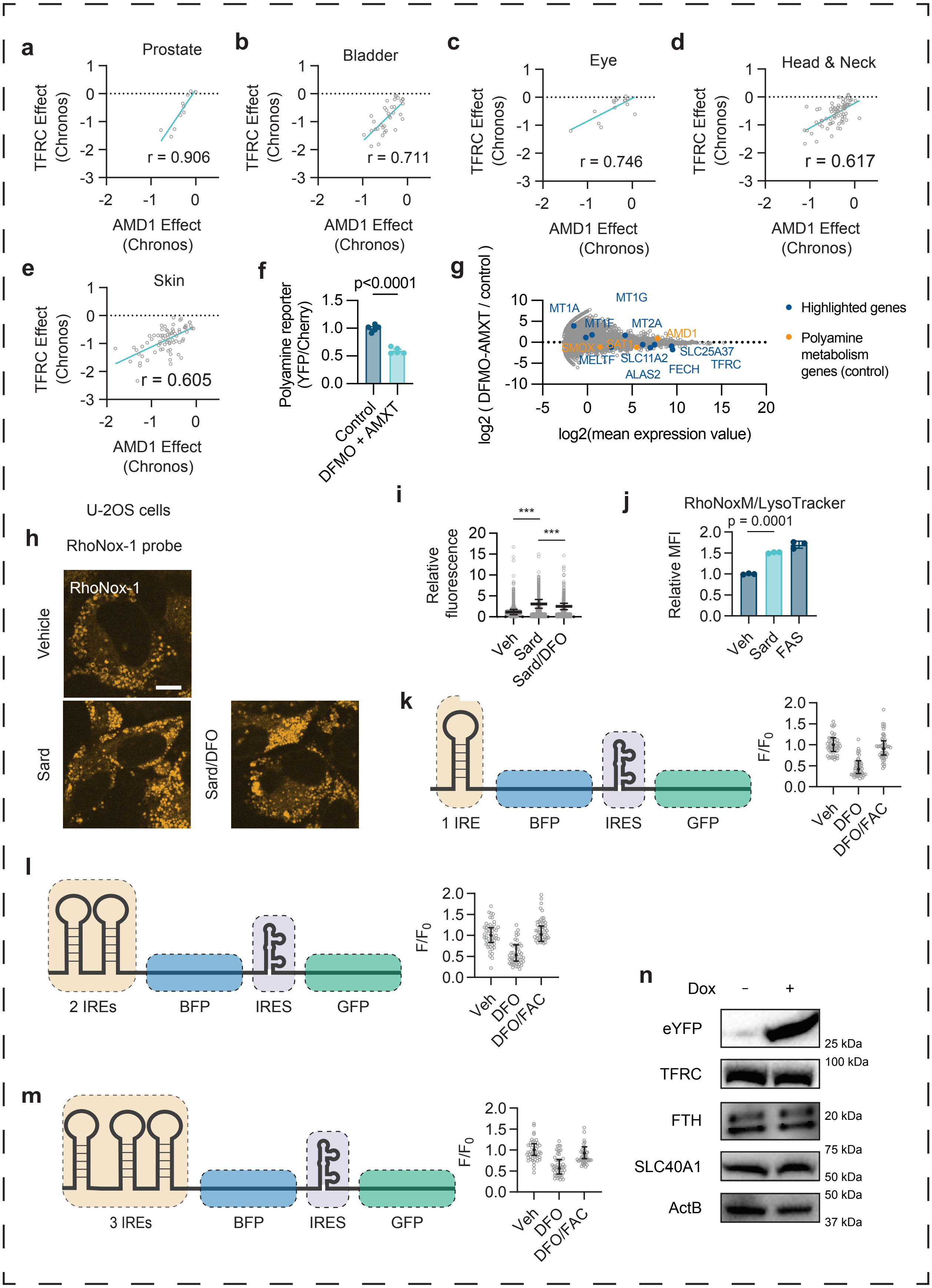

a
Prostate
TFRC Effect (Chronos)
AMD1 Effect (Chronos)
r = 0.906
b
Bladder
r = 0.711
c
Eye
r = 0.746
d
Head & Neck
r = 0.617
e
Skin
r = 0.605
f
p<0.0001
Polyamine reporter (YFP/Cherry)
Control
DFMO + AMXT
g
log2 ( DFMO-AMXT / control )
log2(mean expression value)
MT1A
MT1G
MT1F
MT2A
AMD1
SMOX
SAT1
MELTF
SLC11A2
SLC25A37
ALAS2
FECH
TFRC
Highlighted genes
Polyamine metabolism genes (control)
U-2OS cells
h
RhoNox-1 probe
RhoNox-1
Vehicle
Sard
Sard/DFO
i
Relative fluorescence
***
Veh
Sard
Sard/DFO
j
RhoNoxM/LysoTracker
p = 0.0001
Relative MFI
Veh
Sard
FAS
k
1 IRE
BFP
IRES
GFP
F/F0
Veh
DFO
DFO/FAC
l
2 IREs
BFP
IRES
GFP
m
3 IREs
BFP
IRES
GFP
n
Dox
-
+
eYFP
TFRC
FTH
SLC40A1
ActB
25 kDa
100 kDa
20 kDa
75 kDa
50 kDa
50 kDa
37 kDa

**Figure S6. Additional biochemical evidence for $Fe^{2+}$ coordination by polyamines from fluorescence competition and Mössbauer spectroscopy, related to Figure 6**

(A) Effect of addition of increasing concentrations of spermidine and spermine on formation of FerroOrange-$Fe^{2+}$ complex. Experimental conditions: 200 mM CAPS, pH 10.5, 50 μM $FeCl_2$, 1 mM DFO, 10 μM FerroOrange, 5-100 mM spermidine/spermine. DFO used as a positive control (n=3 except 5 for control).

Effects of ATP and spermine on the Mössbauer spectra of $^{57}Fe^{2+}$. (Left) Mössbauer spectra of 1 mM $^{57}Fe^{2+}$ alone (top), in the presence of 1 mM ATP (middle), and in the presence of 1 mM ATP plus 15 mM spermine (bottom). (Right) Difference spectra highlighting ligand-induced spectral changes. Positive features correspond to $Fe^{2+}$ species formed upon addition of ATP (top) or ATP plus spermine (middle), whereas negative features arise from depletion of free $Fe^{2+}$. The spectral contribution associated with spermine binding in the presence of ATP was isolated by second-order differencing (bottom). This component is well described by a quadrupole doublet (teal fit) supporting a spermine-dependent perturbation of the $Fe^{2+}$ coordination environment even in the presence of ATP. Quantitative analysis indicates that spermine accounts for ~5% of the total $Fe^{2+}$ population under these conditions. Spectra were recorded at 80 K in zero applied magnetic field. Experimental conditions: 1 mM $^{57}Fe^{2+}$, 50 mM HEPES (pH 7.5). Experiments are representative of n = 2 independent sample preparations.

Effects of putrescine and spermine on the Mössbauer spectra of $^{57}Fe^{2+}$ at pH 7.5. (Top) Mössbauer spectrum of 1 mM $^{57}Fe^{2+}$ in the presence of 40 mM putrescine (black trace). The putative $Fe^{2+}$-putrescine component (pink trace) was obtained by subtraction of 95% of the $Fe^{2+}$ control spectrum (green trace), indicating that only a minor fraction (~5%) of the $Fe^{2+}$ population is perturbed by putrescine under these conditions. (Bottom) Mössbauer spectrum of 1 mM $^{57}Fe^{2+}$ in the presence of 40 mM spermine (black trace). The $Fe^{2+}$-spermine component (gold trace) was obtained by subtraction of 90% of the $Fe^{2+}$ control spectrum (green trace), corresponding to approximately 10% of the total $Fe^{2+}$ population. Spectra were recorded at 80 K in zero applied magnetic field. Experimental conditions: 1 mM $^{57}Fe^{2+}$, 40 mM polyamine, and 200 mM HEPES (pH 7.5). Experiments are representative of n = 2 independent sample preparations.

Effects of putrescine and spermine on the Mössbauer spectra of $^{57}Fe^{2+}$ at pH 10. (Top) Mössbauer spectrum of $^{57}Fe^{2+}$ in the presence of 40 mM putrescine (black trace) overlaid with the $^{57}Fe^{2+}$ control spectrum (green trace). Putrescine does not exhibit detectable binding under these conditions, as evidenced by the near-complete overlap of the two spectra. The residual difference spectrum (pink trace) is consistent with minor variations in Fe concentration between samples rather than formation of a distinct $Fe^{2+}$-putrescine complex. (Middle) Mössbauer spectrum of $^{57}Fe^{2+}$ in the presence of 40 mM spermine (black trace). The gold trace corresponds to the $Fe^{2+}$-spermine component obtained after subtraction of 75% of the $Fe^{2+}$ control spectrum (green trace), indicating that approximately 25% of the $Fe^{2+}$ is present as an $Fe^{2+}$-spermine complex. All spectra were recorded at 80 K and in zero applied magnetic field. (Bottom) Difference spectrum corresponding to the accumulated $Fe^{2+}$-spermine complex (black vertical lines) together with simulations of two spectral subcomponents, consistent with the presence of two distinct $Fe^{2+}$-spermine species, likely arising from different spermine protonation states at pH 10. Experimental conditions: 1 mM $^{57}Fe^{2+}$, 40 mM polyamine, 50 mM CHES (pH 10). Experiments are representative of n = 2 independent sample preparations.

Figure S6:

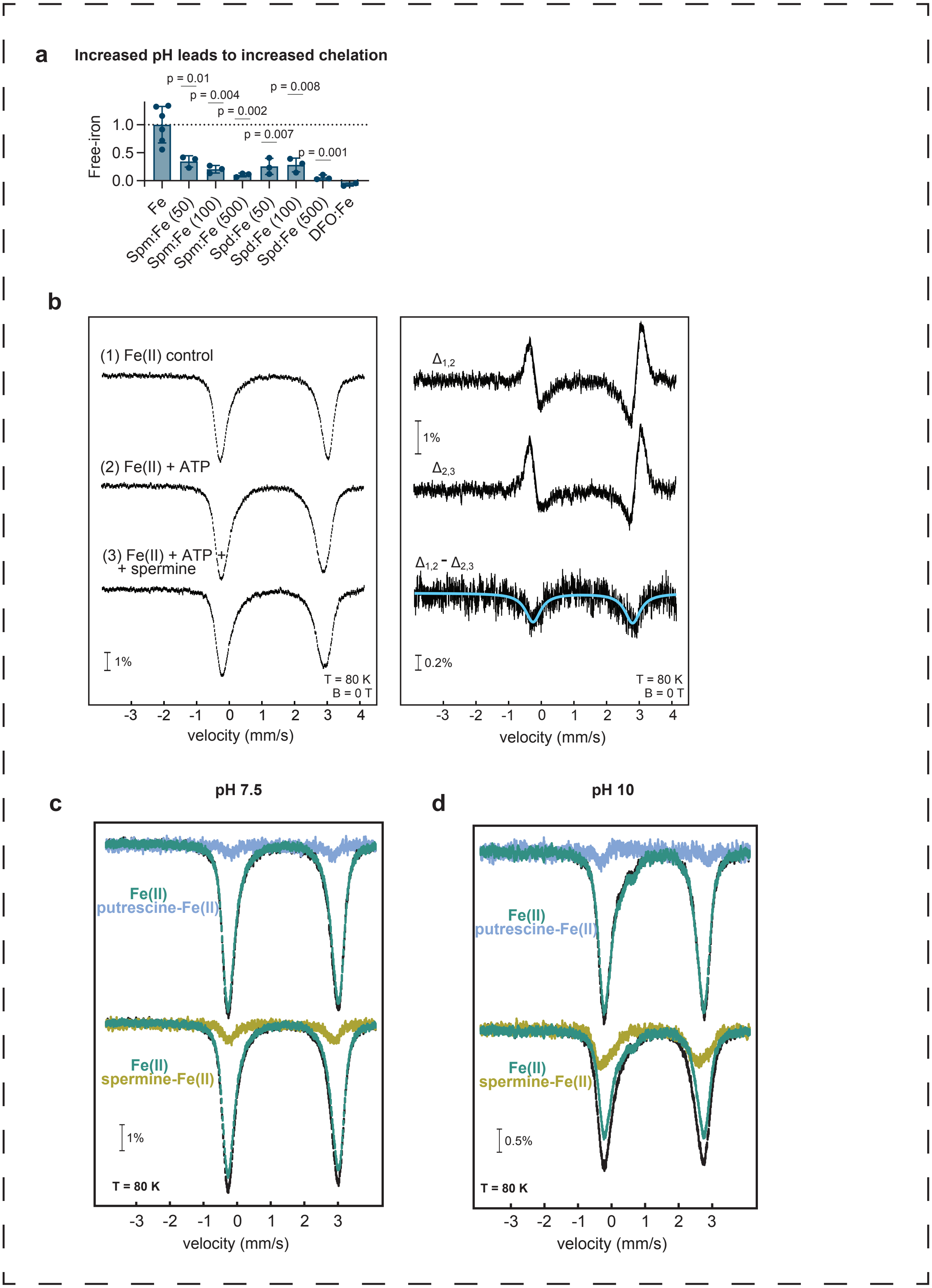
a
Increased pH leads to increased chelation
p = 0.01
p = 0.004
p = 0.002
p = 0.007
p = 0.008
p = 0.001
Free-iron
1.0
0.5
0.0
Fe
Spm:Fe (50)
Spm:Fe (100)
Spm:Fe (500)
Spd:Fe (50)
Spd:Fe (100)
Spd:Fe (500)
DFO:Fe
b
(1) Fe(II) control
(2) Fe(II) + ATP
(3) Fe(II) + ATP + + spermine
1%
T = 80 K
B = 0 T
velocity (mm/s)
$\Delta_{1,2}$
$\Delta_{2,3}$
$\Delta_{1,2} - \Delta_{2,3}$
1%
0.2%
T = 80 K
B = 0 T
velocity (mm/s)
c
pH 7.5
Fe(II)
putrescine-Fe(II)
Fe(II)
spermine-Fe(II)
1%
T = 80 K
velocity (mm/s)
d
pH 10
Fe(II)
putrescine-Fe(II)
Fe(II)
spermine-Fe(II)
0.5%
T = 80 K
velocity (mm/s)

**Figure S7. Polyamine biosynthesis genes are upregulated across human cancers and trientine shares structural similarity with higher-order polyamines, related to Figure 7**

(A-B) Relative mRNA expression of (A) ODC1 and (B) SRM in patient cancer tissues and patient-matched normal tissues. The plot is obtained from GEPIA2 (http://gepia2.cancer-pku.cn/#general), a web gene expression profiling tool that plots normalized mRNA-seq data from patient tumor tissues and normal tissues obtained from TCGA and GTEx.

(C) Chemical structure of trientine compared to spermidine and spermine.

Figure S7:

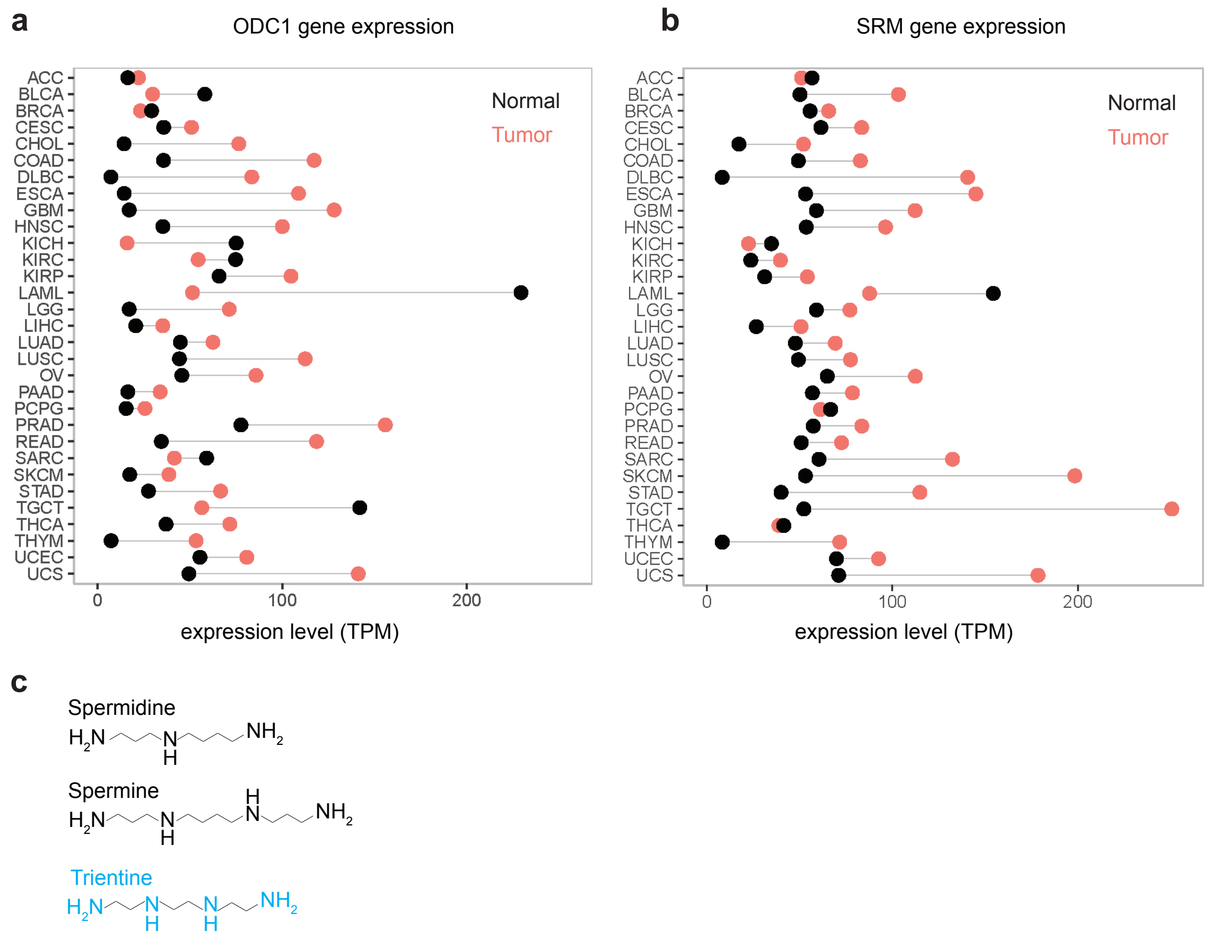
a
ODC1 gene expression
ACC
BLCA
BRCA
CESC
CHOL
COAD
DLBC
ESCA
GBM
HNSC
KICH
KIRC
KIRP
LAML
LGG
LIHC
LUAD
LUSC
OV
PAAD
PCPG
PRAD
READ
SARC
SKCM
STAD
TGCT
THCA
THYM
UCEC
UCS
Normal
Tumor
0
100
200
expression level (TPM)
b
SRM gene expression
Normal
Tumor
0
100
200
expression level (TPM)
c
Spermidine
Spermine
Trientine

**Supplemental table legends.**

**Table S1. Genome-wide CRISPR-Cas9 screen results under polyamine depletion, related to Figure 1.** Gene-level MAGeCK/RRA results from the K562 genome-wide CRISPR-Cas9 screen comparing DFMO-treated and vehicle-treated cells. The table includes negative-selection and positive-selection scores, p values, false discovery rates, ranks, number of contributing sgRNAs, and log2 fold changes used to identify genes selectively depleted or enriched under polyamine-depleted conditions.

**Table S2. C8-positive lipidomics dataset, related to Figures S2B-H, S3F-G, and S4A-E.** Lipidomics measurements from K562 cells treated with vehicle, DFO, sardomozide (SARD), or sardomozide plus DFO. The table includes metabolite/lipid annotations, compound IDs, mass-to-charge ratios (MZ), retention times (RT), HMDB identifiers where available, HMDB annotation specificity, and replicate abundance values for each treatment condition. These data were used for the lipid class and PUFA-containing lipid analyses shown in Figures S2B-H, S3F-G, and S4A-E.

**Table S3. Tumor versus normal expression of ODC1 and SRM across cancer types, related to Figure S7A-B.** Normalized mRNA expression (TPM) of ODC1 and SRM in tumor and matched normal tissues across TCGA cancer types, obtained from GEPIA2 (http://gepia2.cancer-pku.cn; TCGA and GTEx). For each cancer type the tumor-minus-normal difference is given for ODC1 and SRM, with the mean difference across cancer types and a paired two-tailed t-test (tumor vs normal) for each gene.

**Table S4. Plasmid and construct sequences used in this study, related to Methods Details.** Plasmid or construct sequence information for the CRISPR knockout and reporter reagents used in this study, including GPX4, SRM, SMS, and AAVS1-targeting constructs, as well as the genetically encoded iron and polyamine reporter constructs.

**Supplemental note 1**

We engineered constructs containing one, two, or three copies of the ferritin 5′ UTR iron-responsive element (IRE) upstream of an EBFP2 fluorescent reporter, under a doxycycline-inducible promoter, with an IRES-driven EGFP as an internal control. These constructs were stably integrated into K562 cells and assessed for (1) doxycycline-induced EBFP2 expression and (2) iron sensitivity, measured by changes in the EBFP2/EGFP ratio following treatment with the iron chelator DFO and rescue with exogenous iron. All constructs showed iron-dependent repression of EBFP2, with varying dynamic ranges.

- The 1 IRE construct showed a 58.1 % decrease in EBFP2/EGFP upon iron depletion ($0.42 \pm 0.22$), which was reversible with FAC ($0.91 \pm 0.24$).
- The 2 IREs construct showed a 46.4 % decrease in EBFP2/EGFP upon iron depletion ($0.54 \pm 0.28$), which was reversible with FAC ($1.02 \pm 0.27$).
- The 3 IREs construct showed a 43.7 % decrease in EBFP2/EGFP upon iron depletion ($0.56 \pm 0.22$), which was reversible with FAC ($0.92 \pm 0.20$).

We chose the 2 IREs construct because it offers a strong response to iron depletion (46.4% reduction) with full reversibility (EBFP2/EGFP restored to $1.02 \pm 0.27$ with FAC).
