## Supplementary material for "Polyamines buffer labile iron to suppress ferroptosis": Key Resource Table

### Key resources table

| REAGENT or RESOURCE | SOURCE | IDENTIFIER |
| --- | --- | --- |
| Antibodies | | |
| ACSL4 | Thermo Fisher | CAT#PA5-27137 |
| anti-Rabbit HRP Conjugated Secondary | Sigma Aldrich | CAT#A0545 |
| Beta-Actin | Abcam | CAT#ab20272 |
| EGFP | Abcam | CAT#ab6556 |
| FSP1 | Cell Signaling Technology | CAT#24972S |
| FTH | Abcam | CAT#ab75973 |
| GPX4 | Abcam | CAT#ab41787 |
| Hypusine | Millipore | CAT#ABS1064-I |
| SLC40A1 | Novus Biological | CAT#NBP1-21502 |
| SMS | Abcam | CAT#ab156879 |
| SRM | Proteintech | CAT#19858-1-AP |
| TFRC | Invitrogen | CAT#13-6800 |
| Bacterial and virus strains | | |
| CRISPR sgRNA library | This Paper | PMID#37218462 |
| Chemicals, peptides, and recombinant proteins | | |
| 0.1% (w/v) Sodium Dodecyl Sulfate; SDS | Bio-Rad | CAT#1610302 |
| 1,2-didodecanoyl-sn-glycero-3-phosphocholine | Avanti | CAT#850335 |
| 15N-Cysteine | MedChem Express | CAT#HY-Y03375 |
| 4xBolt Lithium Dodecyl Sulfate; LDS | Invitrogen | CAT#B0007 |
| A-1331852 | MedChem Express | CAT#HY-19741 |
| Benzonase nuclease | EMD Millipore | CAT#E1014 |
| Bovine Serum Albumin, BSA | Sigma Aldrich | CAT#A7906-100G |
| C11-BOPIDY | Thermo Fisher Scientific | CAT#D3861 |
| Fe Standard Curve; Cannabis Heavy Metals Environmental Calibration Standard | Agilent Technologies | CAT#5183-4688 |
| CM-H2DCFDA | Thermo Fisher Scientific | CAT#C6827 |
| DL-α-difluoromethylornithin; DFMO | Cayman | CAT#16889 |
| Deferoxamine Mesylate; DFO | Cayman | CAT#14595 |
| Dimethyl Sulfoxide, DMSO | Sigma Aldrich | CAT#276855 |
| Doxycycline, DOX | Sigma Aldrich | CAT#D9891-10G |
| Dithiothreitol; DTT | Thermo Fisher Scientific | CAT#R0861 |
| Ellman's Reagent | Thermo Fisher Scientific | CAT#22582 |
| ExTaq Polymerase | Takara Bio | CAT#RR001B |
| Ferric Ammonium Citrate | Sigma | CAT#F5879 |
| FerroOrange | Dojindo | CAT#F374 |
| Ferrous Ammonium Sulfate | Sigma Aldrich | CAT#215406 |
| Ferrostatin-1 | Cayman | CAT#17729 |
| Ferrozine | Sigma Aldrich | CAT#82950-1G |
| HALT Protease and Phosphatase Inhibitors | Thermo Fisher Scientific | CAT#78429 |
| iFSP1 | MedChem Express | CAT#HY-13607 |
| Iron (II) Chloride | Sigma Aldrich | CAT#220299 |
| Lipofectamine LTX | Invitrogen | CAT#15338-100 |
| Liprostatin-1 | MedChem Express | HY12726 |
| LysoTracker | Invitrogen | CAT#L12492 |
| ML162 | MedChem Express | HY-100002 |
| Nitric Acid, 67% Trace Metal Free | VWR | CAT#87003-226 |
| Non-Fat Dry Milk | BD Biosciences | CAT#232100 |
| Oleic Acid | Sigma Aldrich | CAT#O3008 |
| Palbociclib | MedChem Express | CAT#HY-50767 |
| (2,2,5,7,8-pentamethyl-6-chromanol); PMC | MedChem Express | CAT#HY-111024 |
| Polybrene | Millipore Sigma | TR1003G |
| ProteinaseK | Millipore Sigma | CAT#3115879001 |
| RhoNox-1 | This Paper | N/A |
| RhoNox-M | This Paper | N/A |
| RSL3 | MedChem Express | CAT#HY-100218A |
| Sardomozide Dihydrochloride | MedChem Express | CAT#HY-13746B |
| Sodium Chloride; NaCl | Invitrogen | CAT#AM9760G |
| soy L-α-phosphatidylcholine | PC, Avanti | CAT#441601 |
| Spermidine Trihydrochloride | Sigma Aldrich | CAT#S2501 |
| Spermine Tetrahydrochloride | Thermo Fisher Scientific | CAT#J63060.14 |
| STY-BOPIDY | Cayman | CAT#27089 |
| SuperSignal West Femto Maximum Sensitivity Substrate | Thermo Fisher Scientifc | CAT#34095 |
| Tris-Buffered Saline; TBST | Thermo Fisher Scientific | CAT#AAJ6076K3 |
| Terbium (Tb) Internal Standard | Agilent | CAT#51908590 |
| Tert-Butyl Hydroperoxide | Thermo Fisher Scientific | CAT#180342500 |
| Tween-20 | Thermo Fisher Scientific | CAT#BP337 |
| Warfarin | Sigma Aldrich | CAT#1719000 |
| Water, Trace-Metal Free | VWR | CAT#87003-236 |
| Xtremegene-9 Transfection Reagent | Sigma Aldrich | CAT#06365779001 |
| Z-VAD(OH)-FAK | Cayman | CAT#14467 |
| AMXT-1501 tetrahydrochloride | MedChemExpress | HY-124617A |
| GC7 | MedChemExpress | HY-108314A |
| Tetrathiomolybdate (TTM) | Sigma | 323446 |
| Glutathione ethyl ester | Cayman | 14953 |
| Critical commercial assays | | |
| Blood genomicPrep Mini Spin Kit | Cytiva | Cytivia; CAT#28904264 |
| PureLink RNA Mini Kit | Invitrogen | CAT#12183018A |
| QIAmp DNA Blood Maxiprep Kit | Qiagen | Qiagen; CAT#51192 |
| Qubit dsDNA HS Assay Kit | Thermo Fisher Scientific | CAT#Q32851 |
| Fluorescent Total Polyamine Assay Kit | Abcam | CAT#239728 |
| Deposited data | | |
| CRISPR screen data | This Paper | GEO accession GSE300179 |
| Lipidomics data | This Paper | 10.5281/zenodo.20706003 |
| Experimental models: Cell lines | | |
| HEK-293T | ATCC | CAT#CRL-3216 |
| HEL | Gift From Boston Children’s Hospital | N/A |
| K562 | ATCC | CAT#CCL-243 |
| MEL | Gift From Boston Children's Hospital | N/A |
| MRC-5 | Gift From Punam Bisht, Whitehead Institute | N/A |
| NOMO | Gift From Naama Kanarek, Boston Children’s Hospital | N/A |
| RPE1 | ATCC | CAT#CRL-4000 |
| U2OS | ATCC | CAT#HTB-96 |
| Experimental models: Organisms/strains | | |
| Female NSG mice | Jackson | 005557 |
| Oligonucleotides | | |
| Index Sequencing Primer | Illumina | 5’- TTTCAAGTTACGGTAAGCATATGATAGTCCATTTTAAAACATAATTTTAAAACTGCAAACTA CCCAAGAAA - 3’ |
| PCR Forward Primer | This Paper, IDT | 5’- AATGATACGGCGACCACCGAGATCTACACCCCACTGACGGGCACCGGA - 3’ |
| PCR Reverse Primer | This Paper, IDT | 5’- CAAGCAGAAGACGGCATACGAGATCnnnnnnTTTCTTGGGTAGTTTGCAGTTTT - 3’ |
| Read 1 Sequencing Primer | Illumina | 5’- GTTGATAACGGACTAGCCTTATTTAAACTTGCTATGCTGTTTCCAGCATAGCTCTTAAAC - 3’ |
| Recombinant DNA | | |
| pCMV-VSV-G | Addgene | Addgene; Plasmid#8454 |
| psPAX2 | Addgene | Addgene; Plasmid#12260 |
| sgRNA/Cas9 plasmid (CRISPR Knockout) | This Paper, Heather R. Keys, Whitehead Institute | N/A |
| Software and algorithms | | |
| MAGeCK v0.5.9.3 | MAGeCK SourceForge | RRID:SCR_025016 |
| STAR aligner (v2.7.1a) | STAR GitHub Repository | RRID:SCR_004463 |
| edgeR | Bioconductor | RRID:SCR_012802 |
| featureCounts 2.0.1 | N/A | RRID:SCR_012919 |
| UMICollapse v1.1.0 | GitHub | N/A |
| XenofilteR v1.6 | GitHub | RRID: SCR_026196 |
| GraphPad Prism | GraphPad Software | RRID:SCR_002798 |
| TraceFinder | Thermo Fisher Scientific | RRID:SCR_023045 |
| DESeq2 | Bioconductor | RRID:SCR_015687 |
| fastp v0.24.0 | fastp GitHub Repository | RRID:SCR_016962 |
| Progenesis QI | Nonlinear Dynamics | RRID:SCR_018923 |
| Compound Discoverer 3.1 | Thermo Fisher Scientific | Compound Discoverer v3.1 |
| Agilent MassHunter | Agilent Technologies | RRID:SCR_015040 |
| FlowJo (version 11) | Becton, Dickinson and Company (BD) | RRID:SCR_008520 |
| Other | | |
| Perfluoroalkoxy Tubes, metal free; PFA Tubes | Savillex | CAT#200-915-50 |
| 1% (v/v) NP-40 | Thermo Fisher Scientific | CAT#AAJ19628AP |
| 1% (w/v) Sodium Deoxycholate | Sigma Aldrich | CAT#D6750 |
| 6-Well Plates | Falcon | CAT#353046 |
| DMEM | Gibco | CAT#11965126 |
| Fetal Bovine Serum; FBS | Gibco | CAT#A56707-01; LOT#U2933097RP |
| HBSS | Gibco | CAT#14025092 |
| IMDM | Thermo Fisher Scientific | CAT#1244053 |
| Inactivated Fetal Bovine Serum; inactivated FBS | Gemini Bio | CAT#100-106; GeminiBio |
| Opti-MEM | Thermo Fisher Scientific | CAT#11058021; Reduced Serum CAT#31985-070 |
| PBS, pH 7.4 | Gibco | CAT#14190250 |
| Penicillin-Streptomycin | Gibco | CAT#10378016 |
| Puromycin Dihydrochloride (3 µg/mL) | Gibco | CAT#A1113802 |
| RIPA Lysis Buffer | This Paper | N/A |
| RPMI-1640 | Gibco | CAT#11875093 |
| T175cm^2^ Flasks | Thermo Fisher Scientific | CAT#1243054 |
| T225cm^2^ Flasks | Thermo Fisher Scientific | CAT#159934 |
| Tris-HCl, pH 7.5 | Invitrogen | CAT#15567027 |
